# Integration of mathematical modeling and economics approaches to evaluate strategies for control of *Salmonella* Dublin in a heifer-raising operation

**DOI:** 10.1101/2025.01.08.630693

**Authors:** Sebastian Llanos-Soto, Martin Wiedmann, Aaron Adalja, Christopher Henry, Paolo Moroni, Elisha Frye, Francisco A. Leal Yepes, Renata Ivanek

## Abstract

*Salmonella* Dublin infections in heifer-raising operations (HROs) cause animal health and economic losses for these operations and represent a pathogen source for dairy farms obtaining replacement heifers from HROs. To improve control of *S*. Dublin, we (i) developed a mathematical model of *S.* Dublin transmission on a HRO, (ii) evaluated the vaccine effectiveness and cleaning improvements for controlling the infection, and (iii) evaluated the influence of infection and control strategies on the HRO’s operating income. We developed a modified Susceptible-Infected-Recovered-Susceptible model of *S.* Dublin spread in a batch-stocking HRO post-introduction of an index case, with stochasticity introduced through Monte Carlo simulations. Epidemiological outcomes (*S.* Dublin-induced deaths and abortions during raising and *S.* Dublin carriers and asymptomatic infections among raised replacement heifers) and operating income per 100-head raised on a HRO over a 2-year simulation were compared between control scenarios. We validated our model against *S.* Dublin infection data in cattle. Partial rank correlation coefficient and classification trees were used to determine parameter influence on model outcomes. Our model predicts a median of 37 carriers and 92 asymptomatic infections among raised replacement heifers out of 2,330 heifers that departed the operation by the end of the 2-year simulation period, highlighting the critical role of HROs in spreading *S*. Dublin. Increasing barn floor cleaning frequency (to a maximum of 12x per day) meaningfully reduced the *S.* Dublin epidemiological outcomes and improved the HRO’s operating income. Depending on the cost of cleaning, the median operating income increased between 1.2% to 10.6% in the first year when cleaning 12x per day compared to baseline (cleaning 1x per week). In most cost scenarios, predictions do not support the use of vaccination, even when paired with stringent cleaning measures. The developed model is expected to aid efforts to control *S.* Dublin in HROs.

## Introduction

Cattle movement between heifer-raising operations (HROs) and dairy farms increases exposure opportunities and susceptibility to *Salmonella* infection due to stress associated with the transition between the two environments, dietary changes, and interaction with new animals [1]. Among *Salmonella* serotypes, *Salmonella* Dublin represents a major threat to HROs due to its ability to cause a long-term infection in calves post-recovery from clinical disease (i.e., carrier state; [2]), thus providing an opportunity for the infection to be introduced into the HRO (and eventually spread to other herds) unnoticedly. The importance of HROs in pathogen dissemination among dairy operations is particularly concerning, considering the widespread multi-drug resistance (MDR) among *S.* Dublin isolates in the US [3, 4, 5]. The MDR in *S.* Dublin leads to significant challenges in veterinary treatment and control [6] and serves as a source of resistance genes for other *Salmonella* serotypes and bacterial pathogens affecting dairy cattle [7]. Because *S*. Dublin can spread from dairy farms to HROs via calves and from HROs to dairy farms via pregnant replacement heifers, HROs are positioned as potential amplifiers and disseminators of this pathogen within the dairy farming system. To diminish the occurrence of infection in dairy herds, including MDR strains, it is critical to understand *S.* Dublin infection dynamics in HROs and determine the most effective (i.e., yielding the best outcome in a real-life scenario) approach to reducing transmission of *S.* Dublin in HROs.

By synthesizing previously collected data and information published in the literature, mathematical models can provide valuable insights into *S.* Dublin transmission dynamics and assist in identifying knowledge gaps regarding its microbiology and epidemiology in HROs. For instance, previous modeling studies have explored the importance of environmental contamination in the persistence of *Salmonella* spp. (including *S*. Dublin) in cattle [8, 9, 10, 11] and the role of clinically infected, long-term infected, and supershedder individuals in *S*. Dublin transmission dynamics [11, 12]. Mathematical models have also been used to assess *what-if* scenarios and determine the importance of mitigation and biosecurity measures to prevent *S.* Dublin introduction and dissemination in dairy herds [8, 11, 12, 13, 14, 15, 16]. Importantly, mathematical models can account for the variability in real-world processes and the uncertainty in our understanding of those processes by incorporating stochasticity in parameter values, thus providing more realistic and comprehensive predictions of disease dynamics [17]. Predictions from mathematical models can be used to understand the economic impacts of infectious disease and to identify the most financially appropriate mitigation strategy through economic analyses [18]. Findings obtained through the combined application of epidemiological and economic approaches are expected to assist farmers and veterinarians in controlling *S.* Dublin infections in HROs. To date, available modeling studies have provided valuable insights into understanding *Salmonella* epidemiology in dairy operations and its economic impacts. However, they have not investigated the epidemiology of *S.* Dublin in HROs.

To improve control of *S.* Dublin on HROs, in this study we combined mathematical modeling and economic analysis to (i) develop a mathematical model of *S*. Dublin transmission on a HRO, (ii) evaluate the vaccine effectiveness and cleaning improvements for controlling the infection, and (iii) evaluate the influence of infection and control strategies on the HRO’s operating income.

## Materials and methods

### Heifer-raising operation characteristics

In this study, the model system is a US HRO that raises replacement heifers after weaning. Calves enter the HRO at 90 days old and depart at 630 days of age (i.e., for a total raising period of 540 days), preceding their first calving (note that pre-weaned calves and calved heifers are not considered in the model). Heifers’ arrival and departure age were determined based on data from the USDA [19]. The model HRO represents the system present in 53.9% of HROs in the USA in 2011 (Fig 1, [19]). It considers cattle groups in three age categories in the barn, each one with a designated subindex to help describe model parameters: (i) weaned calves (*_c_*), (ii) growing heifers (*_g_*), and (iii) pregnant heifers (*_p_*). The model was implemented as ordinary differential equations (ODE) in R v. 4.2.0 [20] using the *deSolve* R package [21]. The model is comprised of epidemiological and economic modules.

**Fig 1.**
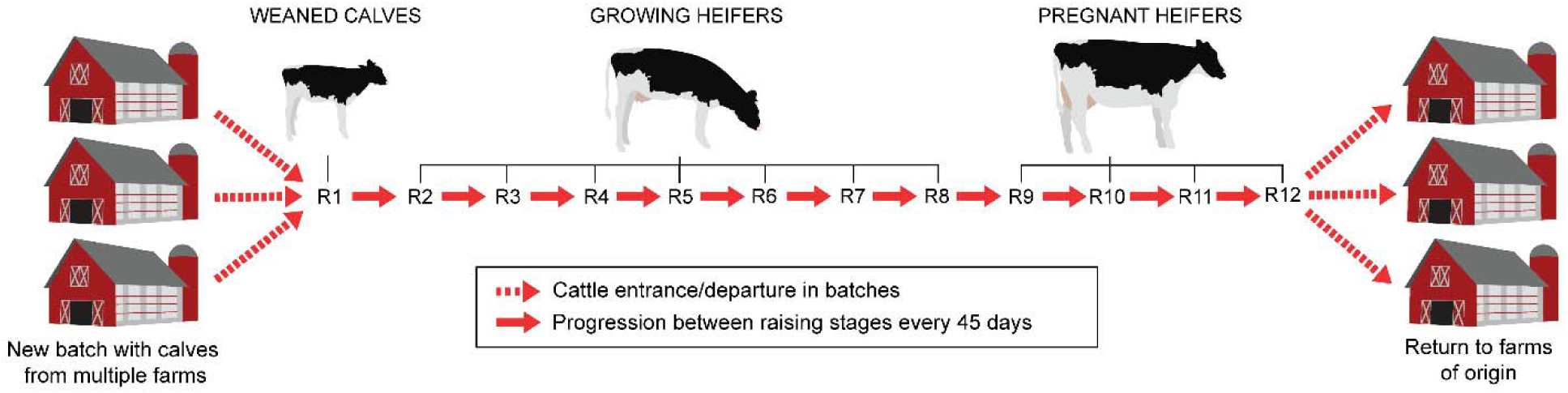
Heifer-raising operation workflow represented in the model. Calves enter the operation being 90 days old (weaned) and return to their farm of origin after 540 days (630 days of age and pregnant). During this period, individuals go through twelve raising stages: one as a weaned calf (raising stage R1; 90 to 135 days old), seven as a growing heifer (raising stages R2 to R8; 136 to 450 days old), and four as a pregnant heifer (raising stages R8 to R12; 451 to 630 days old). Individuals age from one stage to the next every 45 days and return to their farm of origin following raising stage R12.

### Epidemiological module

The epidemiological module is based on a modified Susceptible-Infected-Recovered-Susceptible (SIRS) compartmental model (Fig 2; model parameters are defined in Table 1). This module describes *S.* Dublin spread inside a free-stall barn in a HRO after introducing an asymptomatic infectious weaned calf (index case) at time *t*=0. The time unit of the ODE model is a day. Susceptible (*S*) individuals become infected through indirect transmission via the environment (*E*) contaminated with *S.* Dublin bacteria, at a rate β (per *Salmonella* cell/individual/day). The infected calves progress through three compartments defined based on clinical presentation, *S.* Dublin fecal shedding level, and the length of the infectious period: Asymptomatic (*A*), Clinically ill (*I*), and Carrier (*C*). Upon becoming infected, Susceptible cattle transition into *A* with probabilities 1-*u*_w_, 1-*u*_g_, or 1-*u_p_* depending on the cattle’s age or otherwise move into *I* (with probabilities *u*_w_, *u*_g_, or *u_p_*). The *A* compartment represents individuals experiencing an asymptomatic presentation of *S.* Dublin infection. Weaned calves and growing heifers in the *I* compartment can exhibit severe clinical signs and die at rates *d_w_* and *d_g_* (per day), respectively, due to bacteremia or systemic infection. While in *I* and *A*, infected cattle shed *S.* Dublin in feces at a constant level ε*_n_*(cell/individual/day) and λ*_n_* (cell/individual/day), respectively, where the subscript *n* corresponds to the raising stages within a HRO: weaned heifers (R1), growing heifers (R2 to R8), and pregnant heifers (R9 to R12).

**Fig 2.**
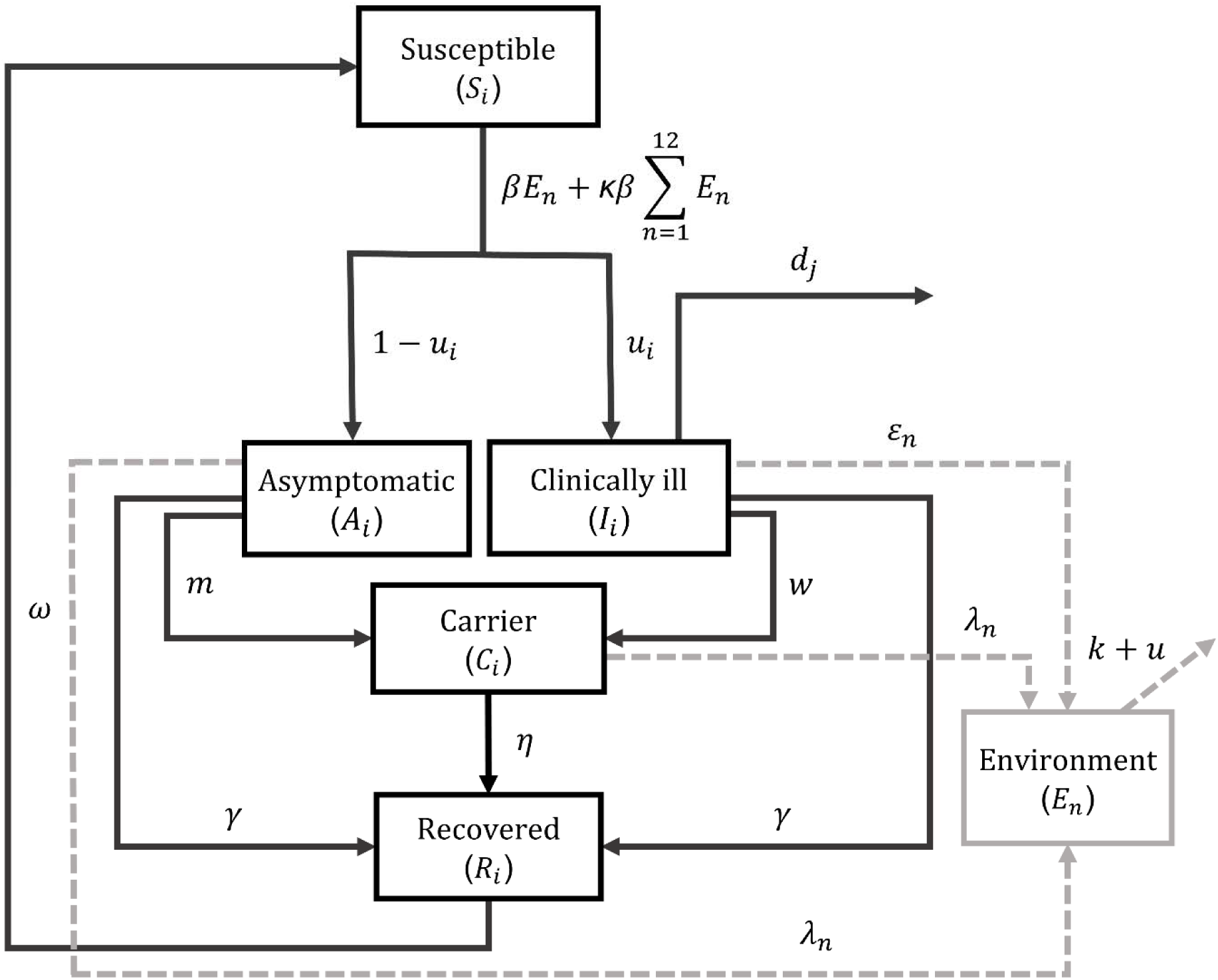
Transmission dynamics of *Salmonella* Dublin in a heifer-raising operation (HRO). Transmission dynamics model of *S.* Dublin infection in a HRO in a baseline scenario in which no mitigation strategy has been implemented. Solid arrows represent an individual animal’s movement among compartments and dashed arrows indicate *S.* Dublin shedding and the dynamics in the environment. A subscript *i* indicates that a parameter value changes according to age category; while a subscript *j* was added in parameter *d* to indicate that only weaned and growing heifers can die from *S.* Dublin infection. Subscript *n* indicates that the parameter varies in value according to the raising stage (R1 to R12). For compartment *E_n_*, subscript *n* represents the pen environment for each raising stage.

**Table 1.**
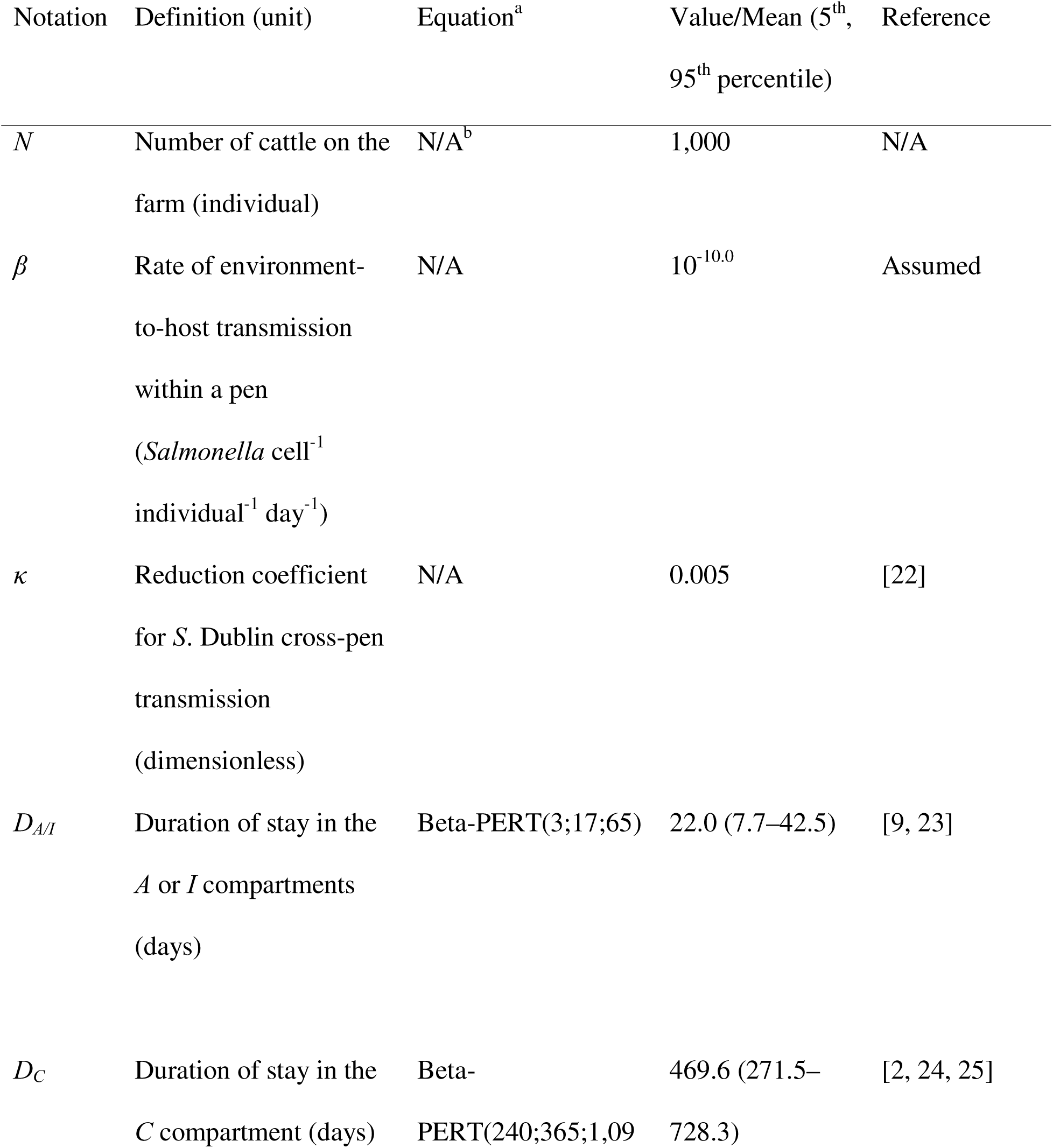
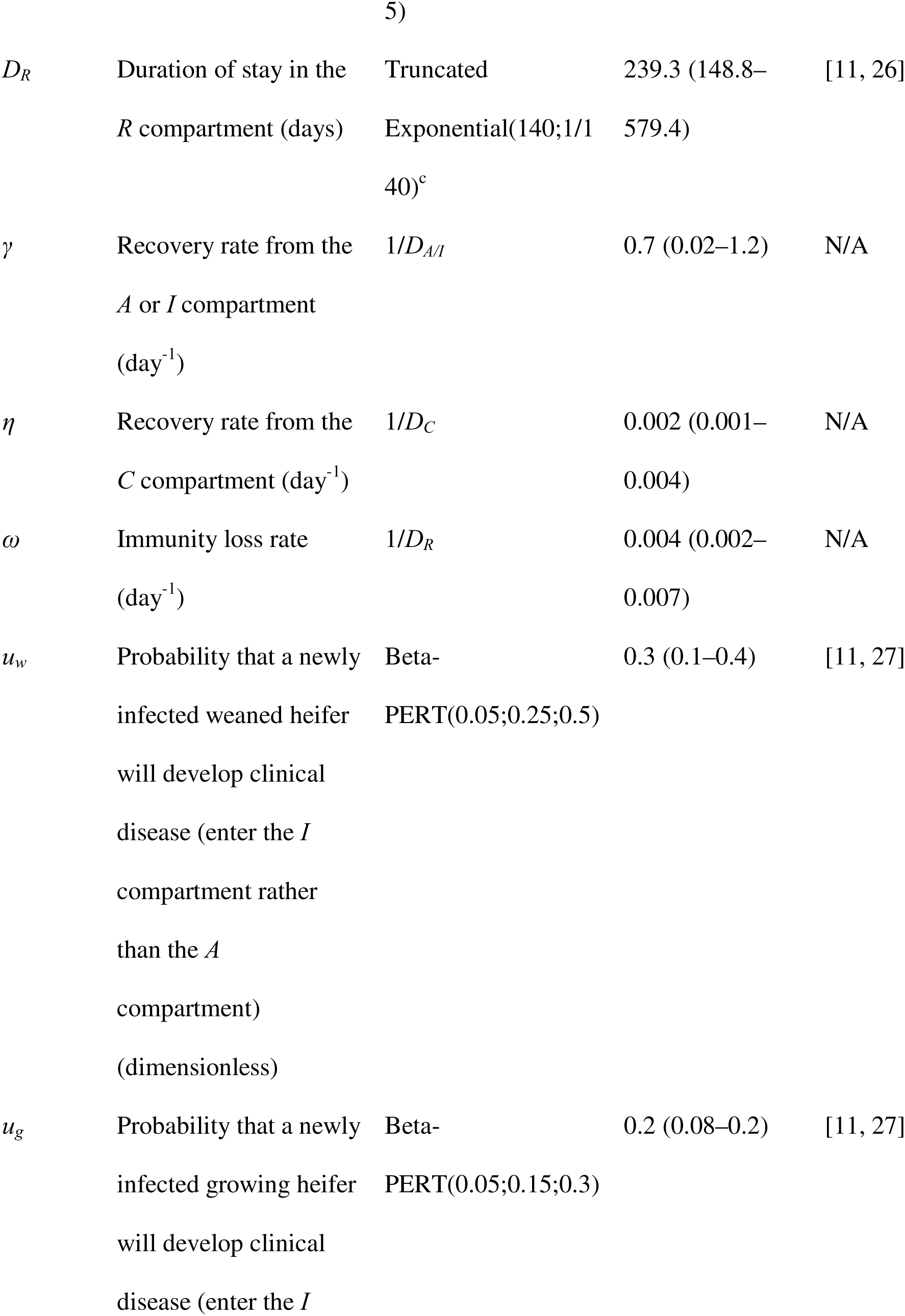
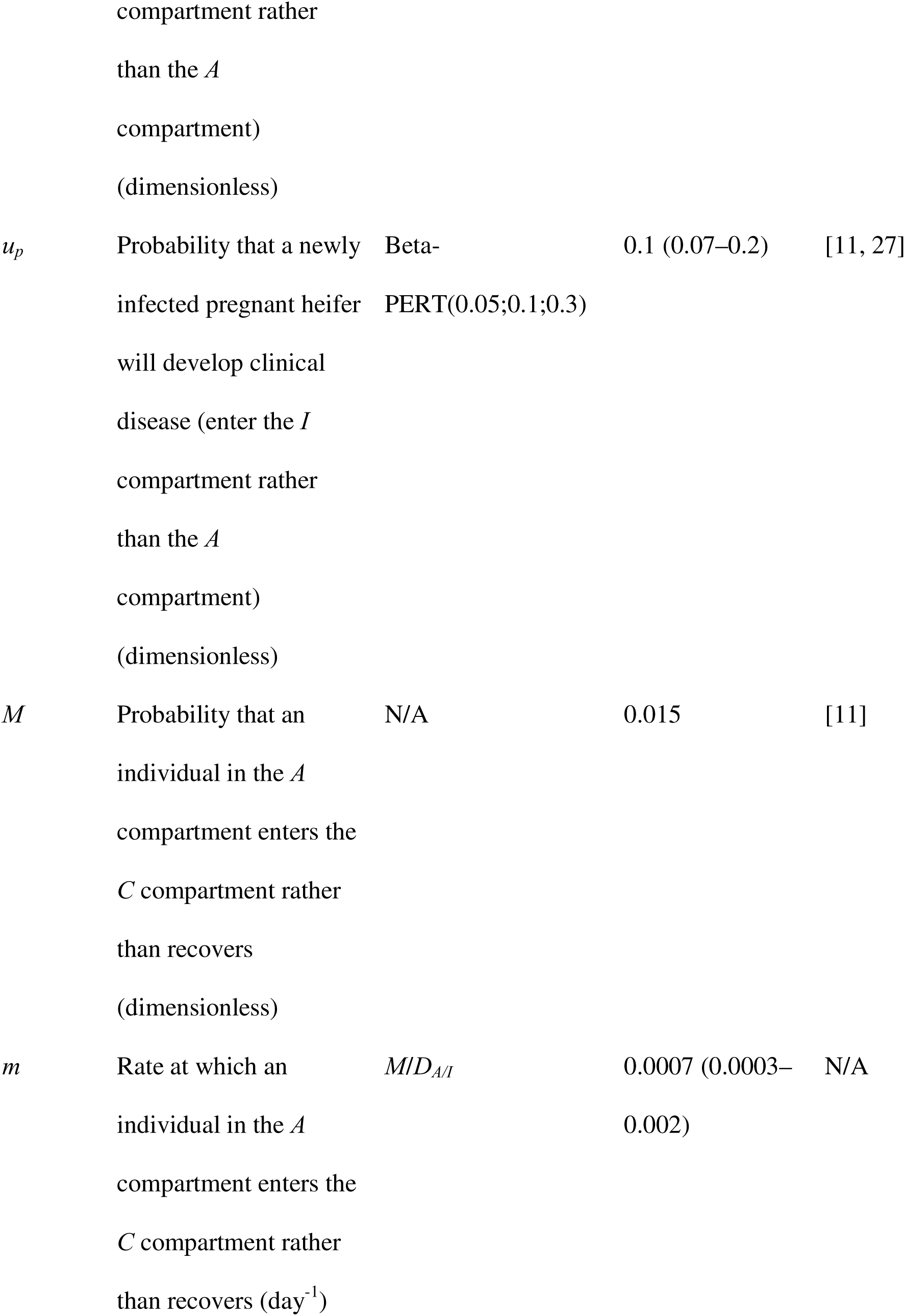
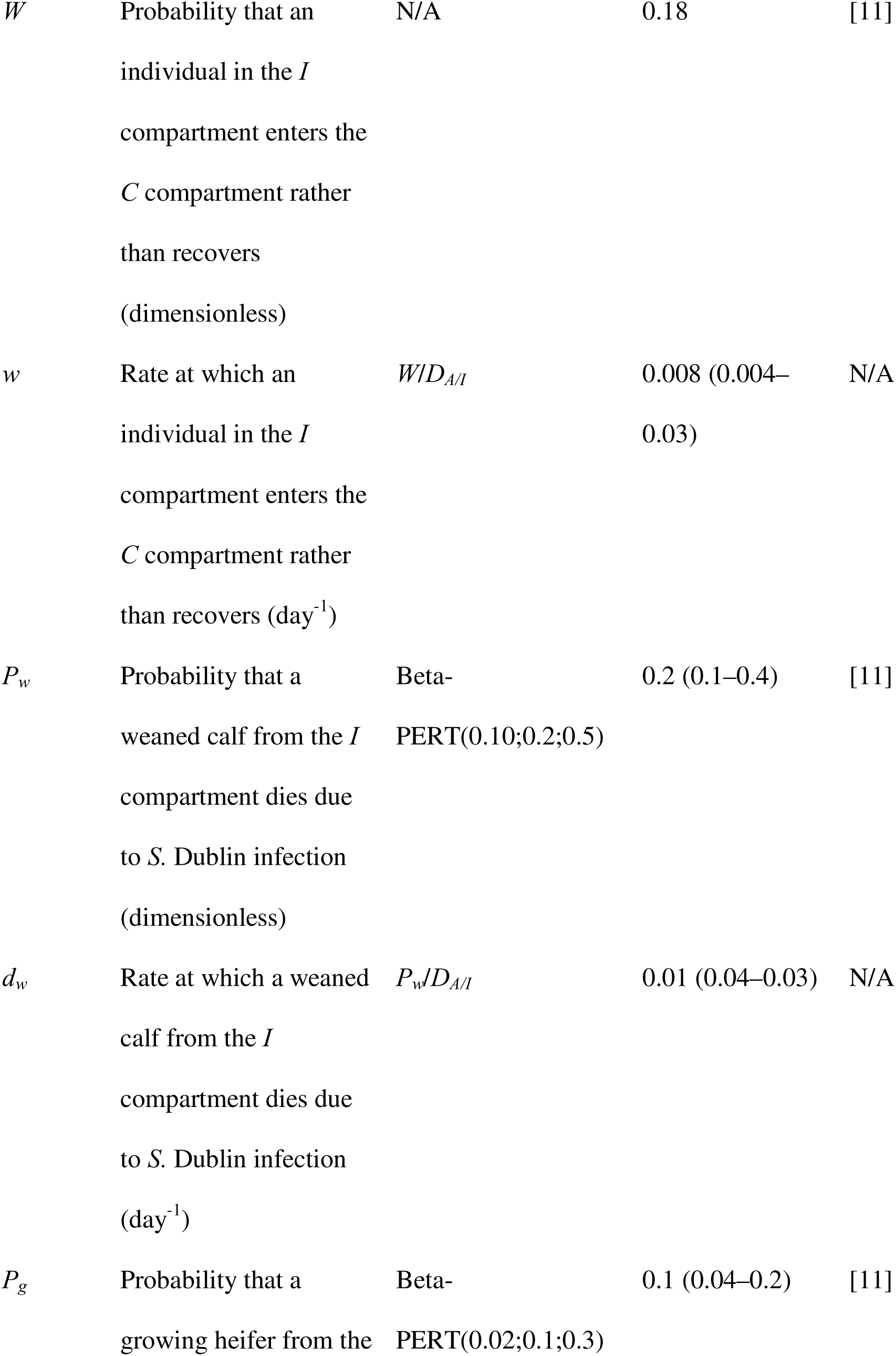
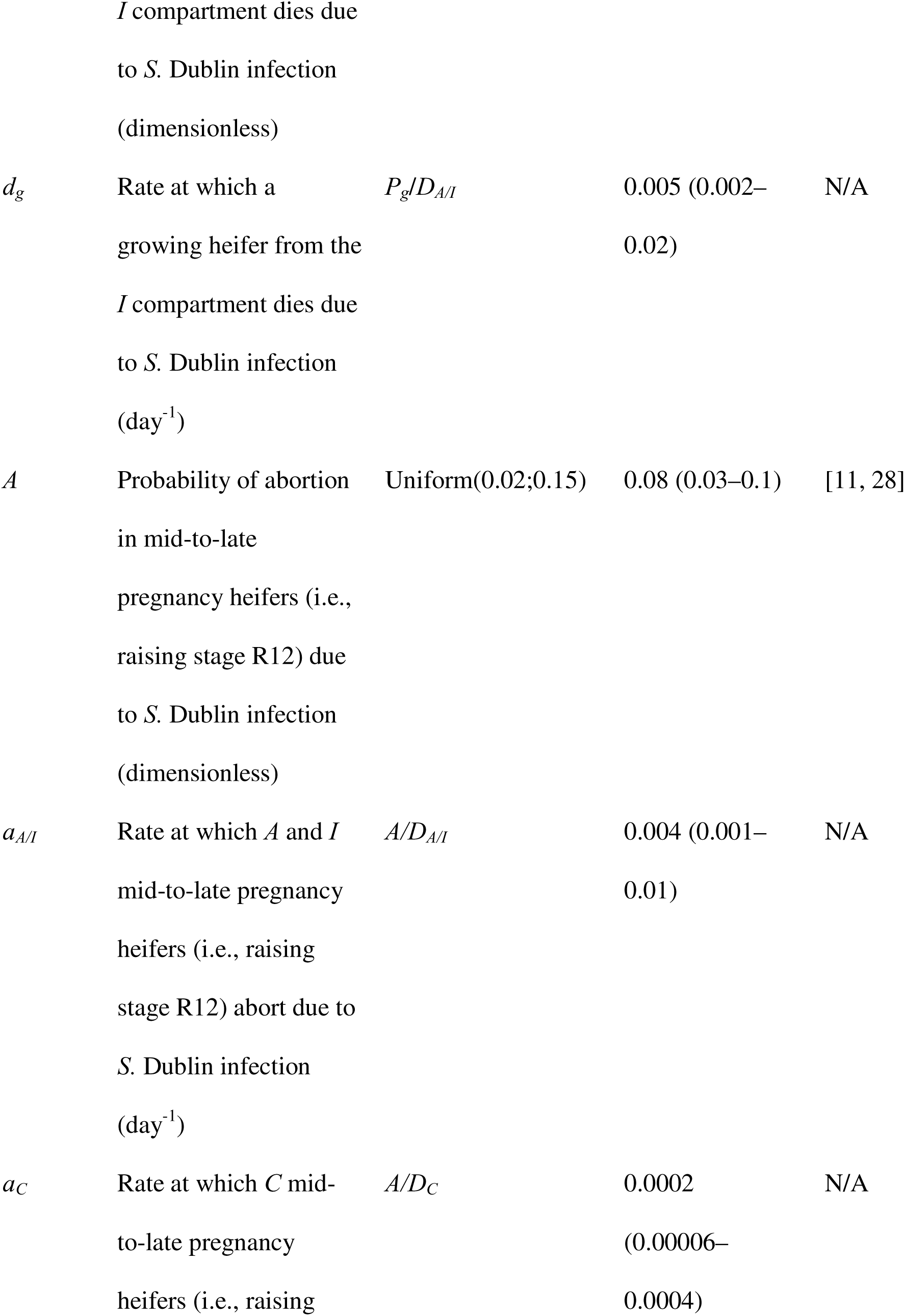
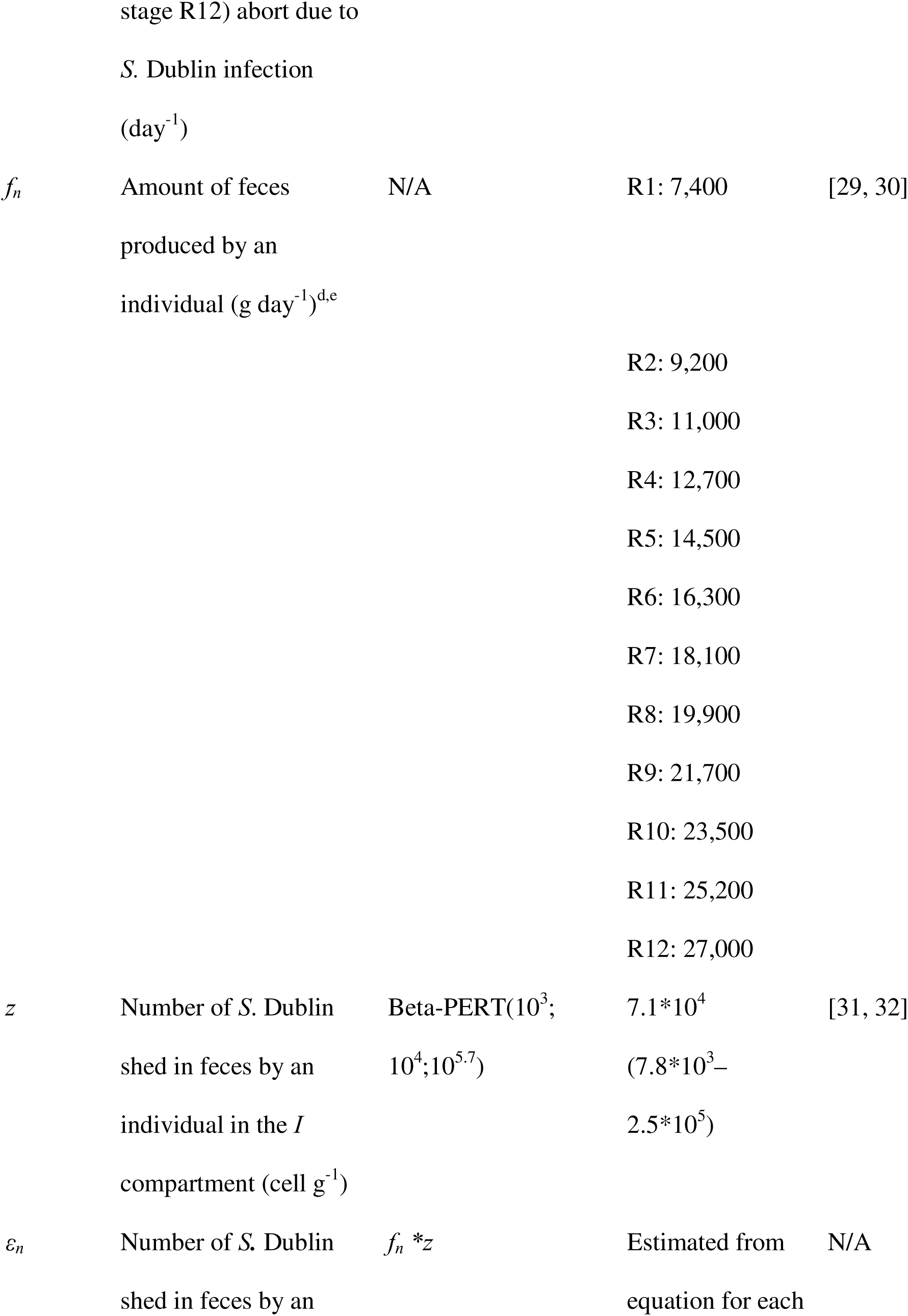
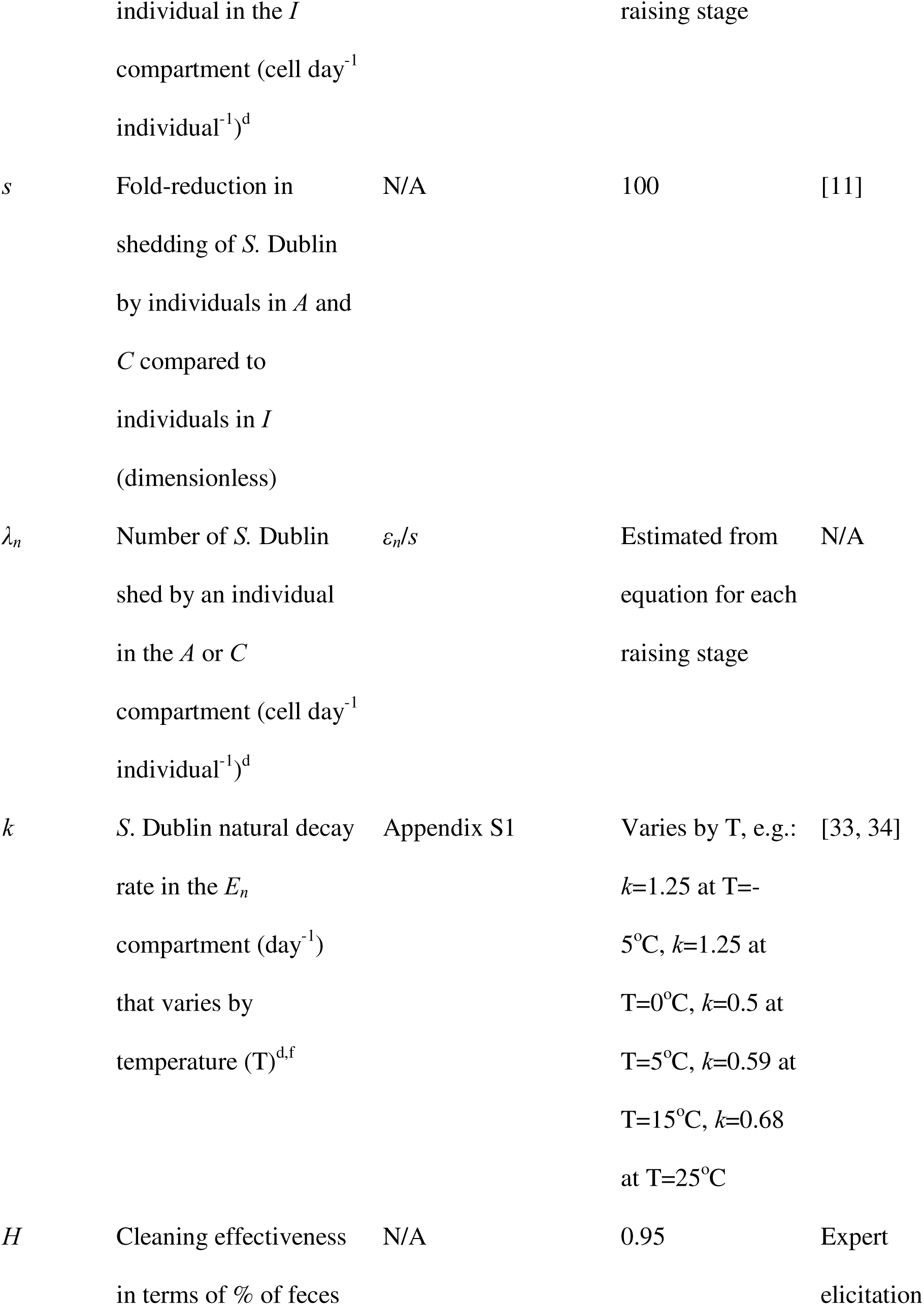
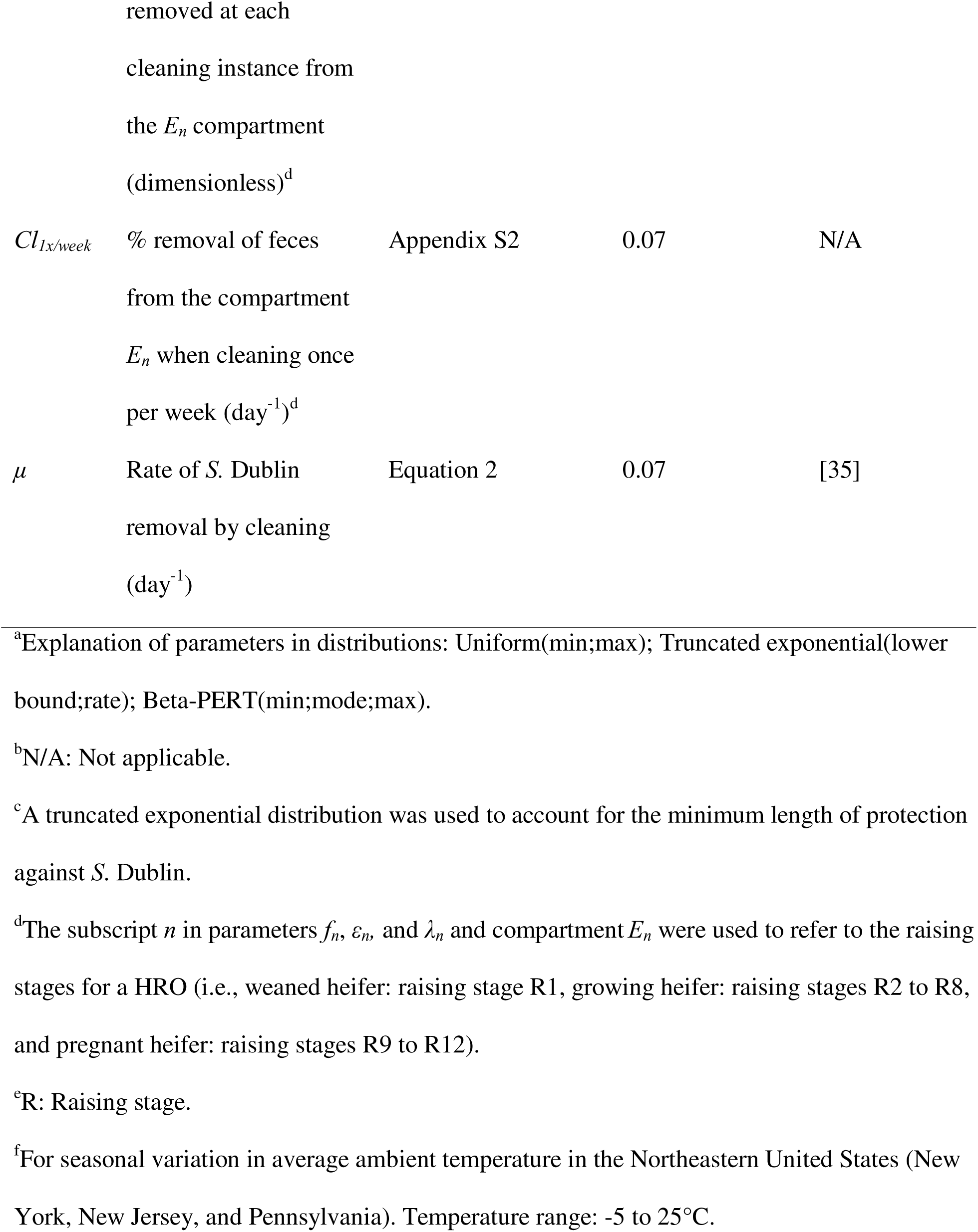
Details about the epidemiological module parameters, including equations and values. Parameter notations, definitions, and values for *Salmonella* Dublin infection in a heifer-raising operation in a baseline model (without control strategies).

Additionally, they either (i) recover from the infection, at a rate γ (per day) and enter the Recovered (*R*) compartment where they are temporarily immune to reinfection, or (ii) they develop a carrier status (compartment *C*) at rates *m* and *w* (per day) from the *A* and *I* compartments, respectively. Individuals in both *A* and *C* compartments shed *S.* Dublin at a rate λ*_n_,* which was considered s-fold (at the baseline s=100) lower than shedding by an *I* individual [11]. Eventually, individuals in *C* recover from infection, at a rate η (per day), and enter *R*. Infectious heifers can abort during late pregnancy (i.e., raising stage R12) with rates *a_A/I_* for *A* and *I*, and *a_C_* for individuals in *C*. An individual becomes once again susceptible to infection after immunity wanes at a rate ω (per day). The pathogen cells are removed from the environment based on a natural decay rate *k* (per day) dependent on the seasonally varying average ambient temperature and a barn cleaning rate μ (per day). Natural mortality among cattle was not included as a parameter as we focused on estimating an unbiased impact of *S*. Dublin on HROs (Fig S1 indicates that including or excluding natural mortality did not influence disease dynamics in the model).

The model accounted for animal aging by dividing each animal compartment into twelve raising stages: i) one for weaned calves (raising stage R1), ii) seven for growing heifers (raising stages R2 to R8), and iii) four for pregnant heifers (raising stages R9 to R12; Fig 1). Accordingly, the environmental compartment *E* was divided into twelve distinct sections (*E_n_*), where the subscript *n* corresponds to one of the twelve raising stages in an HRO. Each section represents the immediate environment specific to its corresponding raising stage (pen). The model also accounted for the spread of *S*. Dublin across pens, albeit at a slower rate (κ) than within pens. Individuals progress from one stage to the next every 45 days, with individuals departing from the HRO being replaced by a new batch of fully susceptible calves (batch size corresponding to one-twelfth of the initial herd size *N*). The subscript *n* (with values 1–12) was also used in model parameters to refer to one of the twelve stages during heifer raising at a HRO.

The initial conditions for the model considered *N*-1 susceptible individuals divided equally across raising stages (i.e., R1: (*N*/12)-1 and R2 to R12: *N*/12 individuals in each stage). After the introduction of an index case in raising stage R1, no additional infectious individuals are imported into the herd. The outcomes of interest predicted in the epidemiological module were (i) across all iterations, the probability of an outbreak (defined as the proportion of model iterations with two or more *S.* Dublin clinical cases) and (ii) at the individual iteration level, *S.* Dublin-induced deaths and abortions during raising, and *S*. Dublin carriers and asymptomatic infections among replacement heifers departing the HRO during the first and second year of the iteration, under the presence or absence of cleaning improvements and/or vaccination. Model predictions obtained in years beyond the second, representing infection endemic persistence in the HRO, were only minimally different and excluded from this assessment.

## Infection control strategies

### Vaccination

When vaccination is implemented in the model, all cattle in the herd are assumed to be fully vaccinated and the protective vaccine effectiveness (VE) is observed in all new arrivals from the moment of application of the first vaccine dose (i.e., the start of the simulation). Vaccination provides partial protection, meaning that vaccinated individuals can still experience the consequences of infection (i.e., clinical disease, death, carrier status, and abortion; [36–43]. We defined VE as 1 minus the relative risk in vaccinated animals versus unvaccinated animals (i.e., VE = 1 – risk ratio), with VE values ranging from 0 to 1 (0=no effectiveness). This approach to assessing vaccination was based on the theoretical study by Lu et al. [15]. Vaccination against *S.* Dublin has been shown to protect vaccinees through a proportional reduction in the probability of death in clinically ill calves ([42]; Table 2, Fig 3). The protection provided by vaccination was reported for calves (i.e., 2 to 5 weeks old), but due to the absence of information, here we extrapolated its effect to older cattle (i.e., weaned calves and growing heifers in the model), and then evaluated the impact of this assumption in sensitivity analysis. Some other effects, such as limiting the development of clinical symptoms, the carrier state in infected cattle, and fecal shedding, have been observed with experimental vaccines, but no similar effects have been reported for vaccines commercially available for use in cattle ([36–51], Fig 3). In the vaccination scenario, both weaned and growing heifers are vaccinated at the beginning of the simulation, with vaccines also being administered to new batches of weaned calves upon arrival to HRO. Growing heifers are revaccinated a year after their initial immunization as weaned heifers. Protection in vaccinated calves persists for their entire time in the HRO. The vaccine effectiveness against *S.* Dublin was represented in the model as a value reducing the probability of death after clinical disease (*d_j_*) as follows:

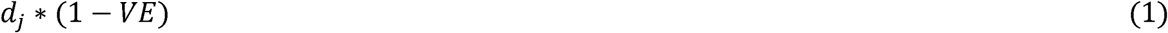

Where the subscript “*j*” refers to either weaned calves (*w*) or growing (*g*) heifers.

**Fig 3.**
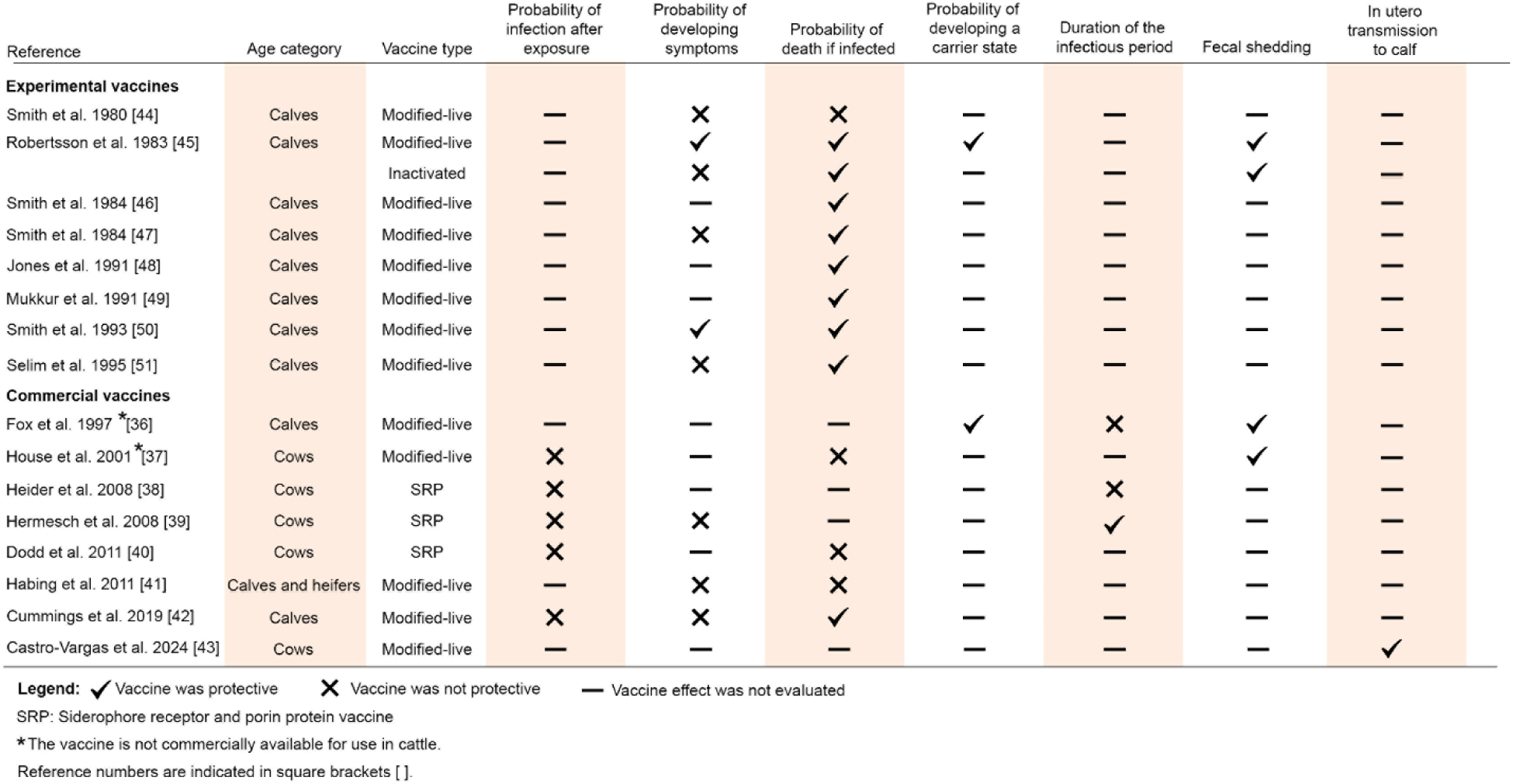
Experimental and commercial vaccine effects against *Salmonella* infection in cattle assessed and reported in the literature.

**Table 2.** Parameter notations, definitions, and values related to cleaning improvements and vaccination as control strategies against *Salmonella* Dublin spread in a heifer-raising operation.

| Notation | Definition (unit) | Equation | Mean (5<sup>th</sup>,<br>95<sup>th</sup><br>percentile) | Reference |
| --- | --- | --- | --- | --- |
| ***Vaccination*** |  |  |  |  |
| *VE* | Vaccine effectiveness in reducing the probability of death after clinical disease (dimensionless) | $VE=1-\text{Beta-PERT}(0.05;0.2;0.8)^a$ | 0.72 (0.48–0.91) | [42] |
| ***Improved cleaning*** |  |  |  |  |
| *Cl<sub>3x/week</sub>* | Average % of feces removed daily from the Environment compartment after cleaning three times per week (day<sup>-1</sup>)<sup>b</sup> | Appendix S2 | 0.20 | N/A<sup>b</sup> |
| *Cl<sub>5x/week</sub>* | Average % of feces removed daily from the Environment compartment after cleaning five times per week (day<sup>-1</sup>) | Appendix S2 | 0.33 | N/A |
| *Cl<sub>7x/week</sub>* | Average % of feces removed daily from the Environment compartment after cleaning seven times per week (day<sup>-1</sup>) | Appendix S2 | 0.47 | N/A |
| *Cl<sub>2x/day</sub>* | Average % of feces removed daily | Appendix | 0.74 | N/A |
|  | from the Environment compartment | S2 |  |  |
| | after cleaning two times per day ($\text{day}^{-1}$) | | | |
| $Cl_{4x/day}$ | Average % of feces removed daily | Appendix | 0.87 | N/A |
|  | from the Environment compartment | S2 |  |  |
| | after cleaning four times per day ($\text{day}^{-1}$) | | | |
| $Cl_{6x/day}$ | Average % of feces removed daily | Appendix | 0.91 | N/A |
|  | from the Environment compartment | S2 |  |  |
| | after cleaning six times per day ($\text{day}^{-1}$) | | | |
| $Cl_{8x/day}$ | Average % of feces removed daily | Appendix | 0.93 | N/A |
|  | from the Environment compartment | S2 |  |  |
| | after cleaning eight times per day ($\text{day}^{-1}$) | | | |
| $Cl_{12x/day}$ | Average % of feces removed daily | Appendix | 0.96 | N/A |
|  | from the Environment compartment | S2 |  |  |
| | after cleaning twelve times per day ($\text{day}^{-1}$) | | | |

<sup>a</sup>Explanation of parameters in distribution: Beta-PERT(min;mode;max). Parameter *VE* can only
vary from 0 to 1. In the baseline model, *VE*=0.
<sup>b</sup>N/A: Not applicable.

### Improved cleaning

The cleaning effectiveness per cleaning instance (*H*), understood as the proportion of feces that is removed from the barn every time the barn is scraped was determined based on expert elicitation from two veterinarians with more than 10 years of experience in dairy production medicine and bovine veterinary practice, one academic in dairy sciences, and one dairy management professional. Experts were asked about the perceived percentage of feces removed from the barn’s floor after one cleaning instance (i.e., the value of *H*) on a HRO or dairy farm they are familiar with, to which, on average, they responded that about 95% of feces (range: 90% to 99%) are removed. It is important to note that *H* does not describe the effectiveness of barn cleaning each day, as cattle will continue to produce feces immediately after every cleaning instance. The overall proportion of feces removed each day from the barn floor (referred to as the daily cleaning effectiveness, *CL_freq_*) was approximated by averaging the maximum (present just before cleaning) and minimum (present just after cleaning) amount of feces on the barn floor and bedding area (i.e., cow mattress) during the day for a particular cleaning frequency (calculation modified from Gautam et al. [35]). The calculation considered the amount of feces produced by an individual, the number of individuals in a given age category in the herd, and the cleaning frequency per day (Appendix S2). Then, the cleaning effectiveness *Cl_1/week_* represents the proportion of feces removed from compartment E when cleaning once per week (day^-1^).

Cleaning practices were assessed in eight scenarios (Table 2). In three of these scenarios, feces on the barn floor were cleaned multiple times per week: three times (3x) per week (*Cl_3x/week_*), 5x per week (*Cl_5x/week_*), and 7x per week (*Cl_7x/week_*). The remaining five scenarios considered cleaning multiple times per day: 2x per day (*Cl_2x/day_*), 4x per day (*Cl_4x/day_*), 6x per day (*Cl_6x/day_*), 8x per day (*Cl_8x/day_*), and 12x per day (*Cl_12x/day_*). The per week cleaning frequencies are achievable using a skid steer, while more frequent cleaning, as assessed in the study, would represent the use of an alley scraper. An additional scenario was included to assess *S.* Dublin dynamics in a bedded pack barn in which bedding is applied 1x per day and then completely replaced every 4 weeks. We assumed that bedding addition covers 95% of the manure present in the pen (i.e., equivalent to 95% cleaning effectiveness). The daily cleaning effectiveness ^(^*CL_freq_*), was converted into the rate μ (day^-1^) at which *S.* Dublin is removed every day from the barn environment using the following formula described in Gautam et al. [35]:

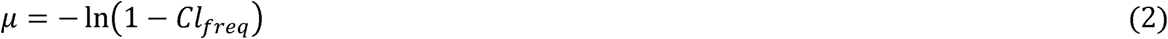

Where subscript “*freq*” refers to cleaning frequencies between 1x/week to 12x/day. When cleaning was performed on a weekly basis, the rate for cleaning once per day was adjusted by multiplying by 1/7, 3/7, or 5/7 for cleaning 1x, 3x, or 5x per week, respectively.

## Economic module

### HRO’s operating costs, income, and operating income

The economic module was based on partial budgeting. It takes the predictions from the epidemiological module to estimate the farm operating income, calculated as the difference between the income from raising heifers and the operating costs for running a HRO. The income encompassed the earnings from pregnant heifers returning to their farms of origin (*TIn)* and from the sale of apparently healthy open heifers for slaughter (i.e., heifers that aborted during mid-to-late pregnancy and prior to their departure from the HRO; *AIn*; Fig 4, Table 3). The number of open heifers sold was estimated as the cumulative number of *S*. Dublin-induced abortions by the end of the 2-year simulation period.

**Fig 4.**
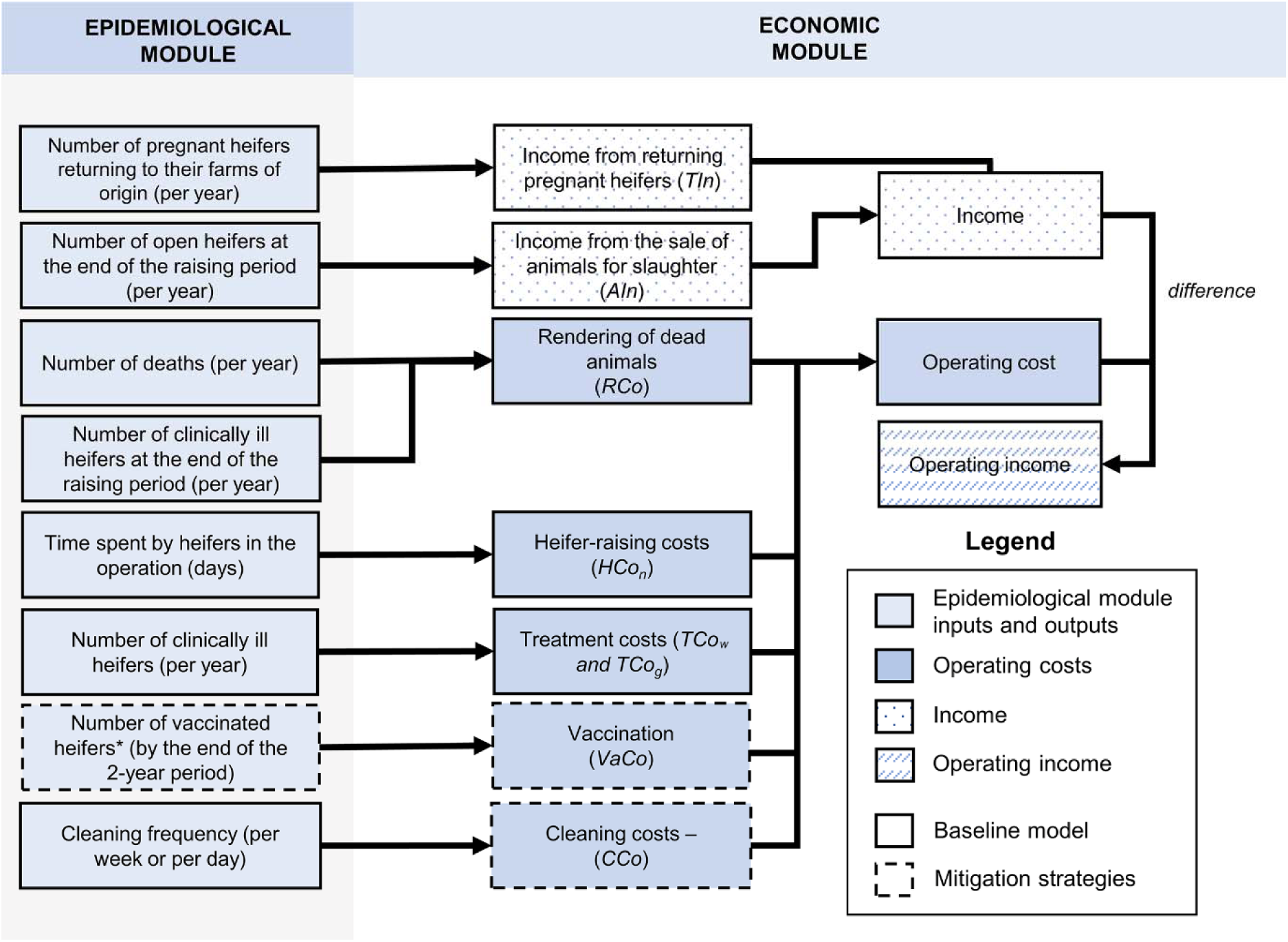
Components of the epidemiological and economic modules determine the operating cost, income, and operating income of a HRO. Inputs and outputs from the epidemiological module are used to calculate the operating cost, income, and operating income under a baseline “do-nothing” scenario (baseline level of cleaning (1x per week) and no vaccination) and scenarios with the implementation of cleaning improvements and/or vaccination.

**Table 3.**
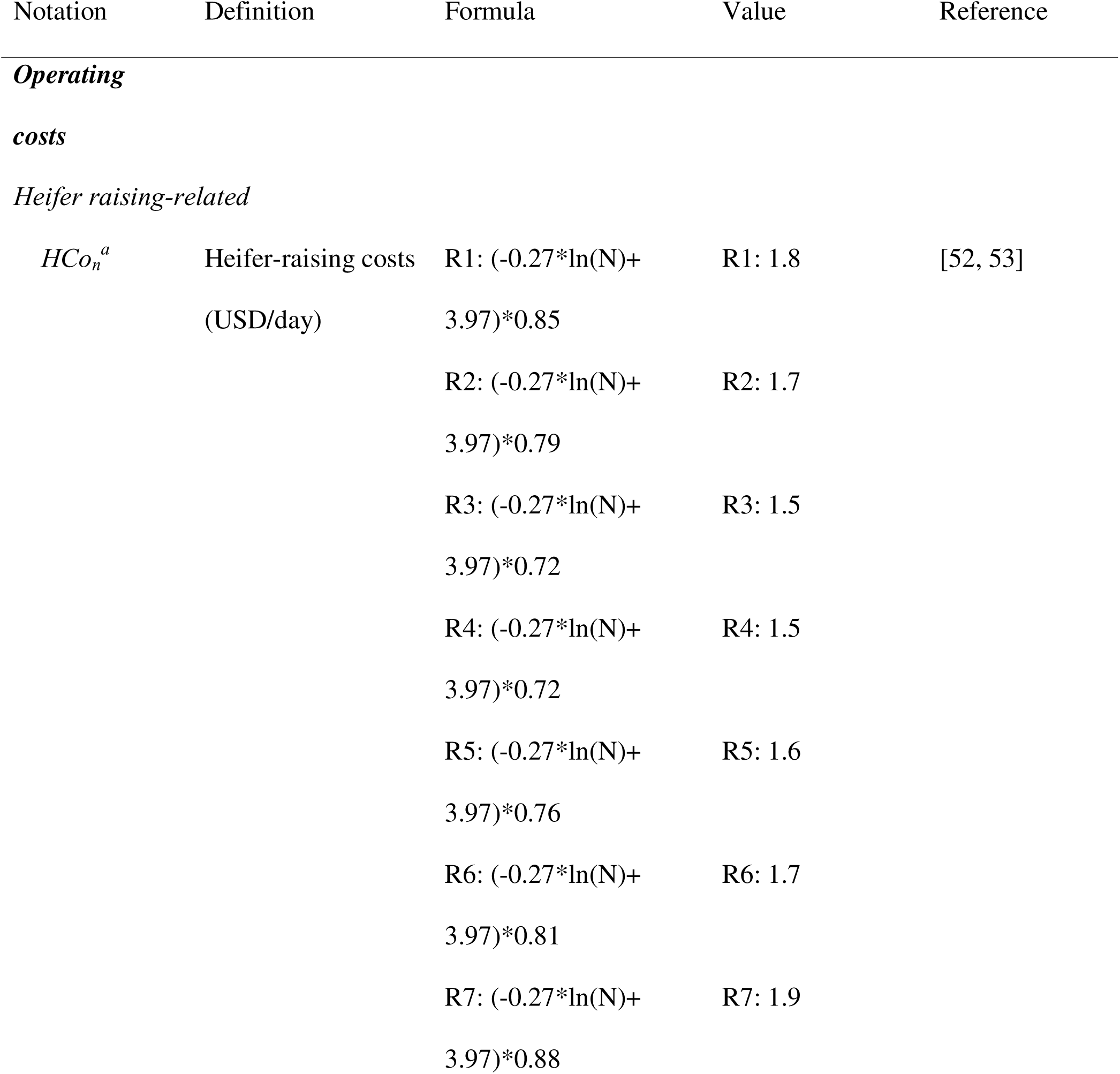
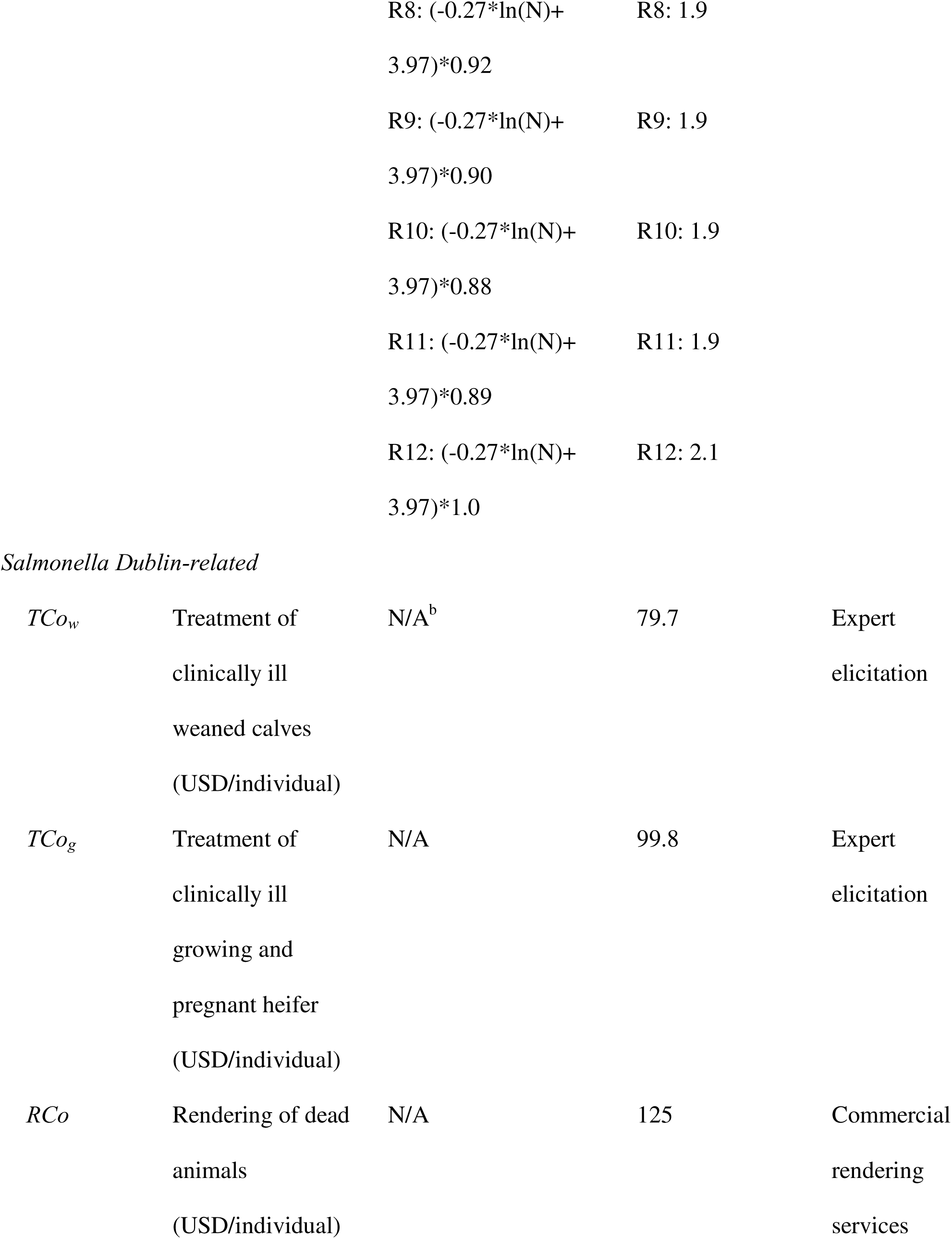
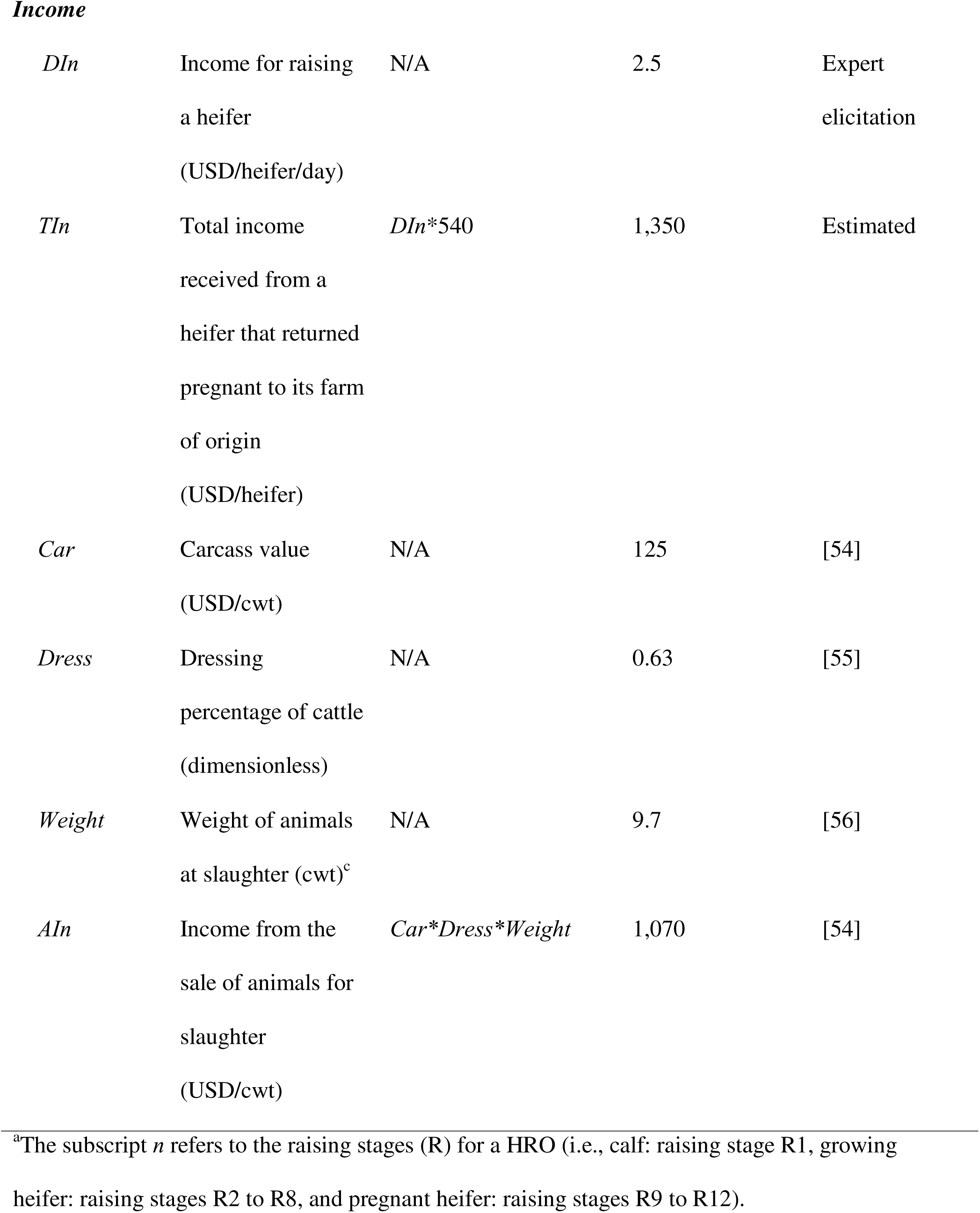
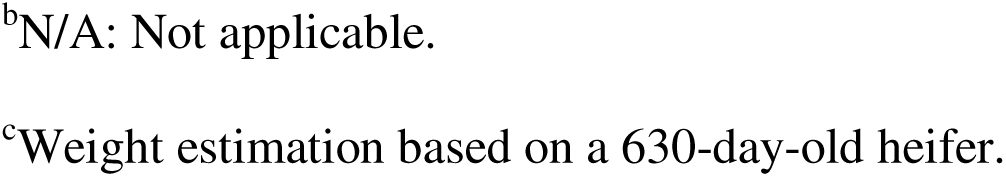
Details about the economic module parameters used to calculate operating costs and income when no mitigation strategies have been implemented (baseline). Operating costs and income are in USD. The number of cattle on the farm (*N*) is 1,000 for the baseline scenario.

| Notation | Definition | Formula | Value | Reference |
| --- | --- | --- | --- | --- |
| *Operating costs* |  |  |  |  |
| *Heifer raising-related* |  |  |  |  |
| $HCo_n^a$ | Heifer-raising costs<br>(USD/day) | R1: $(-0.27 \cdot \ln(N) + 3.97) \cdot 0.85$ | R1: 1.8 | [52, 53] |
| | | R2: $(-0.27 \cdot \ln(N) + 3.97) \cdot 0.79$ | R2: 1.7 | |
| | | R3: $(-0.27 \cdot \ln(N) + 3.97) \cdot 0.72$ | R3: 1.5 | |
| | | R4: $(-0.27 \cdot \ln(N) + 3.97) \cdot 0.72$ | R4: 1.5 | |
| | | R5: $(-0.27 \cdot \ln(N) + 3.97) \cdot 0.76$ | R5: 1.6 | |
| | | R6: $(-0.27 \cdot \ln(N) + 3.97) \cdot 0.81$ | R6: 1.7 | |
| | | R7: $(-0.27 \cdot \ln(N) + 3.97) \cdot 0.88$ | R7: 1.9 | |

|  |  |
| --- | --- |
| R8: $(-0.27 \cdot \ln(N) + 3.97) \cdot 0.92$ | R8: 1.9 |
| R9: $(-0.27 \cdot \ln(N) + 3.97) \cdot 0.90$ | R9: 1.9 |
| R10: $(-0.27 \cdot \ln(N) + 3.97) \cdot 0.88$ | R10: 1.9 |
| R11: $(-0.27 \cdot \ln(N) + 3.97) \cdot 0.89$ | R11: 1.9 |
| R12: $(-0.27 \cdot \ln(N) + 3.97) \cdot 1.0$ | R12: 2.1 |
| $TCO_w$ | Treatment of clinically ill weaned calves (USD/individual) | N/A<sup>b</sup> | 79.7 | Expert elicitation |
| $TCO_g$ | Treatment of clinically ill growing and pregnant heifer (USD/individual) | N/A | 99.8 | Expert elicitation |
| $RCo$ | Rendering of dead animals (USD/individual) | N/A | 125 | Commercial rendering services |
| *Income* |  |  |  |  |
| *DIn* | Income for raising<br>a heifer<br>(USD/heifer/day) | N/A | 2.5 | Expert<br>elicitation |
| *TIn* | Total income<br>received from a<br>heifer that returned<br>pregnant to its farm<br>of origin<br>(USD/heifer) | $DIn*540$ | 1,350 | Estimated |
| *Car* | Carcass value<br>(USD/cwt) | N/A | 125 | [54] |
| *Dress* | Dressing<br>percentage of cattle<br>(dimensionless) | N/A | 0.63 | [55] |
| *Weight* | Weight of animals<br>at slaughter (cwt)<sup>c</sup> | N/A | 9.7 | [56] |
| *AIN* | Income from the<br>sale of animals for<br>slaughter<br>(USD/cwt) | $Car*Dress*Weight$ | 1,070 | [54] |

<sup>a</sup>The subscript *n* refers to the raising stages (R) for a HRO (i.e., calf: raising stage R1, growing heifer: raising stages R2 to R8, and pregnant heifer: raising stages R9 to R12).
<sup>b</sup>N/A: Not applicable.
<sup>c</sup>Weight estimation based on a 630-day-old heifer.

The operating costs were based on the daily cost of heifer-raising (including labor, veterinary treatment unrelated to *S.* Dublin, feed, equipment maintenance, and bedding; *HCo_n_*), additional treatment costs in clinically ill individuals (*TCo_w_* and *TCo_g_*), and rendering of dead animals (*RCo*; Fig 4, Table 3). Treatment costs were included based on prices from The Cornell Ambulatory and Production Service Clinic provided by F.A.L.Y, a veterinarian and epidemiologist with more than 10 years of experience working with dairy cattle and raising cattle operations. *HCo_n_* varied according to the raising stage (subindex *n* in *HCo_n_* represents raising stages from R1 to R12) and decreased with increasing herd size through a logarithmic function (Table 3) [52, 53]. Rendering of dead animals results in costs (*RCo*) without generating any income for the HRO. If vaccination was carried out, the vaccination cost (*VaCo*) covering labor and doses was included. For cleaning, costs were considered for each additional barn floor cleaning per week or per day (*CCo*).

The income derived from raising a heifer (*DIn*) was obtained based on expert opinion from the same veterinarian and epidemiologist who provided information on treatment costs (F.A.L.Y). The operating income was expressed as USD/100-head by the end of the first and second years into the simulation. The proportional change in operating income was calculated by comparing operating incomes for a specific control strategy and a “do-nothing” approach (baseline level of cleaning (1x per week) and no vaccination) over the 2-year simulation period. The consequences of milk yield reductions due to *S.* Dublin infection in the operating income were not assessed as heifers calve after leaving the HRO.

The module considers that calves enter the operation at 90 days of age (weaning age) and become pregnant after 360 days in the operation (at ∼15 months of age; [57]). In cases where cattle died prior to returning to their farms of origin, the operating cost was calculated based on the days spent in the operation until the raising stage prior to their death, plus half of the duration of the raising stage at the time of their death. For example, if an individual died in the seventh raising stage, it would mean that it completed raising stages from one to six (270 days) plus half of the seventh raising stage (22.5 days), adding up to a total of 292.5 days in the HRO.

### Operating income under different cost scenarios

Given the lack of information regarding the specific costs of applying vaccination and increasing cleaning frequency, a scenario analysis was carried out to determine the operating income of a HRO under different hypothetical costs of a mitigation strategy. For the cost assessment of vaccination, we considered that two doses are required for full vaccination status during the first year (i.e., two doses to effectively vaccinate each individual) and revaccination with a single dose after one year in the HRO. The cost for each of the cleaning scenarios represents the hypothetical cost (energy, labor, equipment maintenance) of an additional cleaning event beyond the cost at the frequency of cleaning once per week (baseline frequency). Estimations from the economic assessment were corroborated by the veterinarian expert in dairy farming who provided information on treatment costs and income (F.A.L.Y).

### Model running

We introduced stochasticity into the model through Monte Carlo simulation. This approach involves running multiple iterations of the model repeatedly and, in each iteration, specific values for parameters are randomly drawn from specified probability distributions to mimic the variation found in natural biological systems [58]. Each simulation consisted of 1,000 iterations. This number of iterations was chosen based on recommendations in Winston [59] and because the mean, 2.5^th^ and 97.5^th^ percentiles, and coefficient of variation (CV) of outcomes (the cumulative numbers of individuals in *A* and *I* compartments, carriers, and recovered individuals by the end of a 2-year long simulation) were similar under 5,000 and 1,000 iterations (i.e., the differences in the mean number of individuals were within ±2 and the CV was within ±0.5; Table S1). The R script and accompanying notes are available for open use at: https://github.com/IvanekLab/SDublin.git

### Model validation

Since there are no available reports on *S*. Dublin spread in HROs, the model was validated by comparing predicted seroprevalence with data from a dairy farm in Tennessee affected by *S*. Dublin [60]. Predictions were obtained for multiple scenarios in which a different assumed number of infected pregnant cattle entered the operation to reflect the different possibilities of infection introduction into this Tennessee dairy farm. Predictions from the model were considered valid if its interquartile range (IQR) encompassed the true seroprevalence value and its median predicted seroprevalence value was within the 95% confidence interval around the average true seroprevalence (Appendix S1).

### Sensitivity analysis

Sensitivity analysis was done using Partial Rank Correlation Coefficient (PRCC) with the R package *epiR* [61]. The analysis examined the relationship between input parameters (*D_A/I_*, *D_c_*, *D_R_*, *d_w_*, *d_g_*, *a_A/I_*, *a_C_*, *u_c_*, *u_g_*, *u_p_*, and *z*) and outcomes from the epidemiological and economic modules by the end of a 2-year simulation period. PRCC allows for determining the strength and direction of a certain parameter’s association over a model outcome of interest while controlling for the effect of other parameters included in the model [62]. Correlations were considered statistically significant based on Bonferroni-corrected *p-*value (significance at *p* < 0.005 estimated as 0.05/11 model parameters) to reduce type-I errors (e.g., [63]) and only correlation coefficient (ρ) values > |ρ|=0.4 were deemed relevant for reporting (e.g., [64]). Results from the PRCC analysis were visualized using a heat map. Classification trees were built in R using the *caret* package [65] to understand better the influence of the interaction among model parameters on the probability of an outbreak (defined as the presence or absence of >= 2 clinical cases over the length of the simulation). A 10-fold cross-validation procedure was carried out to assess the classification model performance (i.e., accuracy and agreement between predicted and expected classifications). Tree post-pruning was performed using the Cost-Complexity Pruning approach to achieve the simplest yet best-performing tree that accurately represents the simulated data [66].

### Scenario analysis

The importance of increasing the cleaning frequency, implementing vaccination, seasonality by means of temperature variation, herd size, and assumptions involving model parameters were assessed in scenario analysis (Table S2).

## Results

The developed model successfully replicated the 41% true seroprevalence estimated from the apparent prevalence in Kent et al. [60], assuming testing at 20, 25, 30, 35, or 40 days post the initial introduction of a single or multiple asymptomatic and carrier pregnant heifers (Appendix S1), thus supporting the validity of the model.

> *Salmonella* Dublin shedding amount in feces, infectious period, and time spent in the Recovered compartment influenced epidemiological outcomes and the operating income the most, whereas the outbreak probability was mostly determined by the interaction between the *S.* Dublin shedding rate and duration of the infectious period

Among the most influential parameters assessed in the sensitivity analysis, *S.* Dublin shedding amount in feces (*z*) and the duration of the infectious period (*D_A/I_*) were positively correlated with all epidemiological outcomes (deaths, abortions, carrier departures, and asymptomatic departures) and negatively correlated with the operating income (Fig 5). A strong correlation in the opposite direction was observed for the duration in the *R* compartment (*D_R_*) as it was negatively correlated with epidemiological outcomes and positively correlated with the operating income (Fig 5). Findings from the classification tree highlight the interaction between the *S.* Dublin shedding amount in feces and the duration of the infectious period as the primary predictors for the probability of an *S*. Dublin outbreak following the introduction of an asymptomatic heifer into a HRO (Fig 6). Specifically, the classification tree indicated that there was a <1% probability of an outbreak if the *S.* Dublin shedding amount was <2.2*10^3^ cells/g of feces and the duration of the infectious period was less than 32 days or if the duration of the infectious period was < 9.1 days and the shedding amount was <5.5*10^3^ cells/g.

**Fig 5.**
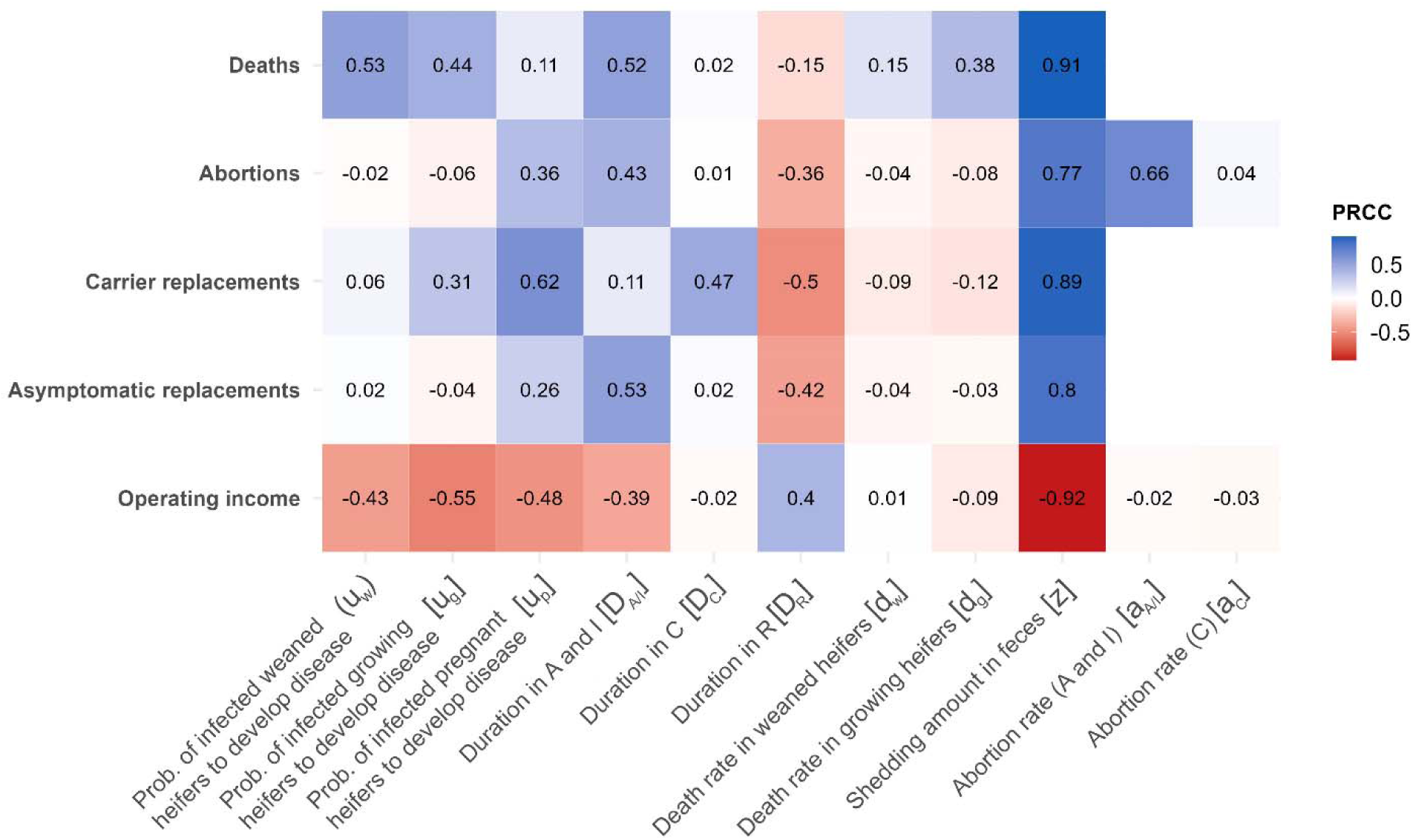
Findings from the sensitivity analysis. Heat map showing the Partial Rank Correlation Coefficient (PRCC) for the correlation between the model parameters (columns) and outcomes (rows), namely, deaths, abortions, and carrier and asymptomatic replacement heifers, and the operating income of a heifer-raising operation (Bonferroni corrected-*p*<0.005; |ρ|>0.4). *A*=asymptomatic, *I*=clinically ill, *C*=carrier, and *R*=recovered.

**Fig 6.**
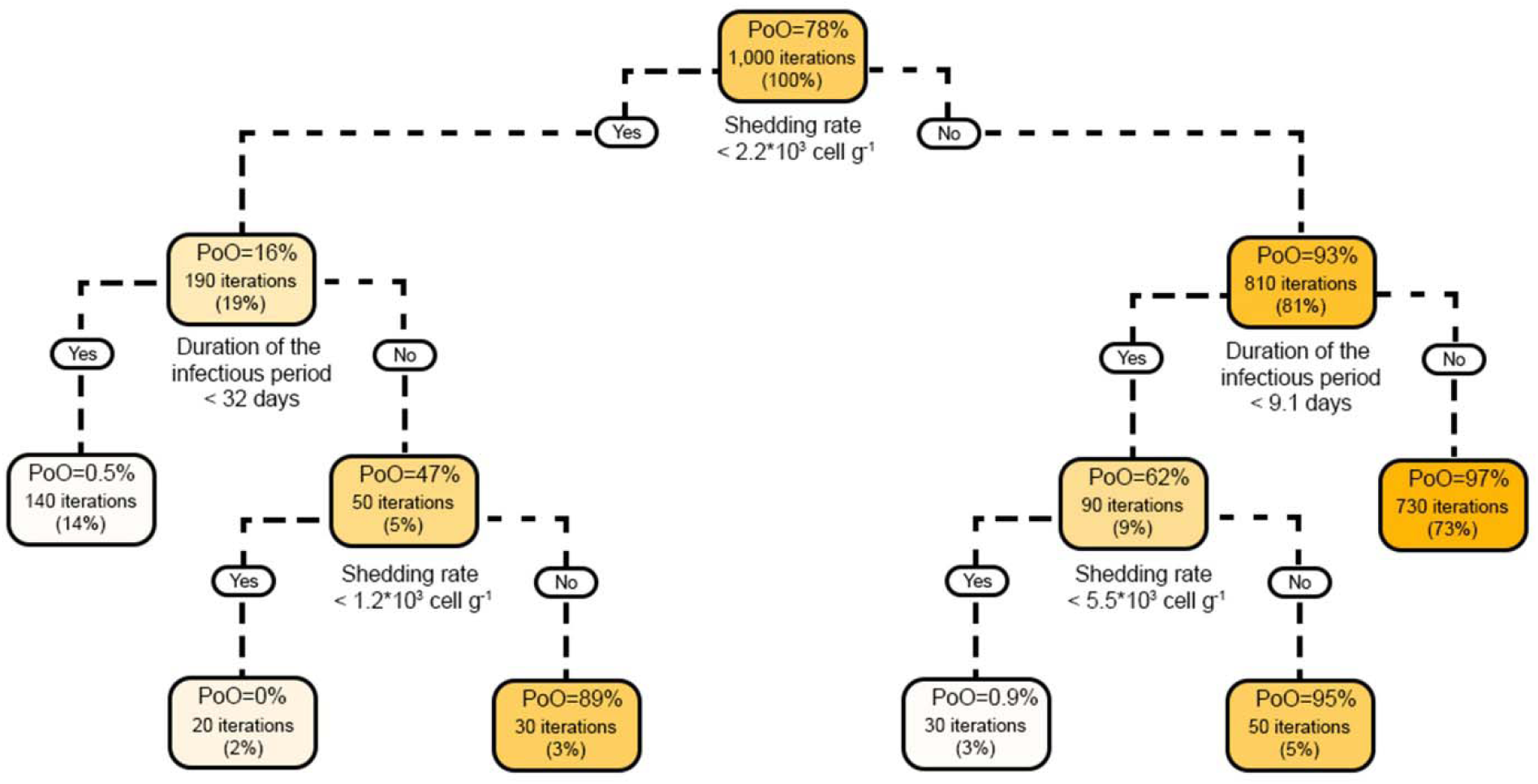
Classification tree of the probability of an outbreak after the introduction of an asymptomatic index case at time t=0. PoO=probability of an outbreak (the proportion of iterations with two or more *S*. Dublin clinical cases following the introduction of an asymptomatic index case). Higher color saturation in the boxes indicates an increased probability of an outbreak.

### Vaccination against *Salmonella* Dublin reduces deaths while increasing the cleaning frequency reduces all values for epidemiological outcomes

Findings indicate a 79% (785/1,000 iterations) probability of an outbreak following the introduction of an asymptomatic individual into a HRO housing 1,000 heifers when cleaning is performed once per week (baseline model; Table S3). Within the first year of the simulation, HROs affected by *S.* Dublin experienced medians of 20 deaths (IQR=9–35), 10 abortions (IQR=5–15), 15 carrier replacement heifers (IQR=11–19), and 57 asymptomatic replacement heifers (IQR=34–84) (Fig 7, Table S3). Vaccination was a very effective standalone control measure in decreasing median deaths, with a 75% reduction (median=5 deaths, IQR=2–10) compared to the baseline scenario. Cleaning at the highest frequency assessed (12x per day) reduced the outbreak probability to 23% (234/1,000 iterations), median deaths to 4 (80% reduction, IQR=0–10), median abortions to 4 (60% reduction, IQR=1–9), median carrier replacement heifers to 5 (67% reduction, IQR=0–9), and median asymptomatic replacement heifers to 37 (35% reduction, IQR=9–64). The scenario combining improvements in cleaning (12x per day) and vaccination showed a meaningful reduction in deaths (95% reduction, median=1, IQR=0–2), abortions (50% reduction, median=5, IQR=2–9), and carrier replacement heifers (67% reduction, median=5, IQR=1–10), with a less pronounced reduction in asymptomatic replacement heifers (23% reduction, median=44, IQR=15–71) compared to the baseline. It is important to note that scenarios involving vaccination showed a slight increase in carrier and asymptomatic infections, thereby marginally countering the impact of cleaning improvements on preventing infection spread to other operations (Fig 7, Table S3).

**Fig 7.**
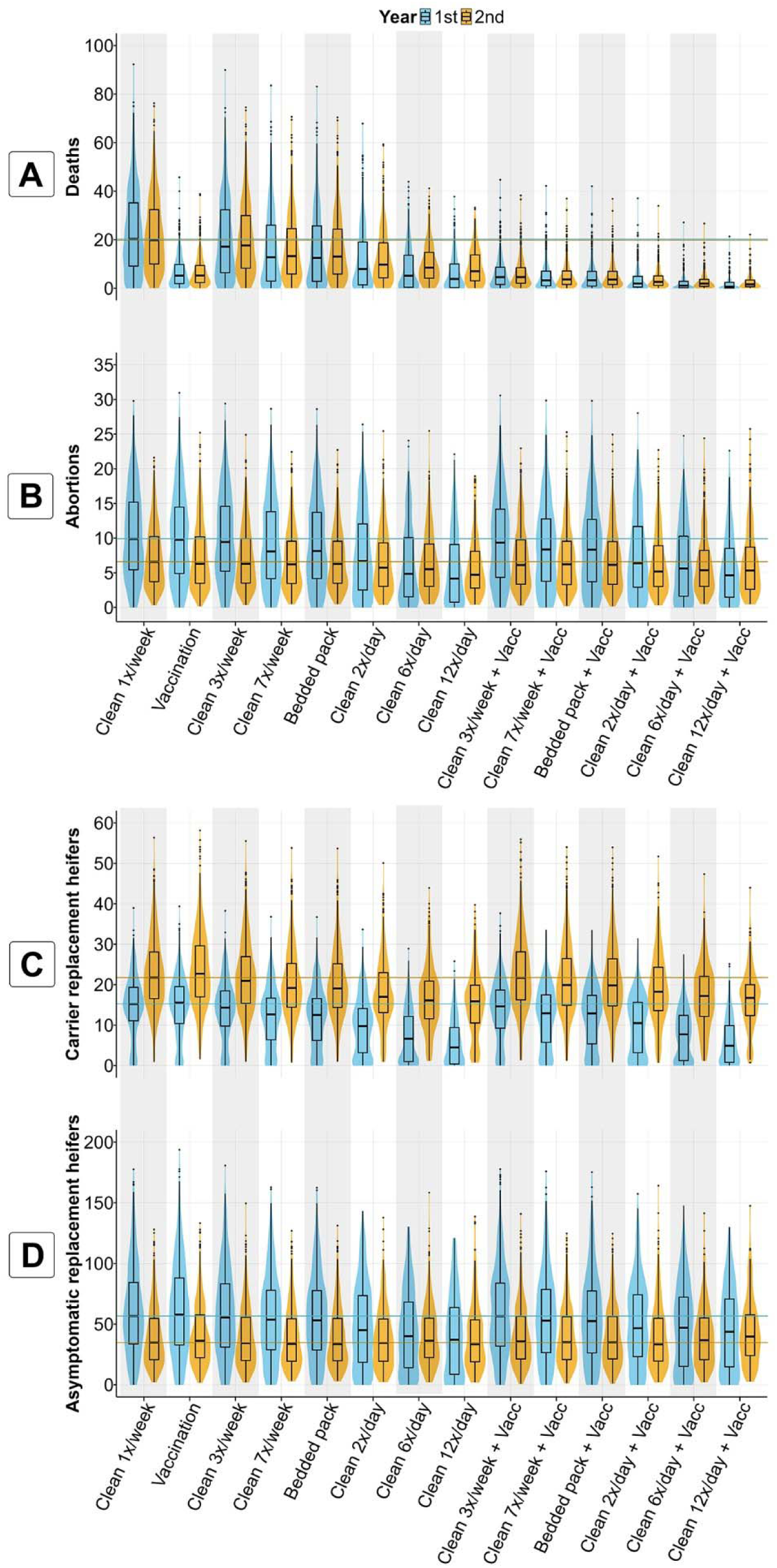
*Salmonella* Dublin-related (A) deaths and (B) abortions and (C) carrier and (D) asymptomatic infections in replacement heifers observed in scenarios with and without the implementation of vaccination or/and increased cleaning frequency in an HRO with N=1,000 heifers at the simulation start. A comparison was made between the baseline scenario (baseline level of cleaning (1x per week) and no vaccination) and scenarios in which control strategies were implemented. These strategies included cleaning at different frequencies per week (3x, 7x) and per day (2x, 6x, and 12x) and/or vaccination reducing mortality due to *S.* Dublin (vaccine effectiveness: median=0.72, 5^th^–95^th^ percentile=0.48–0.91). Horizontal blue and orange lines in each plot indicate the median value of the corresponding outcome under the baseline scenario.

### Stringent cleaning can decrease the impact of *S.* Dublin on the operating income of a HRO, even when it carries a high cost of implementation

In the absence of *S*. Dublin infection, a HRO with 1,000 heifers achieves an operating income of USD 39,123 per 100-head raised each year of production. This value decreases to a median of USD 32,827 (IQR=30,650–34,875) and USD 33,517 (IQR=31,443–35,179) per 100-head for years one and two, respectively (i.e., a median reduction from the initial operating income of 16% for year one and 14% for year two) after an infectious calf introduces *S.* Dublin into the herd and no mitigation strategies are implemented. Findings from our economic module indicate that for the cost scenarios considered, increasing cleaning frequency from 1x/week to 7x/week can increase the median operating income of a HRO post *S*. Dublin introduction between 2.7% to 3.8% during the first year (Fig 8, Table S4). Meanwhile, increasing the cleaning frequency to 12x/day (the most stringent cleaning evaluated in our study) can increase the median operating income of a HRO by 1.2% to 10.6% if the cost of additional daily cleaning per 100-head does not surpass USD 0.45 (Fig 8, Table S4). Furthermore, vaccination can increase the operating income by 1.2% to 2.4% compared to a “do-nothing” scenario during the first year only if costs per 100 vaccine doses do not surpass USD 100 (Fig 8, Table S4). Stringent cleaning measures combined with vaccination, did not improve the operating income in most cost scenarios (maximum operating income increase of 11.3% during the first year when vaccination does not require an additional investment; Table S4, Fig S2).

**Fig 8.**
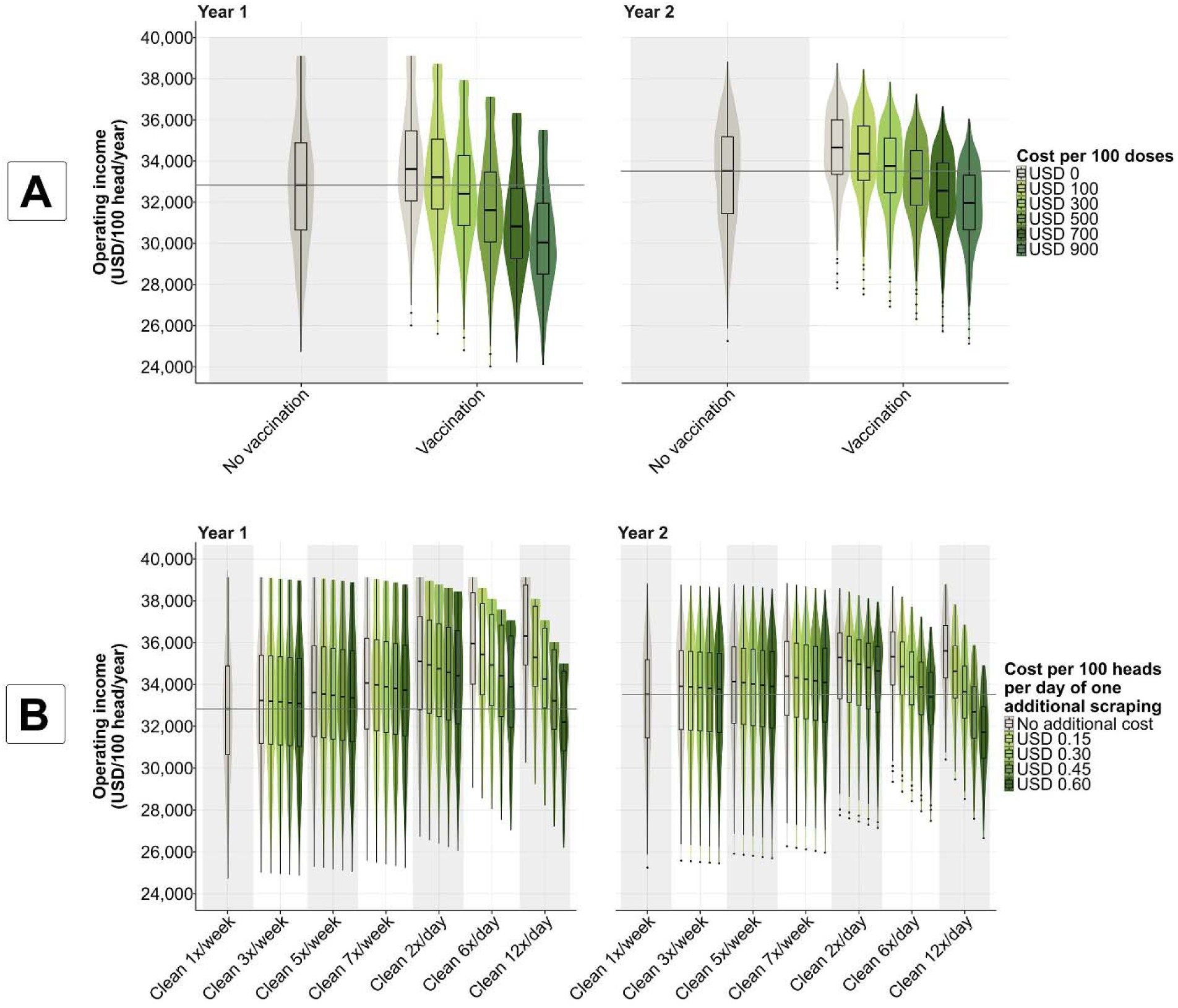
Operating income (USD per 100-head) observed in scenarios with different costs required for the implementation of mitigation strategies. A comparison was made between the baseline scenario (cleaning once (1x) per week and no vaccination), and thus no additional costs, and scenarios in which control strategies were implemented and compared in terms of the operating income (USD per 100-head). These strategies included (A) vaccination reducing *S.* Dublin-related mortality (vaccine effectiveness: median=0.72, 5^th^-95^th^ percentile=0.48–0.91) and (B) cleaning at different frequencies (3x, 5x, and 7x times per week and 2x, 6x, and 12x per day). Findings were divided into the first year and the second year of the simulation. Operating income in a “no infection” scenario = USD 39,123 per 100 head each year of the simulation. Horizontal gray line in each plot-year indicates the median value of the operating income under the baseline (cleaning once (1x) per week and no vaccination).

Findings from the scenario analysis also indicate that reduced shedding levels in asymptomatic and carrier individuals relative to clinically ill meaningfully influenced all epidemiological outcomes (Table S3). Furthermore, clinical cases are critical to triggering a *S.* Dublin outbreak and its epidemiological consequences in HROs (Table S3).

## Discussion

We developed a mathematical model that integrates available knowledge of the epidemiology of *S*. Dublin in HROs and conducted an *in silico* assessment of the effectiveness of cleaning improvements and vaccination as control strategies. *S.* Dublin isolates from cattle in US dairy farms have displayed resistance to multiple antibiotics [5, 42, 67]. Considering that no differences in transmission between susceptible and MDR *S.* Dublin have been identified to date, the model presented in our study can be considered to represent the transmission of both susceptible and MDR *S.* Dublin strains. However, economic impacts would likely be larger for MDR strains given the higher treatment costs and potentially prolonged infections associated with MDR strains. The main study findings are: (i) HROs play an important role in the spread of *S.* Dublin to dairy cattle herds; (ii) frequent cleaning meaningfully reduced the probability of *S.* Dublin introduction into HROs and its consequences, while also improving the operating income of these operations depending on the implementation costs; (iii) vaccination greatly prevents deaths from *S.* Dublin infection, but does not prevent its transmission; and (iv) *S*. Dublin presence in HROs can be sustained through the continuous arrival of naïve individuals, not requiring the repeated introduction of carriers, while the occurrence of clinical disease in infected heifers promotes *S.* Dublin spread in the operation. We discuss the implications of those findings and study limitations in the following sections.

### Novel mathematical model of *S.* Dublin transmission and its implications within the context of HROs

Our mathematical model of the transmission dynamics of *S.* Dublin in a US HRO represents a novel addition to an array of models developed previously for *Salmonella* spp. in cattle herds [8, 11, 15] but not in HROs. The model assessed the epidemiological impacts of *S*. Dublin on HROs by estimating cattle deaths and abortions and identified the role that these operations might be playing as disseminators of the infection to dairy herds. Furthermore, hypothetical cleaning improvements and vaccination were evaluated, providing insight into the expected epidemiological population-level impacts of these control strategies, if accessible to farmers. The economic module allows exploring the consequences of *S.* Dublin in HROs and determining under what conditions vaccination and cleaning improvements could help mitigate the economic impacts of these effects. This is relevant as the economic feasibility of implementing mitigation strategies against *S*. Dublin has not been explored in the US. Overall, we expect the model to be used by stakeholders to predict the epidemiological and economic consequences of *S.* Dublin in HROs.

### HROs play a relevant role in the dissemination of *S.* Dublin to other herds

The model yielded meaningful insights into the contribution of HROs in the spread of *S.* Dublin to dairy farms by estimating the number of departing asymptomatic and carrier replacement heifers over a 2-year period, encompassing the entire raising period of 540 days. The number of asymptomatic infections among replacement heifers departing the HRO was particularly relevant during the first year when the herd was entirely naïve to *S*. Dublin, while the number of carriers that departed back to their farms of origin increased during the second year once multiple individuals had been infected. These predictions suggest that affected HROs will disseminate *S.* Dublin in the short and long term if the pathogen is not removed from the operation. The use of HROs by dairy farmers has been previously pointed out as a risk factor for introducing bacterial pathogens, including *Salmonella* [68]. Furthermore, Edrington et al. [69] indicated that 80% of fecal samples collected from heifers in a HRO were *Salmonella* positive right before returning to their farm of origin at 24 months old. Although our model predicted that the development of carriers during the first year could be reduced by more than half with stringent cleaning measures, the occurrence of asymptomatic individuals was less sensitive to this control strategy. This observation is likely due to the influx of susceptible individuals arriving in batches that are exposed to environmental contamination with the pathogen, allowing the infection to persist in the herd. These findings emphasize that HROs free of *S*. Dublin should prioritize preventing the introduction of the pathogen, as once established, even stringent cleaning measures may be insufficient to eradicate it and prevent its spread to other farms. Furthermore, since most infected individuals do not become carriers and remain infectious for about two weeks [9, 23], these findings suggest that strategies such as testing cattle prior to returning to their original farms and quarantining animals upon arrival at the farm could help prevent the introduction of *S*. Dublin to farms using off-site heifer raising. Nonetheless, this approach would first require the improvement of diagnostics for *S*. Dublin infection in cattle, given the reported low sensitivity of available tests [43, 70].

### Vaccination using a hypothetical vaccine reduces *S*. Dublin-induced mortality but slightly contributes to infection persistence in the herd

Reducing mortality resulting from *S*. Dublin infection allows farmers to have heifers ready to return to their farms of origin after the production period. However, findings from the model revealed possible adverse epidemiological consequences of vaccination based on the current knowledge of their effectiveness (Fig 3). Since the vaccine does not prevent infection, nor shorten the duration of the infectious period, the infection continues to spread through the vaccinated herd (even better than in the absence of vaccination due to prevented deaths of infected individuals). Vaccinating infectious individuals to prevent them from dying eventually led to a slightly higher number of long-term carriers during the first and second years after the initial introduction of *S.* Dublin and the departure of a larger number of replacement heifers with an asymptomatic infection during the second year compared to baseline. The use of vaccination also led to a reduction in the effectiveness of cleaning in preventing the development of carrier and asymptomatic individuals (Table S3). A similar finding was observed by Lanzas et al. [13], who indicated that halving the probability of death after *Salmonella* infection in calves increased the median number of infected individuals. Overall, the silent spread of *S.* Dublin increases the level of infection in the herd, leading to the dissemination of the pathogen to other farms and the zoonotic risk to farm workers. Currently, the only confirmed effect of vaccines observed in practical settings is a decrease in the likelihood of death in young cattle after an extra-label administration [42], while a reduction in *S*. Dublin symptoms among vaccinated calves has been explored but lacks supporting evidence [41, 42]. Therefore, there is a need for further vaccine research and development to enable the use of vaccination in infection control.

### Stringent cleaning reduces *S*. Dublin outbreaks, helps prevent deaths and abortions, and limits the departure of infectious individuals to their farm of origin

Our study findings demonstrate that increasing the frequency of cleaning in the barn through multiple scrapings per day can decrease the likelihood of a *S*. Dublin outbreak and its consequences (i.e., deaths, abortions, and asymptomatic and carrier infections among replacement heifers) following the introduction of an infectious individual. This improvement is based on the continuous removal of feces, which further limits the chances of cattle contacting *S*. Dublin in the environment as the cleaning frequency increases. This is supported by findings from the classification tree analysis indicating that the amount of *S.* Dublin shed in feces and the length of shedding are the main predictors of an outbreak post-introduction of an index case. Previous studies have highlighted the importance of feces removal from the environment as a way to prevent the occurrence of secondary cases of *Salmonella* [8, 11, 15] and limit the duration of outbreaks [13]. In their study, Nielsen et al. [11] found that improving hygiene on a farm, understood as a reduction in the probability of infection from the environment, reduced by more than half the probability of an outbreak and meaningfully decreased the duration and size of the epidemic. Similarly, Xiao et al. [8] found that frequently and effectively removing feces from the barn meaningfully contributed to controlling *Salmonella* infection by limiting its indirect transmission among cattle. Recent findings from a cross-sectional study done by Perry et al. [71] in Ontario dairy farms contradict predictions from our model by indicating that frequent manure removal from the calving area increases the risk of *S.* Dublin infection, suggesting that bacteria might spread across the barn during the manure removal process. However, Perry et al. [71] acknowledge that their findings might be a false positive related to farmers adopting frequent cleaning in the calving area after experiencing a *S*. Dublin outbreak. Nevertheless, the potential positive and negative impacts of floor scrapers on the spread of enteric bacteria in the barn need further investigation [72].

Unlike prior modeling efforts assessing hygiene as a control strategy, our approach provides a more realistic evaluation by accounting for the weekly or daily cleaning frequency of the barn’s floor, representing observed or hypothetical cleaning practices on dairy farms. Further research is warranted to comprehensively understand the impact of cleaning on preventing *S*. Dublin outbreaks and reducing transmission. This should consider various management measures beyond increasing cleaning frequency, evaluated while considering animal welfare and economic factors. Additionally, relying solely on cleaning is not sufficient to fully protect HROs from an outbreak and control the spread of the pathogen. Therefore, we suggest the adoption of a comprehensive approach to disease management that includes measures in addition to improvements in cleaning, such as limiting the number of farms providing heifers to the HRO, isolation and quarantine practices when heifers arrive at the operation, and efforts to early detect and remove *S.* Dublin-infected cattle.

### HROs experience a meaningful reduction in operating income following the introduction of *S.* Dublin, which can be partly prevented via stringent cleaning practices

Available studies assessing economic losses associated with *S.* Dublin occurrence in dairy farms have been restricted to European countries [27, 73, 74, 75]. Our study found that *S*. Dublin meaningfully impacts a HRO’s median operating income following the introduction of an infectious individual (Fig 8, Table S4). This reduction in the operating income is a consequence of increased costs of treating severely diseased heifers, raising individuals that end up dying in the operation, the opportunity cost of selling heifers for meat instead of returning them pregnant to their farm of origin, and the expenses related to the rendering of dead animals.

Our findings indicate that the reduction in mortality via a vaccine with the characteristics considered in our model is not an effective standalone approach to prevent *S.* Dublin dissemination in a HRO (i.e., raising heifers from 90 to 630 days of age). Although the reduction in mortality due to vaccination increased the HRO’s income by reducing heifer losses, in most scenarios this benefit was offset by the cost of administering the vaccine. It is important to note that the vaccine’s effectiveness in improving the operating income may be more relevant in production settings that include newborns, such as calf raisers and some HROs, as they are more susceptible to experiencing severe symptoms and dying from the infection. It is also important to highlight that the Gram-negative modified-live nature of the assessed vaccines can lead to anaphylactic episodes in young cattle [76]. Although these anaphylactic events have been described as “low”, “occasional”, or “anecdotal” in the literature [41, 76, 77], communications with dairy producers and veterinarians describe these events as a serious concern (as indicated by E.F.). The rate at which these anaphylactic events occur among individuals of different ages remains to be systematically evaluated, as well as its economic relevance for heifer raisers.

In most scenarios, daily cleaning appeared as a more economically feasible approach to improving the operating income of a HRO affected by *S.* Dublin compared to the sole implementation of vaccination and cleaning on a per week basis, particularly during the first year. This is the result of preventing mortality and, in contrast to vaccination, reducing the occurrence of severe cases and the overall level of infection in the farm through the frequent removal of feces. Moreover, stringent cleaning measures greatly reduced the chance of an outbreak in the HRO, which should be prioritized to prevent financial losses due to *S.* Dublin altogether. When cleaning was done on a weekly basis, the improvements in operating income were much less noticeable, indicating that cleaning on a per day basis is a more effective approach for both reducing the infection level and increasing the operating income of a HRO. Our findings from the economic assessment of cleaning improvements conflict with the assessment by Bergevoet et al. [75], who characterized hygienic measures as the least cost-effective approach for controlling *Salmonella* (serovars Typhimurium and Dublin) compared to other strategies such as segregating cattle by age groups and preventing the introduction of new animals, although in that study it still reduced prevalence by half over a period of three years. However, it is important to note that Bergevoet et al. [75] assessed cleaning measures solely in terms of reducing the probability of developing chronic infection (i.e., carrier state), without considering the broader significance of indirect transmission in *Salmonella* dynamics. On the other hand, Nielsen et al. [27] found that dairy farms applying very good management practices, including excellent hygiene, had the highest improvements in profit. The improvement in the operating income observed when cleaning on a per day basis, particularly in scenarios with lower costs and frequent cleaning, suggests that innovations allowing farmers to clean cheaper and better can meaningfully improve a HRO’s profitability.

We acknowledge that our assessment of cleaning frequency did not consider the potential negative effects on cattle behavior and comfort, which could lead to negative financial consequences. For example, Cramer et al. [78] and Crossley et al. [79] have found that increased cleaning frequency is associated with a higher occurrence of tail and hoof lesions in dairy cattle. While increased cleaning frequency may help mitigate *S*. Dublin, particularly alongside other measures, a better understanding of the effects of frequent cleaning (e.g. cleaning with an alley scraper multiple times per day) on cattle productivity and well-being would improve economic estimations.

### Carriers are relevant for the initial introduction of *S*. Dublin into HROs but have minimal impact on transmission dynamics once the infection reaches an endemic state

Findings from the scenario analysis revealed that carriers play a meaningful role in the initial introduction and spread of *S*. Dublin in a HRO, however, their contribution in spreading the pathogen once the infection has been introduced in the herd is rather minor. In fact, the long-term persistence of *S.* Dublin in a HRO was primarily attributed to the occurrence of clinical cases within the herd and the continuous arrival of susceptible individuals. This finding aligns with the study by Lanzas et al. [12], which showed that long-term shedders and subclinical individuals with lower *Salmonella* shedding had a minimal impact on the occurrence of infection cases in a dairy farm. The findings of Nielsen et al. [11] challenge this notion by highlighting the essential role played by carriers in the long-term persistence of *S*. Dublin in dairy herds. The disparity in carrier importance between their model and ours could stem from their consideration of increased resistance to reinfection in previously infected individuals, highlighting the significance of persistent shedders in sustaining the infection within a growing population of individuals with reduced susceptibility. Additionally, the HROs have a faster turnover rate compared to dairy farms (animals typically spend ∼1.3 years in a HRO, whereas they spend ∼5 years in a dairy farm [19, 80]), which might have also contributed to the difference in the relative importance of long-term shedders in sustaining the disease within these operations.

The limited role of carriers in spreading *S*. Dublin within an HRO suggests that a testing-and-culling approach targeting carriers is likely ineffective for controlling the infection. Despite efforts to identify and eliminate persistent shedders in a high-risk operation, incoming cattle may still become infected and develop a carrier state due to exposure to *S*. Dublin from other infectious cattle. Furthermore, the intermittent shedding of *S*. Dublin by carriers and the limitations of testing methods in detecting ongoing infections make it difficult to control this pathogen in HROs through testing-and-culling strategies [81, 82]. As a result, our model supports previous findings that targeting carriers alone is unlikely to be a cost-effective strategy for managing *S*. Dublin in HROs.

### Identified knowledge gaps in *S.* Dublin epidemiology

The duration of immunity to *S.* Dublin infection (*D_R_*) was identified as an important parameter determining model outcomes. This has also been reported by other mathematical modeling studies [8, 12, 15]. Unfortunately, the duration of the period included in our model is based on the humoral response of cattle [23, 26], which does not necessarily indicate protection against *S.* Dublin reinfection. Indeed, the cellular response is the primary means of protection against *Salmonella* infection, with antibodies playing a secondary role [70].

In our study, we did not consider super shedders (i.e., individuals that shed into the environment a much larger amount of *Salmonella* Dublin than other infectious individuals and for an extended period) in the transmission of *S.* Dublin due to its importance still being under discussion. For instance, Lanzas et al. [12] highlighted super shedders as important for transmitting *Salmonella* in dairies, with its occurrence leading to an increase in the number of secondary infections. In contrast, Nielsen et al. [11] indicated that super shedders are not an essential component in *S.* Dublin epidemiology, with their occurrence being extremely rare. In both studies, super shedders were modeled as infectious individuals shedding the same amount of *Salmonella* into the environment as clinically infected cattle but for an extended period. While our model did not explicitly consider super shedders, scenario analysis evaluated the impact of carrier individuals shedding the same amount of *S.* Dublin as clinically ill cattle (Table S3). The results revealed a moderate difference in epidemiological outcomes, supporting that the super shedders could have a meaningful impact on *S*. Dublin transmission dynamics.

Another important knowledge gap is the dynamics of intermittent shedding, i.e., the frequency with which carriers interrupt and restart their shedding of *S.* Dublin. Elucidating this would allow researchers to more precisely assess the role of carriers in *S.* Dublin dynamics in dairy cattle operations, since considering uninterrupted shedding (as we did in our model) may under- or overestimate their role as disseminators.

It is crucial to acknowledge the lack of information regarding the costs associated with implementing cleaning improvements and vaccination. Additional information is needed on the costs of using a skid steer or alley scraper at varying frequencies (e.g., per day or week) and across operations of different sizes to accurately determine their influence on the operating income of an HRO. Considering the benefits of cleaning improvements observed in our study, developing new technologies that enable frequent floor scraping at a lower cost could meaningfully improve the HRO’s operating income in the face of a *S.* Dublin outbreak. For vaccination, understanding its economic implications is essential, particularly regarding labor-related costs for its initial administration and revaccination, as well as potential side effects in young cattle. Addressing these knowledge gaps will be key to developing economically sound strategies for controlling *S*. Dublin transmission.

### Model limitations

Because data suitable for model validation from a HRO were not available, we validated the model using information from a dairy farm that experienced a *S.* Dublin outbreak after the introduction of cattle in batches from another dairy farm [60]. While not ideal, this was considered acceptable because of the similarities between *S.* Dublin epidemiology on dairy farms and HROs. As more *S.* Dublin data on HROs become available, the model parameters and validation analysis can be updated. The model did not account for increased resistance to *S*. Dublin infection among susceptible individuals who had previously been infected and returned to the susceptible compartment upon the loss of immunity [83]. This exclusion was due to the limited data for parameterization. The inclusion of this feature in the model could have reduced the incidence of *S*. Dublin reinfections in previously exposed cattle and ultimately decreased health impacts. However, our findings are relevant to a naive herd and thus represent conservative predictions. Subject to data availability, future modeling efforts should consider resistance to reinfection when emulating *S.* Dublin dynamics in cattle. The homogeneous mixing of cattle in a pen was chosen as a simplification for model parameterization, a choice also made by other *Salmonella* modeling studies [13, 15, 16]. The assumption of homogeneous mixing considered in our model contributes to increasing the chances of a single individual causing an outbreak and spreading disease. Despite this limitation, the validation process done to the model indicates that it is capable of predicting the level of infection observed in cattle farms, therefore although a simplification, it provides a valuable approximation to the situation in real life. Due to the compartmental nature of the model, vaccination was incorporated at the start of the simulation, rather than in response to an outbreak. The limitations of the model also prevented considering a two-week interval between vaccine doses, most likely leading to a slight overestimation of its effectiveness in preventing deaths among calves and young heifers. Additionally, we only evaluated scenarios where mitigation strategies, such as cleaning improvements and vaccination, were implemented simultaneously with the introduction of *S*. Dublin into a naïve herd. We did not assess these interventions implemented as preventative measures during times when the introduction of the infection into the herd is uncertain (not occurring at all or occurring with a low probability). The epidemiological and economic outcomes of these interventions would obviously be different in such scenarios compared to the results presented here. Analysis of such uncertainty scenarios was not conducted because it would require assumptions about the timing and scale of *S*. Dublin introduction events and the history of infection and immunity in the herd. While this presents a limitation, we believe the scenarios analyzed here are informative, considering that HROs receive animals from multiple dairy herds, increasing the risk of infection introduction with the most severe consequences for a naïve herd as assumed here.

### Conclusions

The developed mathematical model of *S.* Dublin transmission dynamics in a HRO predicts the epidemiological and economic consequences of *S.* Dublin infection in a US HRO over a 2-year simulation period, offering valuable insights for disease management. Our findings highlight the key role that HROs play in the dissemination of *S*. Dublin to other herds through the movement of asymptomatic and persistently infected individuals. Additionally, the model reveals that, although carriers are important in the initial introduction of *S*. Dublin into HROs, the short production period for heifers in HRO (less than two years) prevents carriers from meaningfully contributing to the persistence of the infection in the herd. Among evaluated control strategies, stringent cleaning practices substantially mitigate *S*. Dublin impacts on a HRO, preventing deaths and abortions, and limiting the departure of infectious heifers to their farm of origin. Meanwhile, vaccination with a hypothetical vaccine, capturing current knowledge of the effectiveness of commercial *S*. Dublin vaccines, reduces *S*. Dublin-induced mortality but also slightly contributes to infection persistence within the herd. Thus, based on the current information about these vaccines, alternative measures to vaccination are necessary to sustainably control *S*. Dublin on HROs. This also calls for more research into vaccines against *S.* Dublin infection. The study also demonstrated that *S*. Dublin introduction leads to a meaningful reduction in a HRO’s operating income, a loss that can be partly offset by implementing frequent cleaning. Knowledge gaps, including the cost of interventions, intermittent shedding dynamics, and the duration of immunity, remain barriers to fully understanding *S*. Dublin dynamics on HROs. Addressing these gaps is essential for designing more effective control measures against the pathogen and improving economic outcomes for HROs. Overall, our findings support the importance of addressing *S*. Dublin as a notable threat to dairy cattle production in the US and emphasize the need for control at HROs to prevent its dissemination to dairy farms.

## Acknowledgments

The authors would like to extend their appreciation to Dr. Pablo Pinedo, Dr. Angel Abuelo, Dr. Kaitlyn Kremer, Dr. Erin Goodrich, Dr. Eleni Casseri, Dr. Trent Westhoff, and Casey Havekes for their insights about cleaning effectiveness and vaccination. We very much appreciate the assistance of Dr. Peter Thomsen and Dr. Nanna Krogh Skjølstrup for the valuable information that they provided about Danish dairy farms.

This material is based upon work supported by the National Institute of Food and Agriculture, USDA (Washington, DC), Hatch, under accession number 7000433, as well as Multistate Research Funds, accession numbers 1016738 and 7005699 awarded to R. I. This project was also supported by the Cornell Institute of Digital Agriculture (CIDA) through the Research Innovation Fund (RIF) awarded to SL-S.

---

**Appendix S1. Details about the methodology and description of additional findings.**

**Appendix S2. Tool sheet used to determine cleaning effectiveness per day at different cleaning frequencies in a heifer-raising operation.** Information about cleaning effectiveness was obtained through expert elicitation.

**Fig S1.**
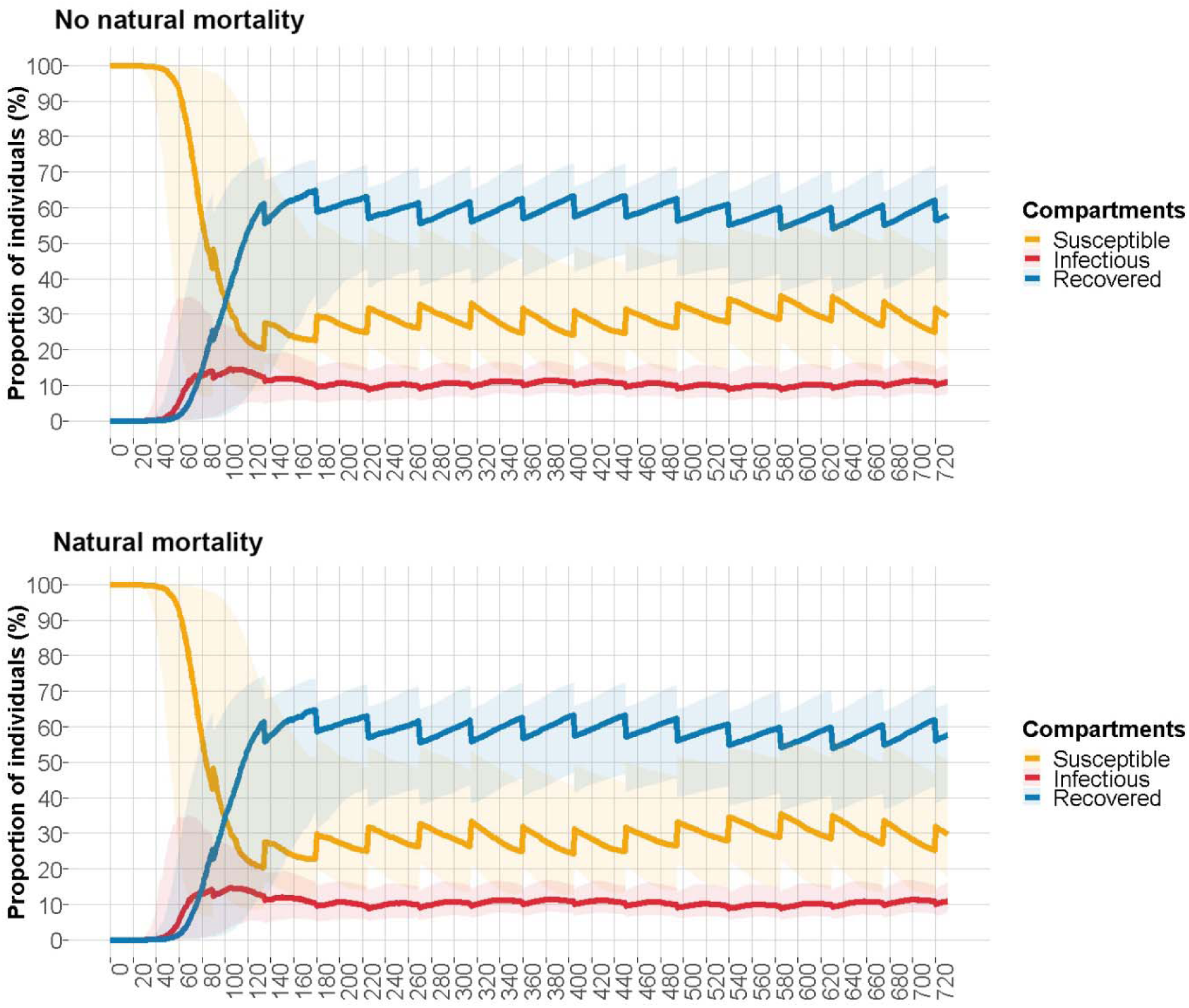
*Salmonella* Dublin dynamics in a heifer-raising operation with (top) and without (bottom) considering natural mortality among cattle.

**Fig S2.**
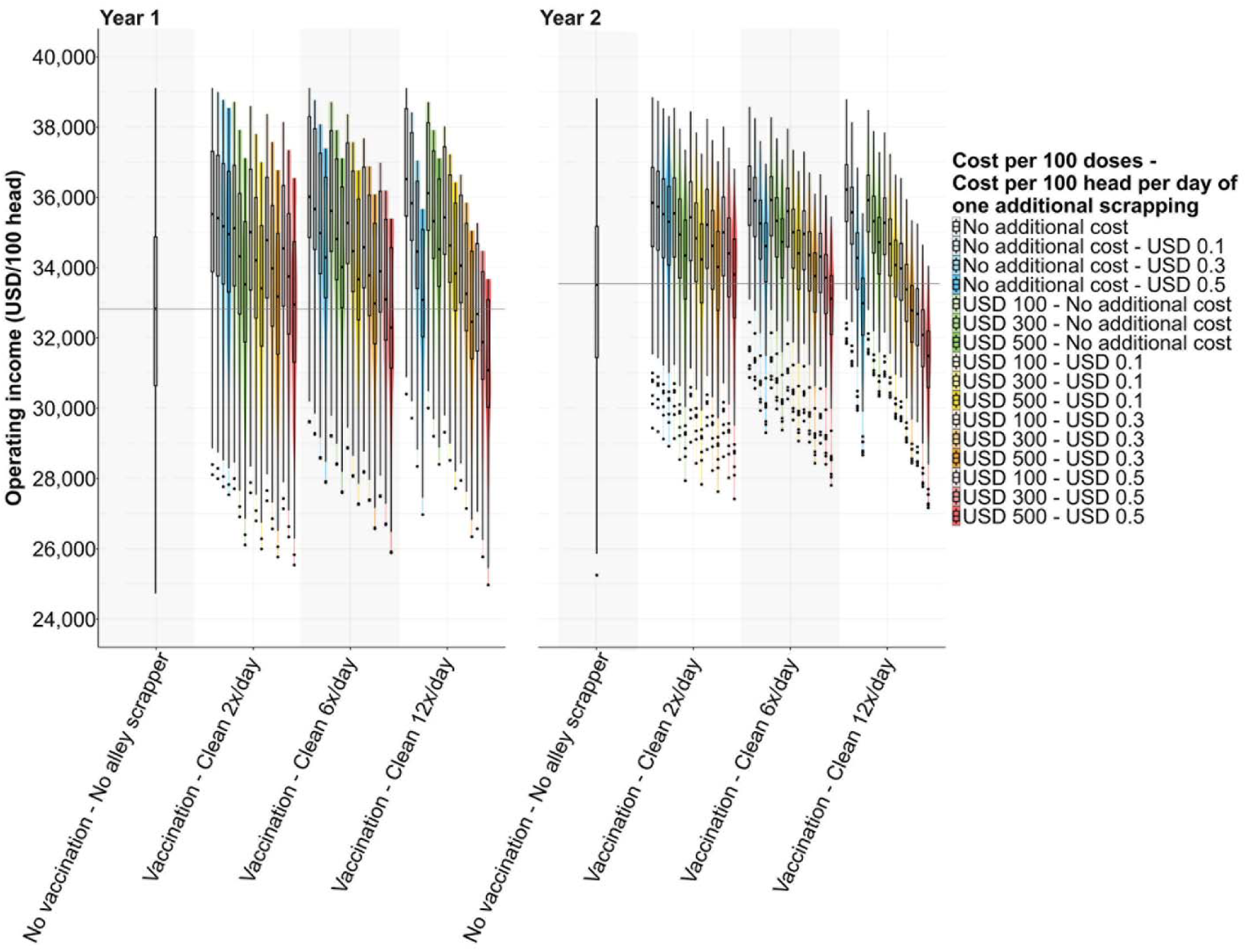
Operating income (USD per 100-head) observed in scenarios with different costs required for the implementation of mitigation strategies. A comparison in terms of the operating income (USD per 100-head) was made between the baseline scenario (cleaning once (1x) per week and no vaccination) and scenarios in which multiple control strategies were implemented. Observations for the first and second years into the simulation are presented in the figure. Horizontal grey lines in each plot indicate the median value of the corresponding outcome under the baseline scenario.

## References

1. Hegde NV, Cook ML, Wolfgang DR, Love BC, Maddox CC, Jayarao BM. Dissemination of *Salmonella enterica* subsp. *enterica* serovar Typhimurium var. Copenhagen clonal types through a contract heifer-raising operation. J Clin Microbiol. 2005;43: 4208–4211. doi: 10.1128/JCM.43.8.4208-4211.2005

2. Nielsen LR, Schukken YH, Gröhn YT, Ersbøll AK. *Salmonella* Dublin infection in dairy cattle: risk factors for becoming a carrier. Prev Vet Med. 2004;65: 47–62. doi: 10.1016/j.prevetmed.2004.06.010

3. Harvey RR, Friedman CR, Crim SM, Judd M, Barrett KA, Tolar B, et al. Epidemiology of *Salmonella enterica* serotype Dublin infections among humans, United States, 1968–2013. Emerg Infect Dis. 2017; 23: 1493–1501. doi: 10.3201/eid2309.170136

4. Hsu CH, Li C, Hoffmann M, McDermott P, Abbott J, Ayers S, et al. Comparative genomic analysis of virulence, antimicrobial resistance, and plasmid profiles of *Salmonella* Dublin isolated from sick cattle, retail beef, and humans in the United States. Microb Drug Resist. 2019;25: 1238–1249. doi: 10.1089/mdr.2019.0045

5. Srednik ME, Lantz K, Hicks JA, Morningstar-Shaw BR, Mackie TA, Schlater LK. Antimicrobial resistance and genomic characterization of *Salmonella* Dublin isolates in cattle from the United States. PLOS ONE. 2021;16: e0249617. doi: 10.1371/journal.pone.0249617

6. Davidson KE, Byrne BA, Pires AF, Magdesian KG, Pereira RV. Antimicrobial resistance trends in fecal *Salmonella* isolates from northern California dairy cattle admitted to a veterinary teaching hospital, 2002-2016. PLOS ONE. 2018;13: e0199928. doi: 10.1371/journal.pone.0199928

7. McMillan EA, Jackson CR, Frye JG. Transferable plasmids of *Salmonella* enterica associated with antibiotic resistance genes. Front Microbiol. 2020;11: 562181. doi: 10.3389/fmicb.2020.562181

8. Xiao Y, Bowers RG, Clancy D, French NP. Understanding the dynamics of *Salmonella* infections in dairy herds: a modelling approach. J Theor Biol. 2005;23: 159–175. doi: 10.1016/j.jtbi.2004.09.015

9. Nielsen LR, van den Borne B, van Schaik G. *Salmonella* Dublin infection in young dairy calves: Transmission parameters estimated from field data and an SIR-model. Prev Vet Med. 2007; 79:46–58. doi: 10.1016/j.prevetmed.2006.11.006

10. Chapagain PP, Van Kessel JS, Karns JS, Wolfgang DR, Hovingh E, Nelen KA, et al. A mathematical model of the dynamics of *Salmonella* Cerro infection in a US dairy herd. Epidemiol Infect. 2008;136: 263–272. doi: 10.1017/S0950268807008400

11. Nielsen LR, Kudahl AB, Østergaard S. Age-structured dynamic, stochastic and mechanistic simulation model of *Salmonella* Dublin infection within dairy herds. Prev Vet Med. 2012;105: 59–74. doi: 10.1016/j.prevetmed.2012.02.005

12. Lanzas C, Brien S, Ivanek R, Lo Y, Chapagain PP, Ray KA, et al. The effect of heterogeneous infectious period and contagiousness on the dynamics of *Salmonella* transmission in dairy cattle. Epidemiol Infect. 2008;136: 1496–1510. doi: 10.1017/S0950268807000209

13. Lanzas C, Warnick L, Ivanek R, Ayscue P, Nydam D, Gröhn Y. The risk and control of *Salmonella* outbreaks in calf-raising operations: a mathematical modeling approach. Vet Res. 2008;39: 61. doi: 10.1051/vetres:2008038

14. Jordan D, Nielsen LR, Warnick LD. Modelling a national programme for the control of foodborne pathogens in livestock: the case of *Salmonella* Dublin in the Danish cattle industry. Epidemiol Infect. 2008;136: 1521–1536. doi: 10.1017/S0950268807000179

15. Lu Z, Grohn YT, Smith RL, Wolfgang DR, Van Kessel JAS, Schukken YH. Assessing the potential impact of *Salmonella* vaccines in an endemically infected dairy herd. J Theor Biol. 2009;259: 770–784. doi: 10.1016/j.jtbi.2009.04.028

16. Lu Z, Gröhn YT, Smith RL, Karns JS, Hovingh E, Schukken YH. Stochastic modeling of imperfect *Salmonella* vaccines in an adult dairy herd. Bull Math Biol. 2014;76: 541–565. doi: 10.1007/s11538-013-9931-5

17. McBryde ES, Pettitt AN, McElwain DLS. A stochastic mathematical model of methicillin resistant *Staphylococcus aureus* transmission in an intensive care unit: predicting the impact of interventions. J Theoretical Biol, 2007;245: 470–481. doi: 10.1016/j.jtbi.2006.11.008

18. Jit M, Brisson M. Modelling the epidemiology of infectious diseases for decision analysis: a primer. Pharmacoeconomics. 2011;29: 371–386. doi: 10.2165/11539960-000000000-00000

19. United State Department of Agriculture (USDA). Dairy heifer raiser, 2011: an overview of operations that specialize in raising dairy heifers. USDA Animal and Plant Health Inspection Service (APHIS) - Veterinary Services (VS). 2011. Available from: https://web.archive.org/web/20230704160851/https://www.aphis.usda.gov/animal_health/nahms/dairy/downloads/dairyheifer11/HeiferRaiser_1.pdf

20. R Core Team. R: A language and environment for statistical computing. The R Foundation. 2014 [cited 2022 January 10]. Available from: http://www.R-project.org/

21. Soetaert K, Petzoldt T, Setzer RW. Solving Differential Equations in R: Package deSolve. J Stat Softw. 2010;33: 1–25. doi: 10.18637/jss.v033.i09

22. Turner J, Begon M, Bowers RG, French NP. A model appropriate to the transmission of a human food-borne pathogen in a multigroup managed herd. Prev Vet Med. 2003;57: 175–198. doi: 10.1016/S0167-5877(03)00006-0

23. Robertsson JÅ. Humoral antibody responses to experimental and spontaneous *Salmonella* infections in cattle measured by ELISA. Zentralblatt für Veterinärmedizin Reihe B. 1984;31: 367–380. doi: 10.1111/j.1439-0450.1984.tb01314.x

24. Sojka WJ, Thomson PD, Hudson EB. Excretion of *Salmonella* dublin by adult bovine carriers. Br Vet J. 1974;130: 482–488. doi: 10.1016/S0007-1935(17)35791-3

25. House JK, Smith BP, Dilling GW, Roden LD. Enzyme-linked immunosorbent assay for serologic detection of *Salmonella* dublin carriers on a large dairy. Am J Vet Res. 1993;54: 1391–1399.

26. Smith BP, Oliver DG, Singh P, Dilling G, Martin PA, Ram BP, et al. Detection of *Salmonella* dublin mammary gland infection in carrier cows, using an enzyme-linked immunosorbent assay for antibody in milk or serum. Am J Vet Res. 1989;50: 1352–1360.

27. Nielsen TD, Kudahl AB, Østergaard S, Nielsen LR. Gross margin losses due to *Salmonella* Dublin infection in Danish dairy cattle herds estimated by simulation modelling. Prev Vet Med. 2013;111: 51–62. doi: 10.1016/j.prevetmed.2013.03.011

28. Carrique-Mas JJ, Willmington JA, Papadopoulou C, Watson EN, Davies RH. *Salmonella* infection in cattle in Great Britain, 2003 to 2008. Vet Rec. 2010;167: 560–565. doi: 10.1136/vr.c4943

29. Nennich TD, Harrison JH, Vanwieringen LM, Meyer D, Heinrichs AJ, Weiss WP, et al. Prediction of manure and nutrient excretion from dairy cattle. J Dairy Sci. 2005; 88: 3721–3733. doi: 10.3168/jds.S0022-0302(05)73058-7

30. Nennich TD, Harrison JH, VanWieringen LM, St-Pierre NR, Kincaid RL, Wattiaux MA, et al. Prediction and evaluation of urine and urinary nitrogen and mineral excretion from dairy cattle. J Dairy Sci. 2006;89: 353–364. doi: 10.3168/jds.S0022-0302(06)72101-4

31. Hall GA, Jones PW, Aitken MM. The pathogenesis of experimental intra-ruminal infections of cows with *Salmonella* dublin. J Comp Pathol. 1978;88: 409–417. doi: 10.1016/0021-9975(78)90045-2

32. Hall GA, Jones PW. Experimental oral infections of pregnant heifers with *Salmonella* dublin. Br Vet J. 1979;135: 75–82. doi: 10.1016/S0007-1935(17)32991-3

33. García R, Baelum J, Fredslund L, Santorum P, Jacobsen CS. Influence of temperature and predation on survival of *Salmonella enterica* serovar Typhimurium and expression of invA in soil and manure-amended soil. Appl Environ Microbiol. 2010;76: 5025–5031. doi: 10.1128/AEM.00628-10

34. Natvig EE, Ingham SC, Ingham BH, Cooperband LR, Roper TR. *Salmonella enterica* serovar Typhimurium and *Escherichia coli* contamination of root and leaf vegetables grown in soils with incorporated bovine manure. Appl Env Microbiol, 2002;68: 2737–2744. doi: 10.1128/AEM.68.6.2737-2744.2002

35. Gautam R, Lahodny G, Bani-Yaghoub M, Morley PS, Ivanek R. Understanding the role of cleaning in the control of *Salmonella* Typhimurium in grower-finisher pigs: a modelling approach. Epidemiol Infect. 2014;142: 1034–1049. doi: 10.1017/S0950268813001805

36. Fox BC, Roof MB, Carter DP, Kesl LD, Roth JA. Safety and efficacy of an avirulent live *Salmonella* choleraesuis vaccine for protection of calves against *S* dublin infection. Am J Vet Res. 1997;58: 265–271.

37. House JK, Ontiveros MM, Blackmer NM, Dueger EL, Fitchhorn JB, McArthur GR, et al. Evaluation of an autogenous *Salmonella* bacterin and a modified live *Salmonella* serotype Choleraesuis vaccine on a commercial dairy farm. Am J Vet Res. 2001;62: 1897–1902.

38. Heider LC, Meiring RW, Hoet AE, Gebreyes WA, Funk JA, Wittum TE. Evaluation of vaccination with a commercial subunit vaccine on shedding of *Salmonella* enterica in subclinically infected dairy cows. J Am Vet Med. 2008;233: 466–469. doi: 10.2460/javma.233.3.466

39. Hermesch DR, Thomson DU, Loneragan GH, Renter DR, White BJ. Effects of a commercially available vaccine against *Salmonella enterica* serotype Newport on milk production, somatic cell count, and shedding of *Salmonella* organisms in female dairy cattle with no clinical signs of salmonellosis. Am J Vet Res. 2008;69: 1229–1234. doi: 10.2460/ajvr.69.9.1229

40. Dodd CC, Renter DG, Thomson DU, Nagaraja TG. Evaluation of the effects of a commercially available *Salmonella* Newport siderophore receptor and porin protein vaccine on fecal shedding of *Salmonella* bacteria and health and performance of feedlot cattle. Am J Vet Res. 2011;72: 239–247. doi:10.2460/ajvr.72.2.239

41. Habing GG, Neuder LM, Raphael W, Piper-Youngs H, Kaneene JB. Efficacy of oral administration of a modified-live *Salmonella* Dublin vaccine in calves. J Am Vet Med Assoc. 2011;238: 1184–1190. doi:10.2460/javma.238.9.1184

42. Cummings KJ, Rodriguez-Rivera LD, Capel MB, Rankin SC, Nydam DV. Oral and intranasal administration of a modified-live *Salmonella* Dublin vaccine in dairy calves: clinical efficacy and serologic response. J Dairy Sci. 2019;102: 3474–3479. doi: 10.3168/jds.2018-14892

43. Castro-Vargas RE, Cullens-Nobis FM, Mani R, Roberts JN, Abuelo, A. Effect of dry period immunization of *Salmonella* Dublin latent carriers with a commercial live culture vaccine on intrauterine transmission based on the presence of precolostral antibodies in offspring. J Dairy Sci. 2024;107: 11436–11445. doi: 10.3168/jds.2024-24945

44. Smith BP, Habasha FG, Reina-Guerra M, Hardy AJ. Immunization of calves against salmonellosis. Am J Vet Res. 1980;41: 1947–1951.

45. Robertsson JÅ, Lindberg AA, Hoiseth S, Stocker BA. *Salmonella* typhimurium infection in calves: protection and survival of virulent challenge bacteria after immunization with live or inactivated vaccines. Infect Immun. 1983;41: 742–750. doi: 10.1128/iai.41.2.742-750.1983

46. Smith BP, Reina-Guerra M, Stocker BA, Hoiseth SK, Johnson E. Aromatic-dependent *Salmonella* dublin as a parenteral modified live vaccine for calves. Am J Vet Res. 1984;45: 2231–2235.

47. Smith BP, Reina-Guerra M, Stocker BA, Hoiseth SK, Johnson EH. Vaccination of calves against *Salmonella* dublin with aromatic-dependent Salmonella typhimurium. Am J Vet Res.1984;45: 1858–1861.

48. Jones PW, Dougan G, Hayward C, Mackensie N, Collins P, Chatfield SN. Oral vaccination of calves against experimental salmonellosis using a double aro mutant of *Salmonella* typhimurium. Vaccine. 1991;9: 29–34. doi: 10.1016/0264-410X(91)90313-U

49. Mukkur TK, Walker KH, Jones D, Wronski E, Love DN. Immunizing efficacy of aromatic-dependent *Salmonella* dublin in mice and calves. Comp Immunol Microbiol Infect Dis. 1991;14: 243–256. doi: 10.1016/0147-9571(91)90005-X

50. Smith BP, Dilling GW, Da Roden L, Stocker, BA. Vaccination of calves with orally administered aromatic-dependent *Salmonella* dublin. Am J Vet Res. 1993;54: 1249–1255.

51. Selim SA, Cullor JS, Smith BP, Blanchard P, Farver TB, Hoffman R, et al. The effect of *Escherichia coli* J5 and modified live *Salmonella* dublin vaccines in artificially reared neonatal calves. Vaccine, 1995;13: 381–390. doi: 10.1016/0264-410X(95)98262-9

52. Karszes J, Hill L. Dairy Replacement Program: Cost & Analysis Summer 2019. PRO-DAIRY. 2020 [cited 2022 Mar 20]. Available from: https://dyson.cornell.edu/wp-content/uploads/sites/5/2020/09/Dairy-Replacement-Costs-Writeup-Final1-VD.pdf

53. University of Minnesota. FINBIN; 2020 [cited 30 December 2022]. Database: FINBIN [Internet]. Available from: http://finbin.umn.edu

54. United State Department of Agriculture (USDA). National Dairy Comprehensive Report – Monthly; 2022 [cited 2022 Dec 30]. Database: USDA Economics, Statistics and Market Information System [Internet]. Available from: https://web.archive.org/web/20230704161745/https://usda.library.cornell.edu/concern/publications/vm40z396s?locale=es

55. Campbell JA. Understanding Beef Carcass Yields and Losses During Processing. PennState Extension. 2022 [cited 2024 Feb 15]. Available from: https://web.archive.org/web/20241104190259/https://extension.psu.edu/understanding-beef-carcass-yields-and-losses-during-processing

56. Jones C, Heinrichs J. Growth Charts for Dairy Heifers. PennState Extension. 2022 [cited 2024 Feb 21]. Available from: https://web.archive.org/web/20230316142506/https://extension.psu.edu/growth-charts-for-dairy-heifers

57. Wathes DC, Pollott GE, Johnson KF, Richardson H, Cooke JS. Heifer fertility and carry over consequences for life time production in dairy and beef cattle. Animal. 2014;8: 91–104. doi: 10.1017/S1751731114000755

58. Marino S, Hogue IB, Ray CJ, Kirschner DE. A methodology for performing global uncertainty and sensitivity analysis in systems biology. J Theor Biol. 2008;254: 178–196. doi: 10.1016/j.jtbi.2008.04.011

59. Winston WL. Simulation Modeling Using @RISK: Updated for Version 4. 1^s^t ed. Massachusetts: Cengage Learning; 2001.

60. Kent E, Okafor C, Caldwell M, Walker T, Whitlock B, Lear A. Control of *Salmonella* Dublin in a bovine dairy herd. J Vet Intern Med. 2021;35: 2075–2080. doi: 10.1111/jvim.16191

61. Stevenson M, Sergeant E, Nunes T, Heuer C, Marshall J, Sanchez J, et al. epiR: Tools for the analysis of epidemiological data. CRAN. 2015 [cited 2022 Aug 25]. Available from: https://cran.r-project.org/web/packages/epiR/index.html

62. De La Fuente A, Bing N, Hoeschele I, Mendes P. Discovery of meaningful associations in genomic data using partial correlation coefficients. Bioinformatics. 2004;20: 3565–3574. doi: 10.1093/bioinformatics/bth445

63. Heads JR, Vos A, Blanton J, Müller T, Chipman R, Pieracci EG, et al. Environmental distribution of certain modified live-virus vaccines with a high safety profile presents a low-risk, high-reward to control zoonotic diseases. Sci Rep. 2019;9: 6783. doi: 10.1038/s41598-019-42714-9

64. Rivera RC, Bilal S, Michael E. The relation between host competence and vector-feeding preference in a multi-host model: Chagas and Cutaneous Leishmaniasis. Math Biosci Eng. 2020;17: 5561–5583. doi: 10.3934/mbe.2020299

65. Kuhn M. Building Predictive Models in R Using the caret Package. J Stat Softw. 2008;28: 1–26. doi: 10.18637/jss.v028.i05

66. Rokach L, Maimon O. Classification trees. In: Rokach L, Maimon O, editors. Data mining and knowledge discovery handbook 2^nd^ edition. New York: Springer; 2010. pp. 149–174.

67. Davis MA, Hancock DD, Besser TE, Daniels JB, Baker KN, Call DR. Antimicrobial resistance in *Salmonella enterica* serovar Dublin isolates from beef and dairy sources. Vet Microbiol. 2007;119: 221–230. doi: 10.1016/j.vetmic.2006.08.028

68. Adhikari B, Besser TE, Gay JM, Fox LK, Davis MA, Cobbold RN, et al. The role of animal movement, including off-farm rearing of heifers, in the interherd transmission of multidrug-resistant *Salmonella*. J Dairy Sci. 2009;92: 4229–4238. doi: 10.3168/jds.2008-1494

69. Edrington TS, Callaway TR, Anderson RC, Nisbet DJ. Prevalence of multidrug-resistant *Salmonella* on commercial dairies utilizing a single heifer raising facility. J Food Prot. 2008;71: 27–34. doi: 10.4315/0362-028x-71.1.27

70. Nielsen LR. Review of pathogenesis and diagnostic methods of immediate relevance for epidemiology and control of *Salmonella* Dublin in cattle. Vet Microbiol. 2013;162: 1–9. doi: 10.1016/j.vetmic.2012.08.003

71. Perry KV, Kelton DF, Dufour S, Miltenburg C, Sedo SU, Renaud DL. Risk factors for *Salmonella* Dublin on dairy farms in Ontario, Canada. J Dairy Sci. 2023;106: 9426–9439. doi: 10.3168/jds.2023-23517

72. Gonggrijp MA, Santman-Berends IMGA, Heuvelink AE, Buter GJ, Van Schaik G, Hage JJ, et al. Prevalence and risk factors for extended-spectrum β-lactamase-and AmpC-producing *Escherichia coli* in dairy farms. J Dairy Sci. 2016; 99:9001–9013. doi:10.3168/jds.2016-11134

73. Peters AR. An estimation of the economic impact of an outbreak of *Salmonella* dublin in a calf rearing unit. Vet Rec. 1985;117: 25–26. doi: 10.1136/vr.117.25-26.667

74. Visser SC, Veling J, Dijkhuizen AA, Huirne RBM. Economic losses due to *Salmonella* dublin in dairy cattle. In: Kristensen AR, editor. Animal Health and Management Economics. Copenhagen: Proc. Dutch/Spanish Symp; 1997. pp. 143–151.

75. Bergevoet RHM, Van Schaik G, Veling J, Backus GBC, Franken P. Economic and epidemiological evaluation of *Salmonella* control in Dutch dairy herds. Prev Vet Med. 2009;89: 1–7. doi: 10.1016/j.prevetmed.2008.12.007

76. Holschbach CL, Peek SF. *Salmonella* in dairy cattle. Vet Clin Food Anim. 2018;34: 133–154. doi: 10.1016/j.cvfa.2017.10.005

77. Foster D, Jacob M, Stowe D, Smith G. Exploratory cohort study to determine if dry cow vaccination with a *Salmonella* Newport bacterin can protect dairy calves against oral *Salmonella* challenge. J Vet Intern Med. 2019;33: 1796–1806. doi: 10.1111/jvim.15529

78. Cramer G, Lissemore KD, Guard CL, Leslie KE, Kelton DF. Herd-level risk factors for seven different foot lesions in Ontario Holstein cattle housed in tie stalls or free stalls. J Dairy Sci. 2009;92: 1404–1411. doi: 10.3168/jds.2008-1134

79. Crossley RE, Bokkers EAM, Browne N, Sugrue K, Kennedy E, Conneely M. Risk factors associated with indicators of dairy cow welfare during the housing period in Irish, spring-calving, hybrid pasture-based systems. Prev Vet Med. 2022;208: 105760. doi: 10.1016/j.prevetmed.2022.105760

80. Michigan State University Extension. What is the buzz around cow longevity?. Michigan State University. 2022 [cited 2024 Dec 22]. Available from: https://web.archive.org/web/20241222185847/https://www.canr.msu.edu/news/what-is-the-buzz-around-cow-longevity

81. Veling J, Barkema HW, Van der Schans J, Van Zijderveld F, Verhoeff J. Herd-level diagnosis for *Salmonella enterica* subsp. *enterica* serovar Dublin infection in bovine dairy herds. Prev Vet Med. 2002;53: 31–42. doi: 10.1016/s0167-5877(01)00276-8

82. Velasquez-Munoz A, Castro-Vargas R, Cullens-Nobis FM, Mani R, Abuelo A. *Salmonella* Dublin in dairy cattle. Front Vet Sci. 2023;10: 1331767. doi: 10.3389/fvets.2023.1331767

83. Steinbach G, Koch H, Meyer H, Klaus C. Influence of prior infection on the dynamics of bacterial counts in calves experimentally infected with *Salmonella* dublin. Vet Microbiol. 1996;48: 199–206. doi: 10.1016/0378-1135(95)00134-4

