## Supplementary material for "Integration of mathematical modeling and economics approaches to evaluate strategies for control of *Salmonella* Dublin in a heifer-raising operation": Table S1

**Table S1. Comparison of predictions obtained from the *Salmonella* Dublin model at the end of a 2-year long simulation period with 100, 500, 1,000, and 5,000 iterations. The number of cattle on the farm (N) is 1,000.**

| Animal state | Number of iterations |  |  |  |  |  |  |  |
| --- | --- | --- | --- | --- | --- | --- | --- | --- |
|  | 100 |  | 500 |  | 1,000 |  | 5,000 |  |
|  | Mean | CoV^a^ | Mean | CoV | Mean | CoV | Mean | CoV |
| Susceptible | 37.8 | 0.6 | 35.6 | 0.5 | 35.7 | 0.5 | 32.8 | 0.5 |
| Infectious^b^ | 10.0 | 2.3 | 11.7 | 1.6 | 11.3 | 1.6 | 11.4 | 1.6 |
| Recovered | 52.2 | 0.4 | 52.7 | 0.4 | 53.0 | 0.35 | 52.8 | 0.35 |

^a^ CoV=Coefficient of variation

^b^Proportion of individuals in the Asymptomatic (*A*), Clinically ill (*I*), and Carrier (*C*) compartments by the end of the 2-year simulation period (initial herd size = 1,000).
