## Supplementary material for "Integration of mathematical modeling and economics approaches to evaluate strategies for control of *Salmonella* Dublin in a heifer-raising operation": Table S2

**Table S2. Summary of scenarios assessed to determine the probability of an outbreak (PoO), epidemiological outcomes (DACA: deaths, abortions, and carriers and asymptomatic infections among raised replacement heifers), and/or the operating income (OI) in a heifer-raising operation (HRO).**

| Label | Description | Scenarios | PoO | DACA | OI |
| --- | --- | --- | --- | --- | --- |
|  | Baseline model scenario | Baseline scenario:   - Clean 1x/week - Summer season (temperature starts at 25°C) - 1,000 heifers - *s*=100 (100-fold reduction in *A* and *C*) - All infectious shed - 95% cleaning effectiveness - One asymptomatic heifer enters the operation at t=0 - *β*=10^-10.0^ - *M*=1.5% - *W*=18%* |  |  |  |
|  |  |  | X | X | X |
| Commercial vaccine and improvements in cleaning | Evaluate the influence of vaccination and improvements in cleaning on epidemiological and economic outcomes | - Vaccination |  |  |  |
|  |  | - Clean 3x/week |  |  |  |
|  |  | - Clean 5x/week |  |  |  |
|  |  | - Clean 7x/week |  |  |  |
|  |  | - Clean 2x/day |  |  |  |
|  |  | - Clean 4x/day |  |  |  |
|  |  | - Clean 6x/day |  |  |  |
|  |  | - Clean 8x/day |  |  |  |
|  |  | - Clean 12x/day |  |  |  |
|  |  | - Bedded pack |  |  |  |
|  |  | - Clean 3x/week + Vacc |  |  |  |
|  |  | - Clean 5x/week + Vacc |  |  |  |
|  |  | - Clean 7x/week + Vacc |  |  |  |
|  |  | - Clean 2x/day + Vacc |  |  |  |
|  |  | - Clean 4x/day + Vacc |  |  |  |
|  |  | - Clean 6x/day + Vacc |  |  |  |
|  |  | - Clean 8x/day + Vacc |  |  |  |
|  |  | - Clean 12x/day + Vacc |  |  |  |
|  |  | - Bedded pack + Vacc |  |  |  |
| Seasonality | Evaluate the relevance of temperature in the occurrence of *S.* Dublin outbreaks | - Fall (temperature starts at 9.5°C) - Winter (temperature starts at -5°C) - Spring (temperature starts at 12°C) | X |  |  |
| Herd size | Evaluate the influence of herd size on epidemiological and economic outcomes | - 500 heifers | X | X |  |
|  |  | - 2,000 heifers |  |  |  |
| Shedding reduction in A and C | Evaluate the influence of shedding reduction by heifers in *A* and *C* compared to *I* on epidemiological outcomes | - *s*=1 (no reduction) | X | X |  |
|  |  | - *s*=10 (10-fold reduction in *A* and *C*) |  |  |  |
|  |  | - *s*=1,000 (1,000-fold reduction in *A* and *C*) |  |  |  |
|  |  | - *s*=10 (10-fold reduction in *A*) |  |  |  |
|  |  | - *s*=100 (100-fold reduction in *A*) |  |  |  |
|  |  | - *s*=1,000 (1,000-fold reduction in *A*) |  |  |  |
|  |  | - *s*=10 (10-fold reduction in *C*) |  |  |  |
|  |  | - *s*=100 (100-fold reduction in *C*) |  |  |  |
|  |  | - *s*=1,000 (1,000-fold reduction in *C*) |  |  |  |
| Yes/No shedding by infectious states | Evaluate the influence of the three infectious states included in the model (*A*, *I*, and *C*) on epidemiological outcomes | - Only *A* and *I* shed | X | X |  |
|  |  | - Only *A* and *C* shed |  |  |  |
|  |  | - Only *I* and *C* shed |  |  |  |
|  |  | - Only *A* sheds |  |  |  |
|  |  | - Only *I* sheds |  |  |  |
|  |  | - Only *C* sheds |  |  |  |
| Cleaning effectiveness at each cleaning instance | Evaluate the influence of the cleaning effectiveness at each cleaning instance on epidemiological outcomes | - 90% cleaning effectiveness | X | X |  |
|  |  | - 99% cleaning effectiveness |  |  |  |
|  |  | - 99.9% cleaning effectiveness |  |  |  |
|  |  | - 99.99% cleaning effectiveness |  |  |  |
| Introduction of asymptomatic and/or carrier calves at t=0 | Evaluate the influence of the number and infectious state (*A* or *C*) of the index case(s) introduced in the HRO at t=0 on epidemiological outcomes | - One carrier | X | X |  |
|  |  | - Five asymptomatic |  |  |  |
|  |  | - Five carriers |  |  |  |
|  |  | - One asymptomatic and one carrier |  |  |  |
|  |  | - Five asymptomatic and five carriers |  |  |  |
| Transmission rate | Evaluate the influence of the indirect transmission parameter (*β*) on epidemiological outcomes | - *β*=10^-9.0^ | X | X |  |
|  |  | - *β*=10^-9.5^ |  |  |  |
|  |  | - *β*=10^-10.5^ |  |  |  |
|  |  | - *β*=10^-11^ |  |  |  |
| Probability of carrier development in A | Evaluate the influence of the probability of developing a carrier state in asymptomatic individuals (*M*) on epidemiological outcomes | - *M*=0.15% | X | X |  |
|  |  | - *M*=15% |  |  |  |
| Probability of carrier development in I | Evaluate the influence of the probability of developing a carrier state in clinically ill individuals (*W*) on epidemiological outcomes | - *W*=8% | X | X |  |
|  |  | - *W*=28% |  |  |  |
|  |  | - *W*=38% |  |  |  |

^*^Baseline scenario
