## Supplementary material for "Integration of mathematical modeling and economics approaches to evaluate strategies for control of *Salmonella* Dublin in a heifer-raising operation": Table S3

**Table S3. Detailed epidemiological findings from the scenario analyses evaluated using the developed *Salmonella* Dublin model.** The comparison was made based on the probability of an outbreak and the number of deaths, abortions, carrier replacement heifers, and asymptomatic replacement heifers by the end of the first and second years into the simulation. This table corresponds to the results shown in Fig 7 in the main text. Scenarios are described in Table S2.

| Scenario | Year | Outcome | Median/Proportion | Min | Q1 | Q3 | Max | % change  from  median/  proportion |
| --- | --- | --- | --- | --- | --- | --- | --- | --- |
| **Baseline** |  |  |  |  |  |  |  |  |
| Clean 1x/week |  | Outbreak probability | 79% (785/1,000 iterations) | **-** | **-** | **-** | **-** | **-** |
|  | 1^st^ year | Deaths | 20 | 0 | 9 | 35 | 92 | - |
|  |  | Abortions | 10 | 0 | 5 | 15 | 30 | - |
|  |  | Carrier replacement heifers | 15 | 0 | 11 | 19 | 39 | - |
|  |  | Asymptomatic replacement heifers | 57 | 0 | 34 | 84 | 178 | - |
|  | 2^nd^ year | Deaths | 20 | 0 | 10 | 32 | 76 | - |
|  |  | Abortions | 7 | 0 | 4 | 10 | 22 | - |
|  |  | Carrier replacement heifers | 22 | 1 | 17 | 28 | 56 | - |
|  |  | Asymptomatic replacement heifers | 35 | 2 | 21 | 55 | 128 | - |
| **Vaccination and improvements in cleaning** |  |  |  |  |  |  |  |  |
| Vaccination |  | Outbreak probability | 76% (758/1,000 iterations) | **-** | **-** | **-** | **-** | -3% |
|  | 1^st^ year | Deaths | 5 | 0 | 2 | 10 | 46 | -75 |
|  |  | Abortions | 10 | 0 | 5 | 15 | 31 | 0 |
|  |  | Carrier replacement heifers | 16 | 0 | 10 | 20 | 39 | 7 |
|  |  | Asymptomatic replacement heifers | 58 | 0 | 33 | 88 | 194 | 2 |
|  | 2^nd^ year | Deaths | 5 | 0 | 2 | 9 | 39 | -75 |
|  |  | Abortions | 6 | 0 | 4 | 10 | 25 | -14 |
|  |  | Carrier replacement heifers | 23 | 2 | 17 | 30 | 58 | 5 |
|  |  | Asymptomatic replacement heifers | 36 | 2 | 22 | 58 | 133 | 3 |
| Clean 3x/week |  | Outbreak probability | 73% (732/1,000 iterations) | **-** | **-** | **-** | **-** | -6% |
|  | 1^st^ year | Deaths | 17 | 0 | 6 | 32 | 90 | -15 |
|  |  | Abortions | 9 | 0 | 5 | 15 | 29 | -10 |
|  |  | Carrier replacement heifers | 14 | 0 | 10 | 19 | 38 | -7 |
|  |  | Asymptomatic replacement heifers | 56 | 0 | 31 | 83 | 181 | -2 |
|  | 2^nd^ year | Deaths | 18 | 0 | 8 | 30 | 75 | -10 |
|  |  | Abortions | 6 | 0 | 4 | 10 | 25 | -14 |
|  |  | Carrier replacement heifers | 21 | 1 | 15 | 27 | 56 | -5 |
|  |  | Asymptomatic replacement heifers | 34 | 2 | 20 | 56 | 150 | -3 |
| Clean 5x/week |  | Outbreak probability | 68% (679/1,000 iterations) | **-** | **-** | **-** | **-** | -11% |
|  | 1^st^ year | Deaths | 15 | 0 | 5 | 29 | 87 | -25 |
|  |  | Abortions | 9 | 0 | 5 | 14 | 29 | -10 |
|  |  | Carrier replacement heifers | 14 | 0 | 8 | 18 | 38 | -7 |
|  |  | Asymptomatic replacement heifers | 55 | 0 | 32 | 82 | 171 | -4 |
|  | 2^nd^ year | Deaths | 15 | 0 | 7 | 28 | 73 | -25 |
|  |  | Abortions | 6 | 0 | 3 | 10 | 22 | -14 |
|  |  | Carrier replacement heifers | 20 | 1 | 15 | 26 | 55 | -9 |
|  |  | Asymptomatic replacement heifers | 33 | 2 | 20 | 55 | 117 | -6 |
| Clean 7x/week |  | Outbreak probability | 64% (643/1,000 iterations) | **-** | **-** | **-** | **-** | -15% |
|  | 1^st^ year | Deaths | 13 | 0 | 3 | 26 | 84 | -35 |
|  |  | Abortions | 8 | 0 | 4 | 14 | 29 | -20 |
|  |  | Carrier replacement heifers | 13 | 0 | 6 | 17 | 37 | -13 |
|  |  | Asymptomatic replacement heifers | 54 | 0 | 29 | 78 | 163 | -5 |
|  | 2^nd^ year | Deaths | 13 | 0 | 6 | 25 | 71 | -35 |
|  |  | Abortions | 6 | 1 | 4 | 10 | 22 | -14 |
|  |  | Carrier replacement heifers | 19 | 1 | 14 | 25 | 54 | -14 |
|  |  | Asymptomatic replacement heifers | 34 | 3 | 19 | 54 | 127 | -3 |
| Clean 2x/day |  | Outbreak probability | 49% (488/1,000 iterations) | **-** | **-** | **-** | **-** | -30% |
|  | 1^st^ year | Deaths | 8 | 0 | 1 | 19 | 68 | -60 |
|  |  | Abortions | 7 | 0 | 3 | 12 | 26 | -30 |
|  |  | Carrier replacement heifers | 10 | 0 | 3 | 14 | 34 | -33 |
|  |  | Asymptomatic replacement heifers | 45 | 0 | 19 | 74 | 143 | -21 |
|  | 2^nd^ year | Deaths | 10 | 0 | 4 | 19 | 59 | -50 |
|  |  | Abortions | 6 | 0 | 3 | 9 | 25 | -14 |
|  |  | Carrier replacement heifers | 17 | 1 | 13 | 23 | 50 | -23 |
|  |  | Asymptomatic replacement heifers | 35 | 2 | 19 | 54 | 138 | 0 |
| Clean 4x/day |  | Outbreak probability | 36% (359/1,000 iterations) | **-** | **-** | **-** | **-** | -43% |
|  | 1^st^ year | Deaths | 6 | 0 | 1 | 16 | 49 | -70 |
|  |  | Abortions | 6 | 0 | 2 | 11 | 25 | -40 |
|  |  | Carrier replacement heifers | 7 | 0 | 2 | 13 | 30 | -53 |
|  |  | Asymptomatic replacement heifers | 46 | 0 | 19 | 70 | 135 | -19 |
|  | 2^nd^ year | Deaths | 9 | 0 | 4 | 16 | 45 | -55 |
|  |  | Abortions | 6 | 0 | 3 | 9 | 23 | -14 |
|  |  | Carrier replacement heifers | 16 | 1 | 12 | 22 | 46 | -27 |
|  |  | Asymptomatic replacement heifers | 35 | 2 | 21 | 55 | 116 | 0 |
| Clean 6x/day |  | Outbreak probability | 31% (308/1,000 iterations) | **-** | **-** | **-** | **-** | -48% |
|  | 1^st^ year | Deaths | 5 | 0 | 0 | 14 | 44 | -75 |
|  |  | Abortions | 5 | 0 | 2 | 10 | 24 | -50 |
|  |  | Carrier replacement heifers | 7 | 0 | 1 | 12 | 29 | -53 |
|  |  | Asymptomatic replacement heifers | 40 | 0 | 14 | 68 | 130 | -30 |
|  | 2^nd^ year | Deaths | 8 | 0 | 4 | 15 | 41 | -60 |
|  |  | Abortions | 6 | 1 | 3 | 9 | 26 | -14 |
|  |  | Carrier replacement heifers | 16 | 1 | 12 | 21 | 44 | -27 |
|  |  | Asymptomatic replacement heifers | 36 | 2 | 23 | 55 | 158 | 3 |
| Clean 8x/day |  | Outbreak probability | 28% (278/1,000 iterations) | **-** | **-** | **-** | **-** | -51% |
|  | 1^st^ year | Deaths | 5 | 0 | 0 | 12 | 42 | -75 |
|  |  | Abortions | 5 | 0 | 1 | 10 | 23 | -50 |
|  |  | Carrier replacement heifers | 6 | 0 | 1 | 11 | 28 | -60 |
|  |  | Asymptomatic replacement heifers | 39 | 0 | 11 | 66 | 127 | -32 |
|  | 2^nd^ year | Deaths | 8 | 0 | 4 | 14 | 38 | -60 |
|  |  | Abortions | 5 | 0 | 3 | 9 | 23 | -29 |
|  |  | Carrier replacement heifers | 16 | 1 | 11 | 20 | 42 | -27 |
|  |  | Asymptomatic replacement heifers | 38 | 2 | 21 | 59 | 167 | 9 |
| Clean 12x/day |  | Outbreak probability | 23% (234/1,000 iterations) | **-** | **-** | **-** | **-** | -56% |
|  | 1^st^ year | Deaths | 4 | 0 | 0 | 10 | 38 | -80 |
|  |  | Abortions | 4 | 0 | 1 | 9 | 22 | -60 |
|  |  | Carrier replacement heifers | 5 | 0 | 0 | 9 | 26 | -67 |
|  |  | Asymptomatic replacement heifers | 37 | 0 | 9 | 64 | 121 | -35 |
|  | 2^nd^ year | Deaths | 7 | 0 | 3 | 14 | 33 | -65 |
|  |  | Abortions | 5 | 0 | 3 | 8 | 19 | -29 |
|  |  | Carrier replacement heifers | 16 | 1 | 11 | 20 | 40 | -27 |
|  |  | Asymptomatic replacement heifers | 33 | 2 | 19 | 54 | 139 | -6 |
| Bedded pack |  | Outbreak probability | 64% (638/1,000 iterations) | **-** | **-** | **-** | **-** | -15% |
|  | 1^st^ year | Deaths | 13 | 0 | 3 | 26 | 83 | -35 |
|  |  | Abortions | 8 | 0 | 4 | 14 | 29 | -20 |
|  |  | Carrier replacement heifers | 13 | 0 | 6 | 17 | 37 | -13 |
|  |  | Asymptomatic replacement heifers | 53 | 0 | 29 | 78 | 163 | -7 |
|  | 2^nd^ year | Deaths | 13 | 0 | 6 | 24 | 70 | -35 |
|  |  | Abortions | 6 | 1 | 4 | 10 | 23 | -14 |
|  |  | Carrier replacement heifers | 19 | 1 | 14 | 25 | 54 | -14 |
|  |  | Asymptomatic replacement heifers | 34 | 3 | 20 | 55 | 131 | -3 |
| Clean 3x/week + Vacc |  | Outbreak probability | 70% (701/1,000 iterations) | **-** | **-** | **-** | **-** | -9% |
|  | 1^st^ year | Deaths | 5 | 0 | 2 | 9 | 45 | -75 |
|  |  | Abortions | 9 | 0 | 4 | 14 | 31 | -10 |
|  |  | Carrier replacement heifers | 15 | 0 | 9 | 19 | 38 | 0 |
|  |  | Asymptomatic replacement heifers | 57 | 0 | 32 | 84 | 178 | 0 |
|  | 2^nd^ year | Deaths | 5 | 0 | 2 | 9 | 38 | -75 |
|  |  | Abortions | 6 | 0 | 3 | 10 | 23 | -14 |
|  |  | Carrier replacement heifers | 22 | 1 | 16 | 28 | 56 | 0 |
|  |  | Asymptomatic replacement heifers | 36 | 1 | 21 | 57 | 141 | 3 |
| Clean 5x/week + Vacc |  | Outbreak probability | 65% (650/1,000 iterations) | **-** | **-** | **-** | **-** | -14% |
|  | 1^st^ year | Deaths | 4 | 0 | 1 | 8 | 44 | -80 |
|  |  | Abortions | 9 | 0 | 4 | 14 | 30 | -10 |
|  |  | Carrier replacement heifers | 14 | 0 | 8 | 18 | 36 | -7 |
|  |  | Asymptomatic replacement heifers | 56 | 0 | 31 | 81 | 176 | -2 |
|  | 2^nd^ year | Deaths | 4 | 0 | 2 | 8 | 38 | -80 |
|  |  | Abortions | 6 | 0 | 3 | 10 | 23 | -14 |
|  |  | Carrier replacement heifers | 21 | 1 | 16 | 27 | 55 | -5 |
|  |  | Asymptomatic replacement heifers | 35 | 2 | 20 | 57 | 134 | 0 |
| Clean 7x/week + Vacc |  | Outbreak probability | 60% (606/1,000 iterations) | **-** | **-** | **-** | **-** | -19% |
|  | 1^st^ year | Deaths | 3 | 0 | 1 | 7 | 42 | -85 |
|  |  | Abortions | 8 | 0 | 4 | 13 | 30 | -20 |
|  |  | Carrier replacement heifers | 13 | 0 | 6 | 18 | 34 | -13 |
|  |  | Asymptomatic replacement heifers | 53 | 0 | 27 | 79 | 176 | -7 |
|  | 2^nd^ year | Deaths | 4 | 0 | 2 | 7 | 37 | -80 |
|  |  | Abortions | 6 | 0 | 3 | 10 | 25 | -14 |
|  |  | Carrier replacement heifers | 20 | 1 | 15 | 26 | 54 | -9 |
|  |  | Asymptomatic replacement heifers | 35 | 1 | 21 | 56 | 125 |  |
| Clean 2x/day + Vacc |  | Outbreak probability | 46% (458/1,000 iterations) | **-** | **-** | **-** | **-** | -33% |
|  | 1^st^ year | Deaths | 2 | 0 | 1 | 5 | 37 | -90 |
|  |  | Abortions | 6 | 0 | 3 | 12 | 28 | -40 |
|  |  | Carrier replacement heifers | 11 | 0 | 3 | 16 | 32 | -27 |
|  |  | Asymptomatic replacement heifers | 47 | 0 | 23 | 74 | 157 | -18 |
|  | 2^nd^ year | Deaths | 3 | 0 | 1 | 5 | 34 | -85 |
|  |  | Abortions | 5 | 0 | 3 | 9 | 23 | -29 |
|  |  | Carrier replacement heifers | 18 | 1 | 14 | 24 | 52 | -18 |
|  |  | Asymptomatic replacement heifers | 33 | 2 | 20 | 55 | 164 | -6 |
| Clean 4x/day + Vacc |  | Outbreak probability | 36% (358/1,000 iterations) | **-** | **-** | **-** | **-** | -43% |
|  | 1^st^ year | Deaths | 1 | 0 | 0 | 4 | 31 | -95 |
|  |  | Abortions | 6 | 0 | 2 | 11 | 26 | -40 |
|  |  | Carrier replacement heifers | 9 | 0 | 1 | 13 | 29 | -40 |
|  |  | Asymptomatic replacement heifers | 44 | 0 | 17 | 74 | 147 | -23 |
|  | 2^nd^ year | Deaths | 2 | 0 | 1 | 4 | 30 | -90 |
|  |  | Abortions | 5 | 0 | 3 | 9 | 23 | -29 |
|  |  | Carrier replacement heifers | 17 | 1 | 12 | 23 | 49 | -23 |
|  |  | Asymptomatic replacement heifers | 36 | 2 | 20 | 53 | 150 | 3 |
| Clean 6x/day + Vacc |  | Outbreak probability | 30% (303/1,000 iterations) | **-** | **-** | **-** | **-** | -49% |
|  | 1^st^ year | Deaths | 1 | 0 | 0 | 3 | 27 | -95 |
|  |  | Abortions | 6 | 0 | 2 | 10 | 25 | -40 |
|  |  | Carrier replacement heifers | 8 | 0 | 1 | 12 | 28 | -47 |
|  |  | Asymptomatic replacement heifers | 47 | 0 | 15 | 72 | 148 | -18 |
|  | 2^nd^ year | Deaths | 2 | 0 | 1 | 4 | 27 | -90 |
|  |  | Abortions | 5 | 1 | 3 | 8 | 24 | -29 |
|  |  | Carrier replacement heifers | 17 | 1 | 12 | 22 | 47 | -23 |
|  |  | Asymptomatic replacement heifers | 37 | 2 | 21 | 55 | 141 | 6 |
| Clean 8x/day + Vacc |  | Outbreak probability | 27% (273/1,000 iterations) | **-** | **-** | **-** | **-** | -52% |
|  | 1^st^ year | Deaths | 1 | 0 | 0 | 3 | 25 | -95 |
|  |  | Abortions | 6 | 0 | 2 | 10 | 24 | -40 |
|  |  | Carrier replacement heifers | 7 | 0 | 1 | 12 | 27 | -53 |
|  |  | Asymptomatic replacement heifers | 46 | 0 | 15 | 72 | 139 | -19 |
|  | 2^nd^ year | Deaths | 2 | 0 | 1 | 4 | 25 | -90 |
|  |  | Abortions | 5 | 1 | 3 | 8 | 24 | -29 |
|  |  | Carrier replacement heifers | 17 | 1 | 12 | 21 | 46 | -23 |
|  |  | Asymptomatic replacement heifers | 35 | 5 | 22 | 54 | 155 | 0 |
| Clean 12x/day + Vacc |  | Outbreak probability | 23% (227/1,000 iterations) | **-** | **-** | **-** | **-** | -56% |
|  | 1^st^ year | Deaths | 1 | 0 | 0 | 2 | 21 | -95 |
|  |  | Abortions | 5 | 0 | 2 | 9 | 23 | -50 |
|  |  | Carrier replacement heifers | 5 | 0 | 1 | 10 | 25 | -67 |
|  |  | Asymptomatic replacement heifers | 44 | 0 | 15 | 71 | 130 | -23 |
|  | 2^nd^ year | Deaths | 2 | 0 | 1 | 3 | 22 | -90 |
|  |  | Abortions | 5 | 1 | 3 | 9 | 26 | -29 |
|  |  | Carrier replacement heifers | 17 | 1 | 12 | 20 | 44 | -23 |
|  |  | Asymptomatic replacement heifers | 40 | 3 | 24 | 58 | 148 | 14 |
| Bedded pack + Vacc |  | Outbreak probability | 60% (601/1,000 iterations) | **-** | **-** | **-** | **-** | -19% |
|  | 1^st^ year | Deaths | 3 | 0 | 1 | 7 | 42 | -85 |
|  |  | Abortions | 8 | 0 | 4 | 13 | 30 | -20 |
|  |  | Carrier replacement heifers | 13 | 0 | 5 | 17 | 34 | -13 |
|  |  | Asymptomatic replacement heifers | 53 | 0 | 26 | 77 | 175 | -7 |
|  | 2^nd^ year | Deaths | 4 | 0 | 2 | 7 | 37 | -80 |
|  |  | Abortions | 6 | 0 | 3 | 10 | 25 | -14 |
|  |  | Carrier replacement heifers | 20 | 1 | 15 | 26 | 54 | -9 |
|  |  | Asymptomatic replacement heifers | 35 | 1 | 21 | 56 | 125 | 0 |
| **Seasonality** |  |  |  |  |  |  |  |  |
| Fall (temperature starts at 9.5°C) |  | Outbreak probability | 79% (785/1,000 iterations) | **-** | **-** | **-** | **-** | 0% |
| Winter (temperature starts at -5°C) |  | Outbreak probability | 79% (789/1,000 iterations) | **-** | **-** | **-** | **-** | 0% |
| Spring (temperature starts at 12°C) |  | Outbreak probability | 79% (788/1,000 iterations) | **-** | **-** | **-** | **-** | 0% |
| **Herd size** |  |  |  |  |  |  |  |  |
| 500 heifers |  | Outbreak probability | 58% (582/1,000 iterations) | **-** | **-** | **-** | **-** | -21% |
|  | 1^st^ year | Deaths | 6 | 0 | 1 | 12 | 39 | -70 |
|  |  | Abortions | 4 | 0 | 2 | 7 | 14 | -60 |
|  |  | Carrier replacement heifers | 6 | 0 | 2 | 8 | 18 | -60 |
|  |  | Asymptomatic replacement heifers | 25 | 0 | 14 | 38 | 74 | -56 |
|  | 2^nd^ year | Deaths | 7 | 0 | 3 | 12 | 33 | -65 |
|  |  | Abortions | 3 | 0 | 2 | 5 | 12 | -57 |
|  |  | Carrier replacement heifers | 9 | 1 | 7 | 12 | 26 | -59 |
|  |  | Asymptomatic replacement heifers | 18 | 1 | 10 | 27 | 67 | -49 |
| 2,000 heifers |  | Outbreak probability | 90% (898/1,000 iterations) | **-** | **-** | **-** | **-** | +11% |
|  | 1^st^ year | Deaths | 59 | 0 | 35 | 84 | 194 | 195 |
|  |  | Abortions | 23 | 0 | 14 | 33 | 62 | 130 |
|  |  | Carrier replacement heifers | 36 | 0 | 28 | 44 | 82 | 140 |
|  |  | Asymptomatic replacement heifers | 128 | 0 | 78 | 183 | 371 | 125 |
|  | 2^nd^ year | Deaths | 54 | 0 | 31 | 76 | 160 | 170 |
|  |  | Abortions | 14 | 1 | 8 | 21 | 43 | 100 |
|  |  | Carrier replacement heifers | 50 | 2 | 37 | 63 | 117 | 127 |
|  |  | Asymptomatic replacement heifers | 71 | 5 | 44 | 109 | 270 | 103 |
| **Shedding reduction in A and C** |  |  |  |  |  |  |  |  |
| *s*=1 (no reduction) |  | Outbreak probability | 98% (979/1,000 iterations) | **-** | **-** | **-** | **-** | +19% |
|  | 1^st^ year | Deaths | 39 | 0 | 29 | 51 | 137 | 95 |
|  |  | Abortions | 14 | 0 | 8 | 19 | 57 | 40 |
|  |  | Carrier replacement heifers | 22 | 0 | 19 | 27 | 60 | 47 |
|  |  | Asymptomatic replacement heifers | 61 | 1 | 36 | 90 | 272 | 7 |
|  | 2^nd^ year | Deaths | 33 | 0 | 24 | 44 | 134 | 65 |
|  |  | Abortions | 8 | 1 | 5 | 12 | 42 | 14 |
|  |  | Carrier replacement heifers | 29 | 3 | 22 | 36 | 95 | 32 |
|  |  | Asymptomatic replacement heifers | 39 | 3 | 24 | 60 | 174 | 11 |
| *s*=10 (10-fold reduction) |  | Outbreak probability | 88% (879/1,000 iterations) | **-** | **-** | **-** | **-** | +9% |
|  | 1^st^ year | Deaths | 26 | 0 | 14 | 39 | 95 | 30 |
|  |  | Abortions | 12 | 0 | 7 | 17 | 31 | 20 |
|  |  | Carrier replacement heifers | 18 | 0 | 14 | 22 | 41 | 20 |
|  |  | Asymptomatic replacement heifers | 65 | 1 | 42 | 92 | 182 | 14 |
|  | 2^nd^ year | Deaths | 24 | 0 | 14 | 35 | 78 | 20 |
|  |  | Abortions | 7 | 0 | 4 | 11 | 22 | 0 |
|  |  | Carrier replacement heifers | 25 | 2 | 19 | 31 | 58 | 14 |
|  |  | Asymptomatic replacement heifers | 37 | 4 | 24 | 57 | 135 | 6 |
| *s*=1,000 (1,000-fold reduction) |  | Outbreak probability | 76% (761/1,000 iterations) | **-** | **-** | **-** | **-** | -3% |
|  | 1^st^ year | Deaths | 19 | 0 | 8 | 34 | 91 | -5 |
|  |  | Abortions | 9 | 0 | 5 | 15 | 29 | -10 |
|  |  | Carrier replacement heifers | 14 | 0 | 9 | 18 | 38 | -7 |
|  |  | Asymptomatic replacement heifers | 55 | 0 | 29 | 82 | 181 | -4 |
|  | 2^nd^ year | Deaths | 19 | 0 | 10 | 32 | 76 | -5 |
|  |  | Abortions | 6 | 0 | 3 | 10 | 23 | -14 |
|  |  | Carrier replacement heifers | 21 | 1 | 16 | 27 | 56 | -5 |
|  |  | Asymptomatic replacement heifers | 34 | 1 | 19 | 55 | 156 | -3 |
| *s*=10 (10-fold reduction in *A*) |  | Outbreak probability | 92% (920/1,000 iterations) | **-** | **-** | **-** | **-** | +13% |
|  | 1^st^ year | Deaths | 29 | 0 | 17 | 42 | 98 | 45 |
|  |  | Abortions | 13 | 0 | 7 | 18 | 33 | 30 |
|  |  | Carrier replacement heifers | 19 | 0 | 15 | 23 | 41 | 27 |
|  |  | Asymptomatic replacement heifers | 68 | 0 | 43 | 95 | 185 | 19 |
|  | 2^nd^ year | Deaths | 27 | 0 | 18 | 38 | 80 | 35 |
|  |  | Abortions | 8 | 1 | 5 | 12 | 25 | 14 |
|  |  | Carrier replacement heifers | 27 | 2 | 21 | 34 | 60 | 23 |
|  |  | Asymptomatic replacement heifers | 39 | 3 | 25 | 60 | 140 | 11 |
| *s*=100 (100-fold reduction in *A*) |  | Outbreak probability | 90% (903/1,000 iterations) | **-** | **-** | **-** | **-** | +11% |
|  | 1^st^ year | Deaths | 25 | 0 | 12 | 38 | 96 | 25 |
|  |  | Abortions | 11 | 0 | 6 | 17 | 32 | 10 |
|  |  | Carrier replacement heifers | 17 | 0 | 12 | 21 | 40 | 13 |
|  |  | Asymptomatic replacement heifers | 60 | 0 | 36 | 87 | 180 | 5 |
|  | 2^nd^ year | Deaths | 26 | 0 | 16 | 37 | 80 | 30 |
|  |  | Abortions | 8 | 1 | 5 | 12 | 24 | 14 |
|  |  | Carrier replacement heifers | 26 | 2 | 20 | 33 | 59 | 18 |
|  |  | Asymptomatic replacement heifers | 40 | 3 | 25 | 61 | 140 | 14 |
| *s*=1,000 (1,000-fold reduction in *A*) |  | Outbreak probability | 90% (896/1,000 iterations) | **-** | **-** | **-** | **-** | +11% |
|  | 1^st^ year | Deaths | 24 | 0 | 11 | 38 | 95 | 20 |
|  |  | Abortions | 11 | 0 | 6 | 17 | 32 | 10 |
|  |  | Carrier replacement heifers | 16 | 0 | 11 | 20 | 40 | 7 |
|  |  | Asymptomatic replacement heifers | 58 | 0 | 35 | 85 | 187 | 2 |
|  | 2^nd^ year | Deaths | 26 | 0 | 16 | 37 | 80 | 30 |
|  |  | Abortions | 8 | 1 | 5 | 12 | 23 | 14 |
|  |  | Carrier replacement heifers | 26 | 3 | 20 | 33 | 59 | 18 |
|  |  | Asymptomatic replacement heifers | 40 | 3 | 25 | 62 | 140 | 14 |
| *s*=10 (10-fold reduction in *C*) |  | Outbreak probability | 97% (971/1,000 iterations) | **-** | **-** | **-** | **-** | +18% |
|  | 1^st^ year | Deaths | 38 | 0 | 27 | 49 | 100 | 90 |
|  |  | Abortions | 14 | 0 | 8 | 19 | 33 | 40 |
|  |  | Carrier replacement heifers | 22 | 0 | 18 | 26 | 42 | 47 |
|  |  | Asymptomatic replacement heifers | 60 | 2 | 34 | 88 | 182 | 5 |
|  | 2^nd^ year | Deaths | 32 | 1 | 23 | 42 | 81 | 60 |
|  |  | Abortions | 8 | 1 | 5 | 12 | 23 | 14 |
|  |  | Carrier replacement heifers | 28 | 4 | 22 | 35 | 60 | 27 |
|  |  | Asymptomatic replacement heifers | 38 | 2 | 24 | 59 | 141 | 9 |
| *s*=100 (100-fold reduction in *C*) |  | Outbreak probability | 97% (971/1,000 iterations) | **-** | **-** | **-** | **-** | +18% |
|  | 1^st^ year | Deaths | 38 | 0 | 27 | 49 | 100 | 90 |
|  |  | Abortions | 14 | 0 | 8 | 19 | 33 | 40 |
|  |  | Carrier replacement heifers | 22 | 0 | 18 | 26 | 42 | 47 |
|  |  | Asymptomatic replacement heifers | 60 | 2 | 34 | 88 | 182 | 5 |
|  | 2^nd^ year | Deaths | 32 | 1 | 23 | 42 | 81 | 60 |
|  |  | Abortions | 8 | 1 | 5 | 12 | 23 | 14 |
|  |  | Carrier replacement heifers | 28 | 4 | 22 | 35 | 60 | 27 |
|  |  | Asymptomatic replacement heifers | 38 | 2 | 24 | 59 | 141 | 9 |
| *s*=1,000 (1,000-fold reduction in *C*) |  | Outbreak probability | 97% (971/1,000 iterations) | **-** | **-** | **-** | **-** | +18% |
|  | 1^st^ year | Deaths | 38 | 0 | 27 | 49 | 100 | 90 |
|  |  | Abortions | 14 | 0 | 8 | 19 | 33 | 40 |
|  |  | Carrier replacement heifers | 22 | 0 | 18 | 26 | 42 | 47 |
|  |  | Asymptomatic replacement heifers | 60 | 2 | 35 | 88 | 182 | 5 |
|  | 2^nd^ year | Deaths | 32 | 0 | 23 | 42 | 81 | 60 |
|  |  | Abortions | 8 | 1 | 5 | 12 | 23 | 14 |
|  |  | Carrier replacement heifers | 28 | 4 | 21 | 35 | 60 | 27 |
|  |  | Asymptomatic replacement heifers | 38 | 3 | 24 | 59 | 141 | 9 |
| **Yes/No shedding by infectious states** |  |  |  |  |  |  |  |  |
| *A* and *I* shed |  | Outbreak probability | 79% (785/1,000 iterations) | **-** | **-** | **-** | **-** | 0% |
|  | 1^st^ year | Deaths | 20 | 0 | 9 | 35 | 92 | 0 |
|  |  | Abortions | 10 | 0 | 6 | 15 | 30 | 0 |
|  |  | Carrier replacement heifers | 15 | 0 | 11 | 20 | 39 | 0 |
|  |  | Asymptomatic replacement heifers | 57 | 0 | 34 | 87 | 172 | 0 |
|  | 2^nd^ year | Deaths | 19 | 0 | 10 | 32 | 76 | -5 |
|  |  | Abortions | 6 | 0 | 3 | 10 | 22 | -14 |
|  |  | Carrier replacement heifers | 21 | 1 | 16 | 27 | 56 | -5 |
|  |  | Asymptomatic replacement heifers | 33 | 0 | 19 | 54 | 128 | -6 |
| *A* and *C* shed |  | Outbreak probability | 0% (4/1,000 iterations) | **-** | **-** | **-** | **-** | -79% |
|  | 1^st^ year | Deaths | 0 | 0 | 0 | 0 | 0 | -100 |
|  |  | Abortions | 0 | 0 | 0 | 1 | 4 | -100 |
|  |  | Carrier replacement heifers | 0 | 0 | 0 | 1 | 1 | -100 |
|  |  | Asymptomatic replacement heifers | 6 | 3 | 3 | 18 | 44 | -89 |
|  | 2^nd^ year | Deaths | 2 | 1 | 1 | 3 | 4 | -90 |
|  |  | Abortions | 5 | 2 | 3 | 9 | 17 | -29 |
|  |  | Carrier replacement heifers | 10 | 3 | 5 | 14 | 14 | -55 |
|  |  | Asymptomatic replacement heifers | 79 | 75 | 76 | 89 | 111 | 126 |
| *I* and *C* shed |  | Outbreak probability | 77% (771/1,000 iterations) | **-** | **-** | **-** | **-** | -2% |
|  | 1^st^ year | Deaths | 20 | 0 | 9 | 34 | 92 | 0 |
|  |  | Abortions | 10 | 0 | 5 | 15 | 30 | 0 |
|  |  | Carrier replacement heifers | 15 | 0 | 11 | 19 | 39 | 0 |
|  |  | Asymptomatic replacement heifers | 57 | 0 | 33 | 86 | 174 | 0 |
|  | 2^nd^ year | Deaths | 19 | 0 | 10 | 32 | 76 | -5 |
|  |  | Abortions | 6 | 0 | 4 | 10 | 23 | -14 |
|  |  | Carrier replacement heifers | 21 | 1 | 16 | 28 | 56 | -5 |
|  |  | Asymptomatic replacement heifers | 34 | 1 | 20 | 54 | 150 | -3 |
| Only *A* sheds |  | Outbreak probability | 0% (4/1,000 iterations) | **-** | **-** | **-** | **-** | -79% |
|  | 1^st^ year | Deaths | 0 | 0 | 0 | 0 | 0 | -100 |
|  |  | Abortions | 0 | 0 | 0 | 1 | 3 | -100 |
|  |  | Carrier replacement heifers | 0 | 0 | 0 | 0 | 1 | -100 |
|  |  | Asymptomatic replacement heifers | 3 | 1 | 1 | 12 | 37 | -95 |
|  | 2^nd^ year | Deaths | 1 | 0 | 0 | 2 | 3 | -95 |
|  |  | Abortions | 3 | 1 | 1 | 8 | 17 | -57 |
|  |  | Carrier replacement heifers | 6 | 1 | 2 | 10 | 13 | -73 |
|  |  | Asymptomatic replacement heifers | 59 | 40 | 46 | 80 | 113 | 69 |
| Only *I* sheds |  | Outbreak probability | 77% (768/1,000 iterations) | **-** | **-** | **-** | **-** | -2% |
|  | 1^st^ year | Deaths | 20 | 0 | 8 | 34 | 92 | 0 |
|  |  | Abortions | 10 | 0 | 5 | 15 | 30 | 0 |
|  |  | Carrier replacement heifers | 15 | 0 | 11 | 19 | 39 | 0 |
|  |  | Asymptomatic replacement heifers | 56 | 0 | 33 | 86 | 174 | -2 |
|  | 2^nd^ year | Deaths | 19 | 0 | 9 | 32 | 76 | -5 |
|  |  | Abortions | 6 | 0 | 3 | 10 | 23 | -14 |
|  |  | Carrier replacement heifers | 21 | 1 | 16 | 27 | 56 | -5 |
|  |  | Asymptomatic replacement heifers | 33 | 1 | 19 | 54 | 150 | -6 |
| Only *C* sheds |  | Outbreak probability | 0% (0/1,000 iterations) | **-** | **-** | **-** | **-** | -79% |
|  | 1^st^ year | Deaths | **-** | **-** | **-** | **-** | **-** | - |
|  |  | Abortions | **-** | **-** | **-** | **-** | **-** | - |
|  |  | Carrier replacement heifers | **-** | **-** | **-** | **-** | **-** | - |
|  |  | Asymptomatic replacement heifers | **-** | **-** | **-** | **-** | **-** | - |
|  | 2^nd^ year | Deaths | **-** | **-** | **-** | **-** | **-** | - |
|  |  | Abortions | **-** | **-** | **-** | **-** | **-** | - |
|  |  | Carrier replacement heifers | **-** | **-** | **-** | **-** | **-** | - |
|  |  | Asymptomatic replacement heifers | **-** | **-** | **-** | **-** | **-** | - |
| **Cleaning effectiveness at each cleaning instance** |  |  |  |  |  |  |  |  |
| 90% cleaning effectiveness |  | Outbreak probability | 79% (789/1,000 iterations) | **-** | **-** | **-** | **-** | 0% |
|  | 1^st^ year | Deaths | 21 | 0 | 9 | 35 | 92 | 5 |
|  |  | Abortions | 10 | 0 | 6 | 15 | 30 | 0 |
|  |  | Carrier replacement heifers | 16 | 0 | 12 | 20 | 39 | 7 |
|  |  | Asymptomatic replacement heifers | 58 | 0 | 35 | 87 | 173 | 2 |
|  | 2^nd^ year | Deaths | 20 | 0 | 10 | 32 | 76 | 0 |
|  |  | Abortions | 7 | 0 | 4 | 10 | 21 | 0 |
|  |  | Carrier replacement heifers | 22 | 1 | 16 | 28 | 56 | 0 |
|  |  | Asymptomatic replacement heifers | 35 | 2 | 21 | 54 | 128 | 0 |
| 99% cleaning effectiveness |  | Outbreak probability | 79% (785/1,000 iterations) | **-** | **-** | **-** | **-** | 0% |
|  | 1^st^ year | Deaths | 20 | 0 | 9 | 35 | 92 | 0 |
|  |  | Abortions | 10 | 0 | 6 | 15 | 30 | 0 |
|  |  | Carrier replacement heifers | 15 | 0 | 12 | 20 | 39 | 0 |
|  |  | Asymptomatic replacement heifers | 57 | 0 | 35 | 87 | 172 | 0 |
|  | 2^nd^ year | Deaths | 20 | 0 | 10 | 32 | 76 | 0 |
|  |  | Abortions | 7 | 0 | 4 | 10 | 22 | 0 |
|  |  | Carrier replacement heifers | 22 | 1 | 16 | 28 | 56 | 0 |
|  |  | Asymptomatic replacement heifers | 35 | 2 | 21 | 54 | 128 | 0 |
| 99.9% cleaning effectiveness |  | Outbreak probability | 79% (785/1,000 iterations) | **-** | **-** | **-** | **-** | 0% |
|  | 1^st^ year | Deaths | 20 | 0 | 9 | 35 | 92 | 0 |
|  |  | Abortions | 10 | 0 | 6 | 15 | 30 | 0 |
|  |  | Carrier replacement heifers | 15 | 0 | 12 | 20 | 39 | 0 |
|  |  | Asymptomatic replacement heifers | 57 | 0 | 35 | 87 | 172 | 0 |
|  | 2^nd^ year | Deaths | 20 | 0 | 10 | 32 | 76 | 0 |
|  |  | Abortions | 7 | 0 | 4 | 10 | 22 | 0 |
|  |  | Carrier replacement heifers | 22 | 1 | 16 | 28 | 56 | 0 |
|  |  | Asymptomatic replacement heifers | 35 | 2 | 21 | 54 | 128 | 0 |
| 99.99% cleaning effectiveness |  | Outbreak probability | 79% (785/1,000 iterations) | **-** | **-** | **-** | **-** | 0% |
|  | 1^st^ year | Deaths | 20 | 0 | 9 | 35 | 92 | 0 |
|  |  | Abortions | 10 | 0 | 6 | 15 | 30 | 0 |
|  |  | Carrier replacement heifers | 15 | 0 | 12 | 20 | 39 | 0 |
|  |  | Asymptomatic replacement heifers | 57 | 0 | 35 | 87 | 172 | 0 |
|  | 2^nd^ year | Deaths | 20 | 0 | 10 | 32 | 76 | 0 |
|  |  | Abortions | 7 | 0 | 4 | 10 | 22 | 0 |
|  |  | Carrier replacement heifers | 22 | 1 | 16 | 28 | 56 | 0 |
|  |  | Asymptomatic replacement heifers | 35 | 2 | 21 | 54 | 128 | 0 |
| **Introduction of asymptomatic and/or carrier calves at t=0** |  |  |  |  |  |  |  |  |
| One carrier |  | Outbreak probability | 80% (800/1,000 iterations) | **-** | **-** | **-** | **-** | +1% |
|  | 1^st^ year | Deaths | 20 | 0 | 9 | 35 | 92 | 0 |
|  |  | Abortions | 10 | 0 | 5 | 15 | 30 | 0 |
|  |  | Carrier replacement heifers | 15 | 0 | 11 | 19 | 39 | 0 |
|  |  | Asymptomatic replacement heifers | 56 | 0 | 33 | 84 | 177 | -2 |
|  | 2^nd^ year | Deaths | 19 | 0 | 10 | 32 | 76 | -5 |
|  |  | Abortions | 6 | 0 | 4 | 10 | 21 | -14 |
|  |  | Carrier replacement heifers | 22 | 2 | 16 | 28 | 57 | 0 |
|  |  | Asymptomatic replacement heifers | 34 | 2 | 20 | 54 | 128 | -3 |
| Five asymptomatic |  | Outbreak probability | 79% (794/1,000 iterations) | **-** | **-** | **-** | **-** | 0% |
|  | 1^st^ year | Deaths | 21 | 0 | 9 | 36 | 93 | 5 |
|  |  | Abortions | 10 | 0 | 6 | 15 | 30 | 0 |
|  |  | Carrier replacement heifers | 16 | 0 | 12 | 20 | 39 | 7 |
|  |  | Asymptomatic replacement heifers | 58 | 0 | 35 | 86 | 180 | 2 |
|  | 2^nd^ year | Deaths | 20 | 0 | 10 | 32 | 76 | 0 |
|  |  | Abortions | 6 | 0 | 4 | 10 | 21 | -14 |
|  |  | Carrier replacement heifers | 22 | 1 | 16 | 28 | 56 | 0 |
|  |  | Asymptomatic replacement heifers | 34 | 2 | 20 | 54 | 128 | -3 |
| Five carriers |  | Outbreak probability | 81% (807/1,000 iterations) | **-** | **-** | **-** | **-** | +2% |
|  | 1^st^ year | Deaths | 21 | 0 | 9 | 35 | 92 | 5 |
|  |  | Abortions | 10 | 0 | 6 | 15 | 30 | 0 |
|  |  | Carrier replacement heifers | 16 | 0 | 12 | 20 | 39 | 7 |
|  |  | Asymptomatic replacement heifers | 58 | 0 | 34 | 86 | 181 | 2 |
|  | 2^nd^ year | Deaths | 19 | 0 | 10 | 32 | 76 | -5 |
|  |  | Abortions | 6 | 0 | 4 | 10 | 21 | -14 |
|  |  | Carrier replacement heifers | 23 | 3 | 18 | 30 | 58 | 5 |
|  |  | Asymptomatic replacement heifers | 33 | 2 | 20 | 54 | 128 | -6 |
| One asymptomatic and one carrier |  | Outbreak probability | 80% (801/1,000 iterations) | **-** | **-** | **-** | **-** | +1% |
|  | 1^st^ year | Deaths | 20 | 0 | 9 | 35 | 92 | 0 |
|  |  | Abortions | 10 | 0 | 6 | 15 | 30 | 0 |
|  |  | Carrier replacement heifers | 15 | 0 | 11 | 19 | 39 | 0 |
|  |  | Asymptomatic replacement heifers | 57 | 0 | 33 | 84 | 175 | 0 |
|  | 2^nd^ year | Deaths | 19 | 0 | 10 | 32 | 76 | -5 |
|  |  | Abortions | 6 | 0 | 4 | 10 | 21 | -14 |
|  |  | Carrier replacement heifers | 22 | 2 | 17 | 28 | 57 | 0 |
|  |  | Asymptomatic replacement heifers | 34 | 2 | 20 | 54 | 128 | -3 |
| Five asymptomatic and five carriers |  | Outbreak probability | 81% (807/1,000 iterations) | **-** | **-** | **-** | **-** | +2% |
|  | 1^st^ year | Deaths | 21 | 0 | 9 | 35 | 92 | 5 |
|  |  | Abortions | 10 | 0 | 6 | 15 | 30 | 0 |
|  |  | Carrier replacement heifers | 16 | 0 | 12 | 20 | 39 | 7 |
|  |  | Asymptomatic replacement heifers | 58 | 0 | 34 | 88 | 181 | 2 |
|  | 2^nd^ year | Deaths | 19 | 0 | 10 | 32 | 76 | -5 |
|  |  | Abortions | 6 | 0 | 4 | 10 | 21 | -14 |
|  |  | Carrier replacement heifers | 23 | 3 | 18 | 30 | 58 | 5 |
|  |  | Asymptomatic replacement heifers | 33 | 2 | 20 | 53 | 127 | -6 |
| **Transmission rate** |  |  |  |  |  |  |  |  |
| *β*=10^-9.0^ |  | Outbreak probability | 98% (983/1,000 iterations) | **-** | **-** | **-** | **-** | +19% |
|  | 1^st^ year | Deaths | 41 | 0 | 30 | 52 | 102 | 105 |
|  |  | Abortions | 13 | 0 | 8 | 19 | 33 | 30 |
|  |  | Carrier replacement heifers | 22 | 0 | 18 | 26 | 43 | 47 |
|  |  | Asymptomatic replacement heifers | 59 | 0 | 33 | 88 | 184 | 4 |
|  | 2^nd^ year | Deaths | 35 | 0 | 26 | 45 | 84 | 75 |
|  |  | Abortions | 8 | 0 | 5 | 11 | 23 | 14 |
|  |  | Carrier replacement heifers | 28 | 3 | 22 | 36 | 60 | 27 |
|  |  | Asymptomatic replacement heifers | 37 | 2 | 23 | 59 | 141 | 6 |
| *β*=10^-9.5^ |  | Outbreak probability | 93% (931/1,000 iterations) | **-** | **-** | **-** | **-** | +14% |
|  | 1^st^ year | Deaths | 34 | 0 | 23 | 47 | 99 | 70 |
|  |  | Abortions | 12 | 0 | 8 | 18 | 32 | 20 |
|  |  | Carrier replacement heifers | 20 | 0 | 16 | 24 | 42 | 33 |
|  |  | Asymptomatic replacement heifers | 65 | 0 | 38 | 93 | 202 | 14 |
|  | 2^nd^ year | Deaths | 30 | 0 | 20 | 41 | 81 | 50 |
|  |  | Abortions | 7 | 0 | 4 | 11 | 22 | 0 |
|  |  | Carrier replacement heifers | 26 | 2 | 20 | 33 | 59 | 18 |
|  |  | Asymptomatic replacement heifers | 36 | 2 | 23 | 57 | 138 | 3 |
| *β*=10^-10.5^ |  | Outbreak probability | 41% (412/1,000 iterations) | **-** | **-** | **-** | **-** | -38% |
|  | 1^st^ year | Deaths | 7 | 0 | 1 | 17 | 57 | -65 |
|  |  | Abortions | 5 | 0 | 2 | 11 | 25 | -50 |
|  |  | Carrier replacement heifers | 6 | 0 | 1 | 13 | 30 | -60 |
|  |  | Asymptomatic replacement heifers | 45 | 0 | 14 | 70 | 137 | -21 |
|  | 2^nd^ year | Deaths | 10 | 0 | 4 | 17 | 53 | -50 |
|  |  | Abortions | 6 | 0 | 3 | 9 | 26 | -14 |
|  |  | Carrier replacement heifers | 16 | 1 | 12 | 22 | 47 | -27 |
|  |  | Asymptomatic replacement heifers | 33 | 1 | 20 | 55 | 143 | -6 |
|  |  | Outbreak probability | 6% (57/1,000 iterations) | **-** | **-** | **-** | **-** | **-73%** |
| *β*=10^-11.0^ | 1^st^ year | Deaths | 0 | 0 | 0 | 1 | 13 | -100 |
|  |  | Abortions | 0 | 0 | 0 | 2 | 13 | -100 |
|  |  | Carrier replacement heifers | 0 | 0 | 0 | 2 | 16 | -100 |
|  |  | Asymptomatic replacement heifers | 3 | 0 | 1 | 32 | 92 | -95 |
|  | 2^nd^ year | Deaths | 4 | 0 | 1 | 8 | 23 | -80 |
|  |  | Abortions | 5 | 0 | 2 | 9 | 18 | -29 |
|  |  | Carrier replacement heifers | 10 | 1 | 3 | 16 | 28 | -55 |
|  |  | Asymptomatic replacement heifers | 39 | 2 | 25 | 61 | 126 | 11 |
| **Probability of carrier development in A** |  |  |  |  |  |  |  |  |
| *M*=0.15% |  | Outbreak probability | 78% (782/1,000 iterations) | **-** | **-** | **-** | **-** | **-1%** |
|  | 1^st^ year | Deaths | 20 | 0 | 9 | 35 | 92 | 0 |
|  |  | Abortions | 10 | 0 | 5 | 15 | 30 | 0 |
|  |  | Carrier replacement heifers | 10 | 0 | 7 | 13 | 30 | -33 |
|  |  | Asymptomatic replacement heifers | 58 | 0 | 34 | 86 | 181 | 2 |
|  | 2^nd^ year | Deaths | 20 | 0 | 10 | 32 | 76 | 0 |
|  |  | Abortions | 7 | 0 | 4 | 10 | 22 | 0 |
|  |  | Carrier replacement heifers | 15 | 1 | 11 | 20 | 43 | -32 |
|  |  | Asymptomatic replacement heifers | 35 | 2 | 21 | 57 | 131 | 0 |
| *M*=15% |  | Outbreak probability | 79% (794/1,000 iterations) | **-** | **-** | **-** | **-** | 0% |
|  | 1^st^ year | Deaths | 20 | 0 | 9 | 34 | 92 | 0 |
|  |  | Abortions | 9 | 0 | 5 | 14 | 26 | -10 |
|  |  | Carrier replacement heifers | 63 | 0 | 47 | 74 | 118 | 320 |
|  |  | Asymptomatic replacement heifers | 49 | 0 | 29 | 72 | 152 | -14 |
|  | 2^nd^ year | Deaths | 20 | 0 | 10 | 32 | 76 | 0 |
|  |  | Abortions | 6 | 0 | 4 | 9 | 19 | -14 |
|  |  | Carrier replacement heifers | 80 | 4 | 64 | 100 | 164 | 264 |
|  |  | Asymptomatic replacement heifers | 28 | 2 | 18 | 44 | 106 | -20 |
| **Probability of carrier development in I** |  |  |  |  |  |  |  |  |
| *W*=8% |  | Outbreak probability | 80% (798/1,000 iterations) | **-** | **-** | **-** | **-** | **+1%** |
|  | 1^st^ year | Deaths | 22 | 0 | 10 | 38 | 99 | 10 |
|  |  | Abortions | 10 | 0 | 6 | 15 | 31 | 0 |
|  |  | Carrier replacement heifers | 11 | 0 | 8 | 13 | 25 | -27 |
|  |  | Asymptomatic replacement heifers | 57 | 0 | 34 | 84 | 179 | 0 |
|  | 2^nd^ year | Deaths | 22 | 0 | 11 | 35 | 82 | 10 |
|  |  | Abortions | 7 | 0 | 4 | 10 | 22 | 0 |
|  |  | Carrier replacement heifers | 15 | 1 | 11 | 19 | 36 | -32 |
|  |  | Asymptomatic replacement heifers | 34 | 1 | 21 | 56 | 130 | -3 |
| *W*=28% |  | Outbreak probability | 77% (773/1,000 iterations) | **-** | **-** | **-** | **-** | **-2%** |
|  | 1^st^ year | Deaths | 19 | 0 | 9 | 33 | 86 | -5 |
|  |  | Abortions | 10 | 0 | 5 | 15 | 29 | 0 |
|  |  | Carrier replacement heifers | 19 | 0 | 14 | 25 | 51 | 27 |
|  |  | Asymptomatic replacement heifers | 56 | 0 | 33 | 84 | 176 | -2 |
|  | 2^nd^ year | Deaths | 18 | 0 | 9 | 30 | 71 | -10 |
|  |  | Abortions | 7 | 0 | 4 | 10 | 22 | 0 |
|  |  | Carrier replacement heifers | 28 | 1 | 21 | 36 | 73 | 27 |
|  |  | Asymptomatic replacement heifers | 34 | 2 | 21 | 55 | 131 | -3 |
| *W*=38% |  | Outbreak probability | 76% (762/1,000 iterations) | **-** | **-** | **-** | **-** | **-3%** |
|  | 1^st^ year | Deaths | 18 | 0 | 8 | 30 | 81 | -10 |
|  |  | Abortions | 10 | 0 | 5 | 15 | 29 | 0 |
|  |  | Carrier replacement heifers | 22 | 0 | 16 | 29 | 61 | 47 |
|  |  | Asymptomatic replacement heifers | 56 | 0 | 34 | 84 | 175 | -2 |
|  | 2^nd^ year | Deaths | 17 | 0 | 8 | 28 | 67 | -15 |
|  |  | Abortions | 6 | 0 | 4 | 10 | 22 | -14 |
|  |  | Carrier replacement heifers | 32 | 3 | 24 | 42 | 87 | 45 |
|  |  | Asymptomatic replacement heifers | 34 | 3 | 21 | 54 | 136 | -3 |
