## Supplementary material for "Integration of mathematical modeling and economics approaches to evaluate strategies for control of *Salmonella* Dublin in a heifer-raising operation": Table S4

**Table S4. Operating income (USD per 100-head) observed in scenarios with different costs required for the implementation of mitigation strategies.** A comparison in terms of the operating income (USD per 100-head) was made between the baseline scenario (cleaning once (1x) per week and no vaccination) and scenarios in which control strategies were implemented. These strategies included cleaning at different frequencies (3x, 5x, and 7x times per week and 2x, 4x, 6x, 8x, and 12 per day) and vaccination reducing *S.* Dublin-related mortality (vaccine effectiveness: median=0.72, 5^th^-95^th^ percentile= 0.48–0.91). Operating income in a “no infection” scenario = USD 39,123 per 100-head per year. This table corresponds to the results shown in Fig 8 in the main text.

| **Strategy** | Median | Min | 1^st^ quartile | 3^rd^ quartile | Max | % Difference from baseline median |
| --- | --- | --- | --- | --- | --- | --- |
| **YEAR 1** |  |  |  |  |  |  |
| Clean 1x/week - No vaccination | 32,827 | 24,730 | 30,650 | 34,875 | 39,115 | - |
| **VACCINATION (USD per 100 doses)** |  |  |  |  |  |  |
| **USD 0** |  |  |  |  |  |  |
| Vaccination | 33,611 | 26,009 | 32,068 | 35,464 | 39,115 | 2.4 |
| **USD 100** |  |  |  |  |  |  |
| Vaccination | 33,211 | 25,609 | 31,668 | 35,065 | 38,715 | 1.2 |
| **USD 300** |  |  |  |  |  |  |
| Vaccination | 32,411 | 24,809 | 30,868 | 34,265 | 37,915 | -1.3 |
| **USD 500** |  |  |  |  |  |  |
| Vaccination | 31,612 | 24,010 | 30,069 | 33,465 | 37,116 | -3.7 |
| **USD 700** |  |  |  |  |  |  |
| Vaccination | 30,812 | 23,210 | 29,269 | 32,666 | 36,316 | -6.1 |
| **USD 900** |  |  |  |  |  |  |
| Vaccination | 30,012 | 22,410 | 28,470 | 31,866 | 35,516 | -8.6 |
| **IMPROVEMENTS IN CLEANING (USD per 100 head per day of one additional scraping)** |  |  |  |  |  |  |
| **No additional cost** |  |  |  |  |  |  |
| Clean 3x/week | 33,231 | 25,011 | 31,185 | 35,392 | 39,111 | 1.2 |
| Clean 5x/week | 33,600 | 25,291 | 31,510 | 35,850 | 39,115 | 2.4 |
| Clean 7x/week | 34,070 | 25,578 | 31,875 | 36,211 | 39,115 | 3.8 |
| Clean 2x/day | 35,098 | 26,731 | 32,801 | 37,246 | 39,115 | 6.9 |
| Clean 4x/day | 35,699 | 28,374 | 33,529 | 37,741 | 39,114 | 8.7 |
| Clean 6x/day | 35,953 | 29,067 | 34,012 | 38,378 | 39,113 | 9.5 |
| Clean 8x/day | 35,963 | 29,686 | 34,338 | 38,706 | 39,114 | 9.6 |
| Clean 12x/day | 36,316 | 30,261 | 34,932 | 38,761 | 39,116 | 10.6 |
| **USD 0.15** |  |  |  |  |  |  |
| Clean 3x/week | 33,195 | 24,975 | 31,148 | 35,356 | 39,075 | 1.1 |
| Clean 5x/week | 33,539 | 25,232 | 31,449 | 35,789 | 39,054 | 2.2 |
| Clean 7x/week | 33,984 | 25,495 | 31,790 | 36,125 | 39,029 | 3.5 |
| Clean 2x/day | 34,927 | 26,564 | 32,630 | 37,074 | 38,943 | 6.4 |
| Clean 4x/day | 35,356 | 28,036 | 33,187 | 37,397 | 38,770 | 7.7 |
| Clean 6x/day | 35,438 | 28,559 | 33,498 | 37,862 | 38,597 | 8.0 |
| Clean 8x/day | 35,276 | 29,008 | 33,653 | 38,018 | 38,426 | 7.5 |
| Clean 12x/day | 35,285 | 29,245 | 33,906 | 37,728 | 38,083 | 7.5 |
| **USD 0.30** |  |  |  |  |  |  |
| Clean 3x/week | 33,158 | 24,940 | 31,112 | 35,319 | 39,038 | 1.0 |
| Clean 5x/week | 33,478 | 25,172 | 31,388 | 35,727 | 38,992 | 2.0 |
| Clean 7x/week | 33,898 | 25,412 | 31,705 | 36,039 | 38,943 | 3.3 |
| Clean 2x/day | 34,755 | 26,396 | 32,460 | 36,902 | 38,771 | 5.9 |
| Clean 4x/day | 35,012 | 27,698 | 32,845 | 37,053 | 38,426 | 6.7 |
| Clean 6x/day | 34,923 | 28,051 | 32,984 | 37,346 | 38,081 | 6.4 |
| Clean 8x/day | 34,590 | 28,329 | 32,968 | 37,329 | 37,738 | 5.4 |
| Clean 12x/day | 34,254 | 28,229 | 32,880 | 36,696 | 37,051 | 4.3 |
| **USD 0.45** |  |  |  |  |  |  |
| Clean 3x/week | 33,121 | 24,905 | 31,076 | 35,282 | 39,001 | 0.9 |
| Clean 5x/week | 33,417 | 25,113 | 31,327 | 35,666 | 38,931 | 1.8 |
| Clean 7x/week | 33,812 | 25,329 | 31,620 | 35,953 | 38,857 | 3.0 |
| Clean 2x/day | 34,584 | 26,228 | 32,291 | 36,730 | 38,599 | 5.4 |
| Clean 4x/day | 34,668 | 27,360 | 32,503 | 36,709 | 38,081 | 5.6 |
| Clean 6x/day | 34,408 | 27,543 | 32,470 | 36,830 | 37,564 | 4.8 |
| Clean 8x/day | 33,904 | 27,651 | 32,283 | 36,641 | 37,050 | 3.3 |
| Clean 12x/day | 33,223 | 27,212 | 31,855 | 35,664 | 36,019 | 1.2 |
| **USD 0.60** |  |  |  |  |  |  |
| Clean 3x/week | 33,085 | 24,869 | 31,039 | 35,245 | 38,964 | 0.8 |
| Clean 5x/week | 33,356 | 25,054 | 31,267 | 35,604 | 38,869 | 1.6 |
| Clean 7x/week | 33,726 | 25,246 | 31,535 | 35,867 | 38,771 | 2.7 |
| Clean 2x/day | 34,413 | 26,061 | 32,120 | 36,558 | 38,427 | 4.8 |
| Clean 4x/day | 34,325 | 27,023 | 32,161 | 36,365 | 37,737 | 4.6 |
| Clean 6x/day | 33,893 | 27,035 | 31,956 | 36,314 | 37,048 | 3.2 |
| Clean 8x/day | 33,218 | 26,972 | 31,598 | 35,953 | 36,362 | 1.2 |
| Clean 12x/day | 32,192 | 26,196 | 30,829 | 34,632 | 34,987 | -1.9 |
| **MULTIPLE STRATEGIES (USD per 100 doses - USD per 100 head per day of one additional scraping)** |  |  |  |  |  |  |
| **No additional cost** |  |  |  |  |  |  |
| Vaccination - Clean 2x/day | 35,525 | 28,111 | 33,880 | 37,311 | 39,116 | 8.2 |
| Vaccination - Clean 6x/day | 36,014 | 29,602 | 34,860 | 38,294 | 39,112 | 9.7 |
| Vaccination - Clean 12x/day | 36,522 | 30,405 | 35,467 | 38,525 | 39,112 | 11.3 |
| **No additional cost - USD 0.15** |  |  |  |  |  |  |
| Vaccination - Clean 2x/day | 35,353 | 27,940 | 33,709 | 37,139 | 38,944 | 7.7 |
| Vaccination - Clean 6x/day | 35,498 | 29,087 | 34,344 | 37,778 | 38,596 | 8.1 |
| Vaccination - Clean 12x/day | 35,490 | 29,375 | 34,435 | 37,493 | 38,080 | 8.1 |
| **No additional cost - USD 0.30** |  |  |  |  |  |  |
| Vaccination - Clean 2x/day | 35,181 | 27,768 | 33,538 | 36,967 | 38,772 | 7.2 |
| Vaccination - Clean 6x/day | 34,983 | 28,573 | 33,829 | 37,262 | 38,080 | 6.6 |
| Vaccination - Clean 12x/day | 34,458 | 28,346 | 33,404 | 36,461 | 37,048 | 5.0 |
| **No additional cost - USD 0.45** |  |  |  |  |  |  |
| Vaccination - Clean 2x/day | 35,009 | 27,597 | 33,366 | 36,795 | 38,600 | 6.6 |
| Vaccination - Clean 6x/day | 34,467 | 28,058 | 33,314 | 36,746 | 37,564 | 5.0 |
| Vaccination - Clean 12x/day | 33,426 | 27,316 | 32,372 | 35,428 | 36,016 | 1.8 |
| **USD 100 - No additional cost** |  |  |  |  |  |  |
| Vaccination - Clean 2x/day | 35,125 | 27,711 | 33,481 | 36,912 | 38,716 | 7.0 |
| Vaccination - Clean 6x/day | 35,615 | 29,202 | 34,460 | 37,894 | 38,713 | 8.5 |
| Vaccination - Clean 12x/day | 36,123 | 30,005 | 35,067 | 38,125 | 38,713 | 10.0 |
| **USD 300 - No additional cost** |  |  |  |  |  |  |
| Vaccination - Clean 2x/day | 34,326 | 26,911 | 32,681 | 36,112 | 37,917 | 4.6 |
| Vaccination - Clean 6x/day | 34,815 | 28,402 | 33,660 | 37,094 | 37,913 | 6.1 |
| Vaccination - Clean 12x/day | 35,323 | 29,206 | 34,268 | 37,326 | 37,913 | 7.6 |
| **USD 500 - No additional cost** |  |  |  |  |  |  |
| Vaccination - Clean 2x/day | 33,526 | 26,112 | 31,881 | 35,312 | 37,117 | 2.1 |
| Vaccination - Clean 6x/day | 34,015 | 27,603 | 32,861 | 36,295 | 37,113 | 3.6 |
| Vaccination - Clean 12x/day | 34,523 | 28,406 | 33,468 | 36,526 | 37,113 | 5.2 |
| **USD 100 - USD 0.15** |  |  |  |  |  |  |
| Vaccination - Clean 2x/day | 34,953 | 27,540 | 33,309 | 36,740 | 38,544 | 6.5 |
| Vaccination - Clean 6x/day | 35,099 | 28,687 | 33,945 | 37,378 | 38,196 | 6.9 |
| Vaccination - Clean 12x/day | 35,091 | 28,976 | 34,036 | 37,093 | 37,680 | 6.9 |
| **USD 300 - USD 0.15** |  |  |  |  |  |  |
| Vaccination - Clean 2x/day | 34,154 | 26,740 | 32,510 | 35,940 | 37,745 | 4.0 |
| Vaccination - Clean 6x/day | 34,299 | 27,888 | 33,145 | 36,578 | 37,397 | 4.5 |
| Vaccination - Clean 12x/day | 34,291 | 28,176 | 33,236 | 36,294 | 36,881 | 4.5 |
| **USD 500 - USD 0.15** |  |  |  |  |  |  |
| Vaccination - Clean 2x/day | 33,354 | 25,941 | 31,710 | 35,140 | 36,945 | 1.6 |
| Vaccination - Clean 6x/day | 33,499 | 27,088 | 32,345 | 35,779 | 36,597 | 2.0 |
| Vaccination - Clean 12x/day | 33,491 | 27,376 | 32,436 | 35,494 | 36,081 | 2.0 |
| **USD 100 - USD 0.30** |  |  |  |  |  |  |
| Vaccination - Clean 2x/day | 34,782 | 27,368 | 33,138 | 36,568 | 38,372 | 6.0 |
| Vaccination - Clean 6x/day | 34,583 | 28,173 | 33,429 | 36,862 | 37,680 | 5.3 |
| Vaccination - Clean 12x/day | 34,059 | 27,946 | 33,004 | 36,061 | 36,648 | 3.8 |
| **USD 300 - USD 0.30** |  |  |  |  |  |  |
| Vaccination - Clean 2x/day | 33,982 | 26,569 | 32,338 | 35,768 | 37,573 | 3.5 |
| Vaccination - Clean 6x/day | 33,783 | 27,373 | 32,630 | 36,062 | 36,881 | 2.9 |
| Vaccination - Clean 12x/day | 33,259 | 27,146 | 32,204 | 35,261 | 35,849 | 1.3 |
| **USD 500 - USD 0.30** |  |  |  |  |  |  |
| Vaccination - Clean 2x/day | 33,182 | 25,769 | 31,539 | 34,968 | 36,773 | 1.1 |
| Vaccination - Clean 6x/day | 32,984 | 26,574 | 31,830 | 35,263 | 36,081 | 0.5 |
| Vaccination - Clean 12x/day | 32,459 | 26,347 | 31,405 | 34,462 | 35,049 | -1.1 |
| **USD 100 - USD 0.45** |  |  |  |  |  |  |
| Vaccination - Clean 2x/day | 34,610 | 27,197 | 32,966 | 36,396 | 38,200 | 5.4 |
| Vaccination - Clean 6x/day | 34,067 | 27,659 | 32,914 | 36,346 | 37,164 | 3.8 |
| Vaccination - Clean 12x/day | 33,027 | 26,916 | 31,972 | 35,029 | 35,616 | 0.6 |
| **USD 300 - USD 0.45** |  |  |  |  |  |  |
| Vaccination - Clean 2x/day | 33,810 | 26,398 | 32,167 | 35,596 | 37,401 | 3.0 |
| Vaccination - Clean 6x/day | 33,267 | 26,859 | 32,114 | 35,546 | 36,365 | 1.3 |
| Vaccination - Clean 12x/day | 32,227 | 26,117 | 31,172 | 34,229 | 34,816 | -1.8 |
| **USD 500 - USD 0.45** |  |  |  |  |  |  |
| Vaccination - Clean 2x/day | 33,010 | 25,598 | 31,367 | 34,796 | 36,601 | 0.6 |
| Vaccination - Clean 6x/day | 32,468 | 26,059 | 31,315 | 34,747 | 35,565 | -1.1 |
| Vaccination - Clean 12x/day | 31,427 | 25,317 | 30,373 | 33,429 | 34,017 | -4.3 |
| **YEAR 2** |  |  |  |  |  |  |
| Clean 1x/week - No vaccination | 33,517 | 25,252 | 31,443 | 35,179 | 38,824 | - |
| **VACCINATION (USD per 100 doses)** |  |  |  |  |  |  |
| **USD 0** |  |  |  |  |  |  |
| Vaccination | 34,653 | 27,817 | 33,354 | 36,004 | 38,748 | 3.4 |
| **USD 100** |  |  |  |  |  |  |
| Vaccination | 34,353 | 27,517 | 33,054 | 35,704 | 38,448 | 2.5 |
| **USD 300** |  |  |  |  |  |  |
| Vaccination | 33,754 | 26,917 | 32,455 | 35,104 | 37,849 | 0.7 |
| **USD 500** |  |  |  |  |  |  |
| Vaccination | 33,154 | 26,318 | 31,855 | 34,504 | 37,249 | -1.1 |
| **USD 700** |  |  |  |  |  |  |
| Vaccination | 32,554 | 25,718 | 31,255 | 33,905 | 36,649 | -2.9 |
| **USD 900** |  |  |  |  |  |  |
| Vaccination | 31,955 | 25,118 | 30,656 | 33,305 | 36,049 | -4.7 |
| **IMPROVEMENTS IN CLEANING (USD per 100 head per day of one additional scraping)** |  |  |  |  |  |  |
| **No additional cost** |  |  |  |  |  |  |
| Clean 3x/week | 33,914 | 25,579 | 31,842 | 35,612 | 38,768 | 1.2 |
| Clean 5x/week | 34,135 | 25,913 | 32,139 | 35,790 | 38,797 | 1.8 |
| Clean 7x/week | 34,397 | 26,265 | 32,518 | 36,063 | 38,841 | 2.6 |
| Clean 2x/day | 35,284 | 27,753 | 33,315 | 36,464 | 38,586 | 5.3 |
| Clean 4x/day | 35,392 | 28,810 | 33,706 | 36,494 | 38,794 | 5.6 |
| Clean 6x/day | 35,323 | 29,346 | 33,977 | 36,510 | 38,679 | 5.4 |
| Clean 8x/day | 35,424 | 29,739 | 34,124 | 36,588 | 38,740 | 5.7 |
| Clean 12x/day | 35,603 | 30,410 | 34,310 | 36,807 | 38,785 | 6.2 |
| **USD 0.15** |  |  |  |  |  |  |
| Clean 3x/week | 33,880 | 25,547 | 31,808 | 35,578 | 38,733 | 1.1 |
| Clean 5x/week | 34,078 | 25,860 | 32,084 | 35,733 | 38,739 | 1.7 |
| Clean 7x/week | 34,319 | 26,191 | 32,439 | 35,982 | 38,760 | 2.4 |
| Clean 2x/day | 35,123 | 27,601 | 33,155 | 36,303 | 38,424 | 4.8 |
| Clean 4x/day | 35,070 | 28,500 | 33,387 | 36,172 | 38,470 | 4.6 |
| Clean 6x/day | 34,841 | 28,879 | 33,498 | 36,028 | 38,193 | 4.0 |
| Clean 8x/day | 34,782 | 29,115 | 33,486 | 35,945 | 38,093 | 3.8 |
| Clean 12x/day | 34,633 | 29,468 | 33,351 | 35,838 | 37,814 | 3.3 |
| **USD 0.30** |  |  |  |  |  |  |
| Clean 3x/week | 33,846 | 25,515 | 31,774 | 35,544 | 38,699 | 1.0 |
| Clean 5x/week | 34,021 | 25,807 | 32,028 | 35,676 | 38,681 | 1.5 |
| Clean 7x/week | 34,241 | 26,116 | 32,360 | 35,901 | 38,679 | 2.2 |
| Clean 2x/day | 34,963 | 27,448 | 32,997 | 36,142 | 38,262 | 4.3 |
| Clean 4x/day | 34,749 | 28,191 | 33,069 | 35,850 | 38,146 | 3.7 |
| Clean 6x/day | 34,360 | 28,413 | 33,021 | 35,545 | 37,708 | 2.5 |
| Clean 8x/day | 34,140 | 28,491 | 32,848 | 35,302 | 37,445 | 1.9 |
| Clean 12x/day | 33,664 | 28,526 | 32,391 | 34,869 | 36,843 | 0.4 |
| **USD 0.45** |  |  |  |  |  |  |
| Clean 3x/week | 33,812 | 25,483 | 31,741 | 35,510 | 38,664 | 0.9 |
| Clean 5x/week | 33,964 | 25,754 | 31,972 | 35,619 | 38,623 | 1.3 |
| Clean 7x/week | 34,163 | 26,041 | 32,282 | 35,820 | 38,598 | 1.9 |
| Clean 2x/day | 34,803 | 27,296 | 32,837 | 35,980 | 38,100 | 3.8 |
| Clean 4x/day | 34,428 | 27,882 | 32,751 | 35,528 | 37,822 | 2.7 |
| Clean 6x/day | 33,878 | 27,946 | 32,543 | 35,063 | 37,222 | 1.1 |
| Clean 8x/day | 33,498 | 27,867 | 32,209 | 34,658 | 36,797 | -0.1 |
| Clean 12x/day | 32,694 | 27,585 | 31,432 | 33,900 | 35,871 | -2.5 |
| **USD 0.60** |  |  |  |  |  |  |
| Clean 3x/week | 33,778 | 25,452 | 31,707 | 35,475 | 38,629 | 0.8 |
| Clean 5x/week | 33,907 | 25,700 | 31,917 | 35,562 | 38,566 | 1.2 |
| Clean 7x/week | 34,085 | 25,966 | 32,202 | 35,739 | 38,518 | 1.7 |
| Clean 2x/day | 34,642 | 27,144 | 32,677 | 35,818 | 37,938 | 3.4 |
| Clean 4x/day | 34,107 | 27,573 | 32,432 | 35,206 | 37,499 | 1.8 |
| Clean 6x/day | 33,396 | 27,480 | 32,066 | 34,580 | 36,737 | -0.4 |
| Clean 8x/day | 32,856 | 27,243 | 31,569 | 34,015 | 36,150 | -2.0 |
| Clean 12x/day | 31,725 | 26,643 | 30,471 | 32,931 | 34,900 | -5.3 |
| **MULTIPLE STRATEGIES (USD per 100 doses - USD per 100 head per day of one additional scraping)** |  |  |  |  |  |  |
| **No additional cost** |  |  |  |  |  |  |
| Vaccination - Clean 2x/day | 35,845 | 29,439 | 34,607 | 36,851 | 38,854 | 6.9 |
| Vaccination - Clean 6x/day | 36,231 | 30,881 | 35,209 | 36,890 | 38,576 | 8.1 |
| Vaccination - Clean 12x/day | 36,222 | 31,843 | 35,318 | 36,930 | 38,791 | 8.1 |
| **No additional cost - USD 0.15** |  |  |  |  |  |  |
| Vaccination - Clean 2x/day | 35,683 | 29,283 | 34,446 | 36,689 | 38,693 | 6.5 |
| Vaccination - Clean 6x/day | 35,747 | 30,408 | 34,726 | 36,405 | 38,090 | 6.7 |
| Vaccination - Clean 12x/day | 35,252 | 30,890 | 34,349 | 35,961 | 37,820 | 5.2 |
| **No additional cost - USD 0.30** |  |  |  |  |  |  |
| Vaccination - Clean 2x/day | 35,522 | 29,127 | 34,285 | 36,528 | 38,531 | 6.0 |
| Vaccination - Clean 6x/day | 35,262 | 29,936 | 34,243 | 35,919 | 37,605 | 5.2 |
| Vaccination - Clean 12x/day | 34,281 | 29,937 | 33,379 | 34,992 | 36,848 | 2.3 |
| **No additional cost - USD 0.45** |  |  |  |  |  |  |
| Vaccination - Clean 2x/day | 35,360 | 28,971 | 34,125 | 36,366 | 38,369 | 5.5 |
| Vaccination - Clean 6x/day | 34,777 | 29,464 | 33,759 | 35,434 | 37,119 | 3.8 |
| Vaccination - Clean 12x/day | 33,311 | 28,984 | 32,409 | 34,022 | 35,877 | -0.6 |
| **USD 100 - No additional cost** |  |  |  |  |  |  |
| Vaccination - Clean 2x/day | 35,545 | 29,139 | 34,307 | 36,551 | 38,555 | 6.1 |
| Vaccination - Clean 6x/day | 35,931 | 30,581 | 34,909 | 36,590 | 38,276 | 7.2 |
| Vaccination - Clean 12x/day | 35,922 | 31,543 | 35,018 | 36,631 | 38,491 | 7.2 |
| **USD 300 - No additional cost** |  |  |  |  |  |  |
| Vaccination - Clean 2x/day | 34,946 | 28,539 | 33,707 | 35,951 | 37,955 | 4.3 |
| Vaccination - Clean 6x/day | 35,331 | 29,981 | 34,309 | 35,990 | 37,677 | 5.4 |
| Vaccination - Clean 12x/day | 35,323 | 30,944 | 34,418 | 36,031 | 37,892 | 5.4 |
| **USD 500 - No additional cost** |  |  |  |  |  |  |
| Vaccination - Clean 2x/day | 34,346 | 27,940 | 33,107 | 35,352 | 37,355 | 2.5 |
| Vaccination - Clean 6x/day | 34,732 | 29,381 | 33,710 | 35,391 | 37,077 | 3.6 |
| Vaccination - Clean 12x/day | 34,723 | 30,344 | 33,818 | 35,431 | 37,292 | 3.6 |
| **USD 100 - USD 0.15** |  |  |  |  |  |  |
| Vaccination - Clean 2x/day | 35,384 | 28,983 | 34,146 | 36,390 | 38,393 | 5.6 |
| Vaccination - Clean 6x/day | 35,447 | 30,108 | 34,426 | 36,105 | 37,791 | 5.8 |
| Vaccination - Clean 12x/day | 34,952 | 30,590 | 34,049 | 35,661 | 37,520 | 4.3 |
| **USD 300 - USD 0.15** |  |  |  |  |  |  |
| Vaccination - Clean 2x/day | 34,784 | 28,384 | 33,546 | 35,790 | 37,793 | 3.8 |
| Vaccination - Clean 6x/day | 34,848 | 29,509 | 33,826 | 35,505 | 37,191 | 4.0 |
| Vaccination - Clean 12x/day | 34,352 | 29,991 | 33,449 | 35,061 | 36,920 | 2.5 |
| **USD 500 - USD 0.15** |  |  |  |  |  |  |
| Vaccination - Clean 2x/day | 34,184 | 27,784 | 32,947 | 35,190 | 37,193 | 2.0 |
| Vaccination - Clean 6x/day | 34,248 | 28,909 | 33,226 | 34,905 | 36,591 | 2.2 |
| Vaccination - Clean 12x/day | 33,753 | 29,391 | 32,849 | 34,462 | 36,321 | 0.7 |
| **USD 100 - USD 0.30** |  |  |  |  |  |  |
| Vaccination - Clean 2x/day | 35,222 | 28,827 | 33,985 | 36,228 | 38,231 | 5.1 |
| Vaccination - Clean 6x/day | 34,962 | 29,636 | 33,943 | 35,619 | 37,305 | 4.3 |
| Vaccination - Clean 12x/day | 33,982 | 29,637 | 33,079 | 34,692 | 36,549 | 1.4 |
| **USD 300 - USD 0.30** |  |  |  |  |  |  |
| Vaccination - Clean 2x/day | 34,622 | 28,228 | 33,386 | 35,628 | 37,631 | 3.3 |
| Vaccination - Clean 6x/day | 34,362 | 29,036 | 33,343 | 35,020 | 36,705 | 2.5 |
| Vaccination - Clean 12x/day | 33,382 | 29,038 | 32,479 | 34,092 | 35,949 | -0.4 |
| **USD 500 - USD 0.30** |  |  |  |  |  |  |
| Vaccination - Clean 2x/day | 34,022 | 27,628 | 32,786 | 35,029 | 37,031 | 1.5 |
| Vaccination - Clean 6x/day | 33,763 | 28,437 | 32,743 | 34,420 | 36,105 | 0.7 |
| Vaccination - Clean 12x/day | 32,782 | 28,438 | 31,879 | 33,492 | 35,349 | -2.2 |
| **USD 100 - USD 0.45** |  |  |  |  |  |  |
| Vaccination - Clean 2x/day | 35,060 | 28,672 | 33,825 | 36,066 | 38,069 | 4.6 |
| Vaccination - Clean 6x/day | 34,477 | 29,164 | 33,460 | 35,134 | 36,819 | 2.9 |
| Vaccination - Clean 12x/day | 33,011 | 28,684 | 32,109 | 33,722 | 35,577 | -1.5 |
| **USD 300 - USD 0.45** |  |  |  |  |  |  |
| Vaccination - Clean 2x/day | 34,460 | 28,072 | 33,225 | 35,467 | 37,469 | 2.8 |
| Vaccination - Clean 6x/day | 33,877 | 28,564 | 32,860 | 34,535 | 36,219 | 1.1 |
| Vaccination - Clean 12x/day | 32,411 | 28,084 | 31,509 | 33,123 | 34,977 | -3.3 |
| **USD 500 - USD 0.45** |  |  |  |  |  |  |
| Vaccination - Clean 2x/day | 33,861 | 27,472 | 32,625 | 34,867 | 36,869 | 1.0 |
| Vaccination - Clean 6x/day | 33,277 | 27,964 | 32,260 | 33,935 | 35,620 | -0.7 |
| Vaccination - Clean 12x/day | 31,812 | 27,485 | 30,909 | 32,523 | 34,378 | -5.1 |

Operating income is expressed as USD/100 head/year.
