## Appendix S1 for "Integration of mathematical modeling and economics approaches to evaluate strategies for control of *Salmonella* Dublin in a heifer-raising operation"

**Supplementary Information**

Sebastian Llanos-Soto^1*^, Martin Wiedmann^2^, Aaron Adalja^3^, Christopher Henry^1^, Paolo Moroni^1,4^, Elisha Frye^1^, Francisco A. Leal Yepes^1^, Renata Ivanek^1^

^1^ Department of Population Medicine and Diagnostic Sciences, Cornell University, Ithaca, New York, United States

^2^ Department of Food Science, Cornell University, Ithaca, New York, United States

^3^ School of Hotel Administration, Cornell University, Ithaca, New York, United States

^4^ Dipartimento di Medicina Veterinaria e Scienze Animali, Università degli Studi di Milano, Lodi, Italy

* Corresponding author

**Table of contents**

| **Section I.** Model assumptions | 3 |
| --- | --- |
| **Section II.** Model equations | 5 |
| **Section III.** Parameter description | 6 |
| **Section IV.** Seasonality | 8 |
| **Section V.** Model validation | 10 |
| **Section VI.** Other findings from the model | 13 |
| **References** | 16 |

We developed a mathematical modeling framework that represents the transmission dynamics of *Salmonella* Dublin in a US heifer-raising operation (HRO). The model was used to determine the effectiveness of cleaning improvements (assessed as increases in the cleaning frequency of the barn’s floor) and vaccination in reducing outbreak probability, *S*. Dublin-induced deaths and abortions during raising, and *S*. Dublin carriers and asymptomatic infections among raised replacement heifers. In addition, we use inputs and outputs from the epidemiological components of the model to explore the influence of *S.* Dublin infection and the implementation of vaccination and cleaning improvements on the operating income through multiple cost scenarios. Model assumptions are presented in Section I, the ordinary differential equations (ODEs) defining the model are described in Section II, some relevant model parameters are explained in Section III, the approach to incorporate seasonality into *S.* Dublin dynamics on the farm is described in Section IV, model validation is described in Section V, and additional findings from the model (not described in the manuscript) are presented in Section VI.

I. **Model assumptions**
1. Modes of shedding *S.* Dublin into the environment other than via feces are comparatively negligible and are ignored here. Therefore, the indirect, fecal-oral transmission was the sole mechanism for the spread of *S.* Dublin from infectious to susceptible individuals.
2. Infectiousness occurs without a latent period (i.e., the time between infection and the start of *S.* Dublin shedding), as this period spans only 16-48 hours in *S.* Dublin-infected cattle (Robertsson 1984, Hall et al. 1978, Hall and Jones 1979).
3. Individuals entering the operation are naïve and fully susceptible to infection with *S.* Dublin except for the index case entering the herd at *t*=0. In reality, it is plausible to have recovered, concurrently asymptomatic or carrier calves within an introduced batch. However, the composition of the incoming batches is unknown and thus this assumption was necessary. This assumption was evaluated in scenario analysis (Table S3).
4. There is no reduced susceptibility to *S.* Dublin infection among susceptible individuals who previously experienced the infection and reverted to the *S* compartment upon the loss of immunity. This assumption contrasts with information from Steinbach et al. (1996), however, reduced susceptibility upon loss of immunity was not accounted for due to limited available data and under the principle of parsimony to improve the tractability and interpretability of the model.
5. *S.* Dublin ingestion by calves and heifers was considered to contribute very little to pathogen depletion from the environment (Gautam et al. 2010). For this reason, we only considered the cleaning rate (*μ*) and *S.* Dublin decay rate (*k*) as the factors influencing pathogen removal from the environment. Similarly, the growth of free-living *S.* Dublin in the environment is negligible based on García et al. (2010) and was therefore disregarded.
6. The decay of *S.* Dublin in cattle feces can be approximated by the decay rate (*k*) of *S.* Typhimurium in manure-amended soil (García et al. 2010) or after a single freeze-thaw cycle (Natvig et al. 2002). Utilizing *k* from other *Salmonella* serotypes was necessary due to the lack of information for *S.* Dublin.
7. Carriers are assumed to shed *S*. Dublin in feces continuously, disregarding the possibility of intermittent shedding (Nielsen 2013). This assumption was made because the rate at which individuals cease and restart shedding the pathogen is unknown. By assuming a constant shedding level, this assumption may lead to temporary overestimation or underestimation of *S.* Dublin cell numbers in the environment.
8. Abortions due to *S.* Dublin are assumed to occur only during late pregnancy (i.e., raising stage 12) as they are rarely reported at earlier pregnancy stages (Hinton 1974, Hall and Jones 1976).
9. If vaccination is implemented, all individuals complete their vaccination schedule as soon as they enter the farm (i.e., the period between the first and second vaccination doses is ignored). This may underestimate the *S.* Dublin-induced mortality among weaned and growing calves at the start of the simulation.
II. **Model equations**

The following system of ODEs describes the dynamics of *S.* Dublin transmission for all age categories in a HRO under a baseline scenario where no mitigation strategies have been implemented (*i* in the equations represents weaned calves, growing heifers, and pregnant heifers, while *j* refers only to weaned and growing heifers; subscript *n* refers to one of the twelve progression stages):

$$\frac{dS_{i}}{dt}=-\beta S_{i}E_{n}-{\kappa\beta S}_{i}\sum_{n=1}^{12} E_{n}+\omega R_{i}$$

$$\frac{dA_{i}}{dt}=-\left( \gamma+m \right)A_{i}+\left( 1-u_{i} \right) {(\beta S_{i}E_{n}+\kappa\beta S}_{i}\sum_{n=1}^{12} E_{n})$$

$$\frac{dI_{i}}{dt}=-\left( \gamma+w+d_{j} \right)I_{i}+u_{i}(\beta S_{i}E_{n}+ {\kappa\beta S}_{i}\sum_{n=1}^{12} E_{n})$$

$$\frac{dC_{i}}{dt}=-{\eta C}_{i}+mA_{i}+wI_{i}$$

$$\frac{dR_{i}}{dt}=-\omega R_{i}+\gamma{(A}_{i}+I_{i})+\eta C_{i}$$

$$\frac{dE_{n}}{dt}=-\left( k+\mu\right)E_{n}+\varepsilon_{n}I_{i}+\lambda_{n}(A_{i}+C_{i})$$

III. **Parameter description**

**Recovery rate from the Asymptomatic (*A*) or Clinically ill (*I*) compartments (*γ*)**

The period *D_A/I_* was modeled with a probability distribution $D$*_A/I_*$\sim Beta-PERT(3;17;65)$, where 3, 17, and 65 represented the minimum, mode, and maximum length of time in the compartment, respectively. The value of 17 (mode) represents an average duration of *S.* Dublin shedding in infected calves, as reported by Robertsson (1984) and Nielsen et al. (2007). The rate *γ* was calculated as 1/*D_A/I_*.

**Recovery rate from the Carrier compartment (*η*)**

We considered an average of 365 days based on the limited information available about the duration of shedding in carriers from Sojka et al. (1974), House et al. (1993), Steinbach et al. (1997), and Nielsen et al. (2004a). The value of *D_C_* followed a Beta-PERT distribution ($Beta-PERT\left( 240;365;1,095 \right)$.The minimum value reflects the unlikely stay in the carrier compartment for less than 240 days (see criterion in Nielsen et al. 2004a). Then, the rate *η* was estimated as 1/*D_C_*.

**Immunity loss rate (*ω*)**

The period in the Recovered (R) compartment was modeled with a probability distribution $D$*_R_*$\sim Truncated Exponential(1/140)$, where 140 days is the average duration based on the limited serological findings in the literature (Smith et al. 1989). The minimum duration for *D_R_* was set at 140 days (supported by Smith et al. 1989). The rate *ω* was calculated as 1/*D_R_*.

**Clinically ill (*I*) shedding rate (*ε*)**

The daily fecal shedding was determined from the daily amount of *S.* Dublin shed per gram of feces by cattle showing mild or severe symptoms (*z*), and the daily amount of feces produced by cattle according to the raising stage (*f_n_*, where *n* represents raising stages R1 through R12). The parameter *z* represents the number of *S*. Dublin shed in feces by an average individual in the *I* compartment and was represented with a probability distribution $z\sim Beta-PERT({10}^{3};{10}^{4};{10}^{5.7})$. Then, the daily amount of *S.* Dublin shed in feces for clinically ill individuals was $\varepsilon_{n}=f_{n}*z$*.*

**Asymptomatic individual and carrier shedding rate (*λ*)**

Available data about *S.* Dublin shedding suggest that symptomatic individuals shed a higher amount compared to those asymptomatic or carriers (Sojka et al. 1974, Hall et al. 1978, Hall and Jones 1979, Nielsen et al. 2012). Based on this, we considered that asymptomatic and carrier individuals (*A* and *C*) shed *s*-fold (in baseline *s*=100) less compared to shedding by clinically ill (*I*) individuals (consideration from Nielsen et al. 2012); thus, $\lambda_{n}=\frac{\varepsilon_{n}}{s}$. The implication of this approach was assessed in scenario analysis (see Scenario analysis section of the manuscript).

**Other parameters**

The rates for carrier status development in asymptomatic and clinically ill individuals were reflected in values of parameters *w*=0.18/*D_A/I_* and *m*=0.015/*D_A/I_*, respectively. Parameters *w* and *m* were adopted from Nielsen et al. (2012).

IV. **Seasonality**

Seasonality was incorporated into the model based on the observations in Carrique-Mas et al. (2010) indicating that *S.* Dublin clinical cases in cattle drop during the winter season. To that effect, data about *Salmonella* Typhimurium survival at 5, 15, and 25°C (García et al. 2010) and survival after a freeze/thaw cycle (freezing at -20°C and thawing at 4°C) in manure-amended soil (Natvig et al. 2002) were used to estimate the daily decay rate (*k*) of *S.* Dublin in the farm environment based on the following linearized first-order kinetic model equation (You et al. 2006):

$$k=-\frac{\ln\left( \frac{W}{W_{0}} \right)}{t}$$

where *W_0_* is the initial cell concentration (cell/g) at a time (*t*)=0 and *W* is the cell concentration after *t* days at a certain temperature *T*. We fitted a regression line to *k* values estimated from data for temperatures 5, 15, and 25°C, to extrapolate *k* under seasonally varying average daily ambient temperatures between 0 and 25°C, while a constant value for *k* representing a freeze-thaw cycle event was used for temperatures at and below 0°C (Fig A1). This temperature range (i.e., -5 to 25°C) mimics the daily average temperatures in the Northeastern US (NOAA 2022). Thus, the following formula was used to calculate temperature-specific *k* values for the model:

$$k\left( T \right)=\left\{ \begin{aligned} -0.009*T - 0.46, if 0<T \leq25 \\ 1.25, if-5\leq T\leq0 \end{aligned} \right., T\in\mathbb{R} |-5<T<25$$

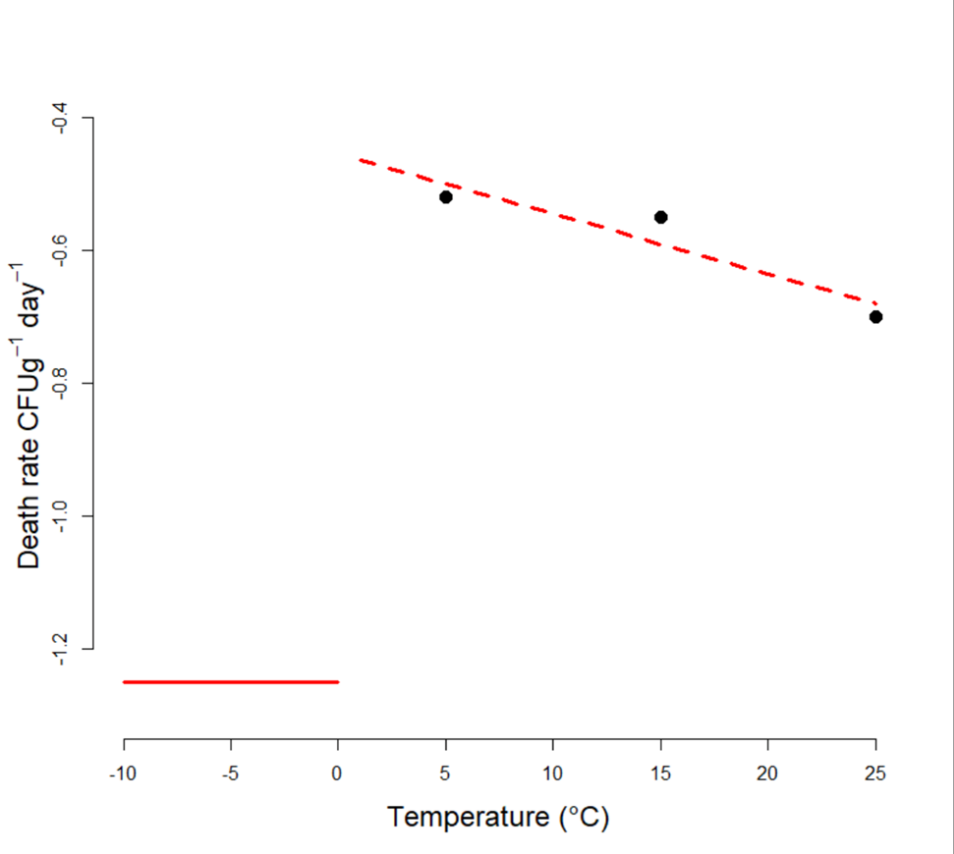

**Fig A1. Temperature-dependent *Salmonella* Dublin decay rate (*k*) in the environment.** Decay rate *(k)* for *S.* Dublin as a function of temperature (*T*) was predicted by a linear function (red broken line) fitted to the data (black circles) from García et al. (2010) and considering the decay of *Salmonella* after a freeze-thaw cycle provided by Natvig et al. (2002) at temperatures ≤ 0°C (red solid line).

V. **Model validation**

No reports on *S*. Dublin spread on HROs were available for model validation. Thus, the model validity was assessed by comparing the predicted seroprevalence of *S.* Dublin-infected animals in a HRO with the reported seroprevalence on a dairy farm in Tennessee (Kent et al., 2021). The validation process consisted of estimating the true seroprevalence at the first serum ELISA testing (conducted in that study one month after *S.* Dublin infection was determined on the farm, but around one month after the last group of animals was introduced into the herd) from the apparent prevalence in Kent et al. (2021) using the formula in Dohoo et al. (2009) considering the sensitivity (*Se*, 65%) and specificities (*Sp*, 97%) for ELISA in Nielsen et al. (2004a). The formula to calculate the true prevalence from apparent prevalence (*AP*) was the following:

$$\frac{AP+Sp-1}{Se+Sp-1}$$

Using this formula, the mean true *S*. Dublin seroprevalence for ELISA in Kent et al. (2021) was 41.9% (95% confidence interval (95% CI): 36.1%, 47.7%). We compared this seroprevalence with the model prediction in terms of the estimated median and the interquartile range (IQR) at the same time point (i.e., around one month since the last batch introduction). For validation, the initial conditions of the model represented the context in Kent et al. (2021), including a herd size of 212 lactating cows (represented by heifers at raising stage 12 in our model), flushing four times per day (assuming that flushing and scraping are equally effective in removing feces), and that the infection likely established during the winter season (between January and February) in Tennessee, coinciding with the arrival of the group of animals from another dairy farm (n=65 cattle). Since the cattle tested with ELISA in Kent et al. (2021) were older than 20 months, we considered the cattle herd in that study to be comparable to the last raising stage in our model (i.e., raising stage 12), which includes pregnant heifers 20 months and older soon to return to their farm of origin to calve. We believe this comparison is fair, as the transmission of the infection between cattle might be similar at that point of maturity. Predictions were generated for various scenarios involving the introduction of pregnant heifers in the batch of 65 individuals in different infectious states, such as single and multiple individuals experiencing infection upon arrival at the farm. Model predictions only considered iterations in which at least three cattle experienced a symptomatic presentation as indicated in Kent et al. (2021). A prediction was considered to approximate the data in Kent et al. (2021) if its interquartile range (IQR) encompassed the true seroprevalence value and its median predicted seroprevalence value was within the 95% CI around the average true seroprevalence estimated considering a sample size of 277 cows (179 from the initial herd, 33 cows that arrived in mid-2015, and 65 cows that arrived in early 2017). The seroprevalence in the model was obtained by adding all *C* and *R* individuals and 60% of the *A* and *I* individuals (representing the proportion of individuals expected to present detectable antibodies one week post infection; Robertsson 1984, Steinbach et al. 1993, Nielsen et al. 2004b). Details about the validation process are provided in Table A1.

**Table A1. The model validation process was done by comparing *Salmonella* Dublin seroprevalence in a Tennessee dairy herd (Kent et al. 2021) and model scenarios in which asymptomatic and/or carrier pregnant heifers were introduced into the herd at t=0 and ELISA serum testing was done 20, 25, 30, 35, and 40 days after the introduction of an infectious cattle.** In Kent et al. (2021), animals were identified to be seropositive to *S.* Dublin around one month after the introduction of multiple groups of cattle into the herd without quarantine. The mean true *S.* Dublin seroprevalence for ELISA in Kent et al. 2021 was estimated to be 41.9% (95% confidence interval (95% CI): 36.1%, 47.7%) considering test sensitivity and specificity of 65% and 97%, respectively (Nielsen et al. 2004a). IQR=interquartile range. Bolded numbers indicate predicted median seroprevalence values that approximate (within the 95% CI of the true seroprevalence of 41.9%) the seroprevalence value observed in Kent et al. (2021)

| Scenario | Predicted prevalence for ELISA at 20 days post introduction (IQR) | Predicted prevalence for ELISA at 25 days post introduction (IQR) | Predicted prevalence for ELISA at 30 days post introduction (IQR) | Predicted prevalence for ELISA at 35 days post introduction (IQR) | Predicted prevalence for ELISA at 40 days post introduction (IQR) | Number of iterations with ≥3 clinical cases |
| --- | --- | --- | --- | --- | --- | --- |
| One asymptomatic | 2.8 (0.9–14.1) | 8.4 (1.8–42.4) | 22.1 (3.7–63.6) | **43.1** (7.6–72.3) | 60.8 (15.1–77.0) | 555/1,000 |
| One carrier | 3.0 (1.1–15.0) | 9.0 (1.9–42.2) | 23.1 (4.1–63.4) | **44.1** (8.1–72.4) | 60.6 (15.8–77.0) | 572/1,000 |
| One carrier one & asymptomatic | 5.0 (1.7–21.8) | 13.4 (3.1–50.7) | 30.1 (6.0–66.6) | 50.4 (11.3–74.0) | 64.2 (20.3–78.3) | 592/1,000 |
| Five asymptomatic | 9.2 (3.5–33.1) | 21.7 (5.8–58.3) | **40.7** (10.2–69.7) | 57.6 (17.5–75.3) | 68.6 (27.9–79.7) | 613/1,000 |
| Five carrier | 10.4 (4.2–34.9) | 23.3 (7.0–59.5) | **42.0** (11.8–70.5) | 58.3 (19.7–75.9) | 68.9 (31.2–80.2) | 626/1,000 |
| Five asymptomatic & five carrier | 15.8 (7.0–43.8) | 31.0 (10.6–63.4) | 49.1 (16.8–72.0) | 62.9 (25.6–77.2) | 71.2 (36.2–81.2) | 640/1,000 |
| Ten asymptomatic | 15.0 (6.4–43.2) | 30.3 (10.0–63.1) | 48.6 (15.7–71.9) | 62.3 (24.9–76.8) | 71.2 (35.9–81.0) | 630/1,000 |
| Ten carrier | 16.6 (7.5–44.8) | 31.9 (11.6–64.1) | 50.1 (17.9–72.6) | 63.7 (26.8–77.5) | 71.4 (37.9–81.6) | 646/1,000 |
| Ten asymptomatic & ten carrier | 23.4 (12.2–51.6) | **39.4** (17.0 –67.4) | 54.6 (24.3–74.2) | 66.4 (33.4–78.6) | 72.9 (43.0–82.2) | 659/1,000 |

VI. **Other findings from the model**

In a scenario with the number of cattle on the farm (N) equal to 2,000, the predicted median deaths, abortions, and carriers and asymptomatic infections among raised replacement heifers all increased compared to the baseline scenario with 1,000 cattle. Herd size was also found to be a predictor of a *S.* Dublin outbreak in a HRO (Table S3). In a HRO of 500 cattle, outbreak probability was lower (58%, 582/1,000 iterations) compared to an operation with 1,000 (79%, 785/1,000 iterations) and 2,000 cattle (90%, 898/1,000 iterations). The median *S.* Dublin-induced deaths, abortions, and carriers and asymptomatic infections among raised replacement heifers in a HRO with 500 cattle were 56% to 70% lower compared to a HRO with 1,000 heifers by the end of year one in the simulation. In contrast, a herd size of 2,000 heifers had values for epidemiological outcomes increased up to 195% compared to a HRO with 1,000 heifers (Table S3).

In the baseline model, we assumed a 100-fold reduction in the shedding rate by asymptomatic and carriers compared to clinically ill individuals. Changing the value of this parameter affected epidemiological outcomes (Table S3). When both carriers and asymptomatic shed *S*. Dublin at the same level as clinically ill individuals, the probability of an outbreak was 98% (979/1,000 iterations), compared to 79% (785/1,000 iterations) at the baseline. Also, compared to the baseline, the median values increased to 39 deaths (IQR=29–51), 14 abortions (IQR=8–19), and 22 carriers (IQR=19–27) and 61 asymptomatic infections (IQR=36–90) among raised replacement heifers by the end of the first year of the simulation. Assuming a 10-fold reduction, the probability of an outbreak increased to 88% (879/1,000 iterations) compared to the baseline. Assuming a 1,000-fold reduction, no substantial changes occurred in epidemiological outcomes compared to baseline. Findings also suggest that reductions in shedding by infectious heifers in either the *A* or *C* compartments have a meaningful impact on *S*. Dublin occurrence and dissemination in an HRO (Table S3). If either *A* or *C* sheds at the same level as *I*, it causes an increase in the probability and consequences of an outbreak.

The importance of the three infectious states (*A*, *I*, and *C*) on model outcomes was assessed in scenarios in which their ability to shed *S*. Dublin into the environment was removed. In these scenarios, instead of the infection being introduced by an asymptomatic individual, the environment was assumed to contain 10^6^ *S*. Dublin cells at t=0. From findings, clinically ill individuals were able to cause an outbreak as sole shedding states, with a 77% (768/1,000 iterations) probability. Meanwhile, carriers were unable to cause and sustain an outbreak without the contribution of acute shedders, while asymptomatic only caused outbreaks as the sole shedders in 4 out of 1,000 iterations. Likewise, the median deaths, abortions, and carriers and asymptomatic infections among raised replacement heifers at the end of the 2-year period were considerably higher when clinically ill were the only shedders (Table S3). For instance, by the end of the first simulated year there were 20 median deaths (IQR=8–34) when clinically ill were the sole shedders, while there were no deaths when asymptomatic were the sole shedders (Table S3).

Introducing multiple infectious index cases into the HRO at time t=0 did not notably increase the risk and consequences of *S.* Dublin infection compared to a single index case introduction. After the introduction of a single or multiple asymptomatic or carrier individuals at time t=0, the likelihood of a *S.* Dublin outbreak occurring when no mitigation strategies were in place was almost identical to the baseline scenario (Table S3). Likewise, only very minor differences were observed in other epidemiological outcomes (Table S3).

Changing the value of the transmission parameter (*β*=10^-10.0^ at baseline) influenced model outcomes (Table S3). The highest *β* value assessed (10^-9.0^*Salmonella* cell^-1^ individual^-1^ day^-1^) led to an outbreak probability of 98% (983/1,000 iterations) and increased the median deaths from a baseline value to 41 (IQR=30–52), abortions to 13 abortions (IQR=8–19), and carriers among raised replacement heifers to 22 (IQR=18–26, Table S3) by the end of the first simulated year. There was no notable impact on asymptomatic infections among raised replacement heifers. For the lowest assessed *β* value (10^-11.0^ *Salmonella* cell^-1^ individual^-1^ day^-1^), the probability of an outbreak was 6% (57/1,000 iterations) and median values for deaths, abortions, and carriers among raised replacement heifers decreased to 0 after the first year into the simulation, with values remaining low by the second year of the simulation (Table S3).

The probability of developing a carrier state in asymptomatic individuals (*M*) and the probability of developing a carrier state in clinically ill individuals (*W*) were at baseline set to 0.015 and 0.18 respectively. As expected, increasing values for these parameters greatly influenced carriers among raised replacement heifers. A value of *M*=15% caused median carriers among raised replacement heifers to increase to 63 (IQR=47–74, Table S3) by the end of year one in the simulation, representing a 320% increase compared to the baseline. Increasing *W* to 28%, the median carriers among raised replacement heifers increased to 19 (IQR=14–25, Table S6) by the end of year one, representing a 27% increase in departures. Neither *M* nor *W* meaningfully influenced the outbreak probability.

When assessing seasonality, the ambient temperature at the time an asymptomatic individual (or a carrier) was introduced into the HRO did not meaningfully influence the probability of an outbreak and estimations for model outcomes (results not shown). Similarly, the cleaning effectiveness in terms of % of feces removed at each cleaning instance (*H*) did not influence outcomes (Table S3).

**References**

1. Robertsson JÅ. Humoral antibody responses to experimental and spontaneous *Salmonella* infections in cattle measured by ELISA*.* Zentralblatt für Veterinärmedizin Reihe B. 1984;31: 367-380. [doi: 10.1111/j.1439-0450.1984.tb01314.x](https://doi.org/10.1111/j.1439-0450.1984.tb01314.x)
2. Hall GA, Jones PW, Aitken MM. The pathogenesis of experimental intra-ruminal infections of cows with *Salmonella* dublin*.* J Comp Pathol. 1978;88: 409-417. [doi: 10.1016/0021-9975(78)90045-2](https://doi.org/10.1016/0021-9975(78)90045-2)
3. Hall GA, Jones PW. Experimental oral infections of pregnant heifers with *Salmonella* dublin. Br Vet J. 1979;135: 75-82. [doi: 10.1016/S0007-1935(17)32991-3](https://doi.org/10.1016/S0007-1935(17)32991-3)
4. Steinbach G, Koch H, Meyer H, Klaus C. Influence of prior infection on the dynamics of bacterial counts in calves experimentally infected with *Salmonella* dublin. Vet Microbiol. 1996;48: 199-206. doi: 10.1016/0378-1135(95)00134-44
5. Gautam R, Lahodny G, Bani-Yaghoub M, Morley PS, Ivanek R. Understanding the role of cleaning in the control of *Salmonella* Typhimurium in grower-finisher pigs: a modelling approach. Epidemiol Infect. 2014;142: 1034-1049. [doi:](https://doi.org/10.1017/S0950268813001805) 10.1017/S0950268813001805
6. García R, Baelum J, Fredslund L, Santorum P, Jacobsen CS. Influence of temperature and predation on survival of *Salmonella enterica* serovar Typhimurium and expression of invA in soil and manure-amended soil. Appl Environ Microbiol. 2010;76: 5025-5031. [doi:](https://doi.org/10.1128/AEM.00628-10) 10.1128/AEM.00628-10
7. Natvig EE, Ingham SC, Ingham BH, Cooperband LR, Roper TR. *Salmonella enterica* serovar Typhimurium and *Escherichia coli* contamination of root and leaf vegetables grown in soils with incorporated bovine manure. Appl Env Microbiol, 2002;68: 2737-2744. doi: 10.1128/AEM.68.6.2737-2744.2002
8. Nielsen LR. Within-herd prevalence of *Salmonella* Dublin in endemically infected dairy herds. Epidemiol Infect. 2013;141: 20742082. doi: 10.1017/S0950268812003007
9. Hinton M. *Salmonella* dublin abortion in cattle: studies on the clinical aspects of the condition. British Veterinary Journal. 1974;130: 556-563. doi: 10.1016/S0007-1935(17)35742-1
10. Hall GA, Jones PW. An experimental study of *Salmonella* dublin abortion in cattle. British Veterinary Journal. 1976;132: 60-65. doi: 10.1016/S0007-1935(17)34788-7
11. Nielsen LR, van den Borne B, van Schaik G. *Salmonella* Dublin infection in young dairy calves: Transmission parameters estimated from field data and an SIR-model. Prev Vet Med. 2007;79: 46-58. doi: [10.1016/j.prevetmed.2006.11.006](https://doi.org/10.1016/j.prevetmed.2006.11.006)
12. Sojka WJ, Thomson PD, Hudson EB. Excretion of *Salmonella* dublin by adult bovine carriers. Br Vet J. 1974;130: 482-488. [doi: 10.1016/S0007-1935(17)35791-3](https://doi.org/10.1016/S0007-1935(17)35791-3)
13. House JK, Smith BP, Dilling GW, Roden LD. Enzyme-linked immunosorbent assay for serologic detection of *Salmonella* dublin carriers on a large dairy. Am J Vet Res. 1993;54: 1391-1399.
14. Steinbach G, Methner U, Koch H, Meyer H. Intercurrent infections as a cause for the development of *Salmonella* carriers. In: Proceedings of the International Symposium *Salmonella* and Salmonellosis; 1997. p 255–260.
15. Nielsen LR, Schukken YH, Gröhn YT, Ersbøll AK. *Salmonella* Dublin infection in dairy cattle: risk factors for becoming a carrier. Prev Vet Med. 2004a;65: 47-62. doi: 10.1016/j.prevetmed.2004.06.010
16. Smith BP, Oliver DG, Singh P, Dilling G, Martin PA, Ram BP, et al. Detection of *Salmonella* dublin mammary gland infection in carrier cows, using an enzyme-linked immunosorbent assay for antibody in milk or serum. Am J Vet Res. 1989;50: 1352–1360.
17. Nielsen LR, Kudahl AB, Østergaard S. Age-structured dynamic, stochastic and mechanistic simulation model of *Salmonella* Dublin infection within dairy herds. Prev Vet Med. 2012;105: 59-74. [doi: 10.1016/j.prevetmed.2012.02.005](https://doi.org/10.1016/j.prevetmed.2012.02.005)
18. Carrique-Mas JJ, Willmington JA, Papadopoulou C, Watson EN, Davies RH. *Salmonella* infection in cattle in Great Britain, 2003 to 2008. Vet Rec. 2010;167: 560-565. doi: 10.1136/vr.c4943
19. You Y, Rankin SC, Aceto HW, Benson CE, Toth JD, Dou Z. Survival of *Salmonella enterica* serovar Newport in manure and manure-amended soils. Appl Environ Microbiol. 2006;72: 5777-5783. doi: 10.1128/AEM.00791-06
20. National Oceanic and Atmospheric Administration (NOAA). Climate Data Online; 2022 [cited 20 November 2022]. Database: CDO [Internet]. Available from: <https://www.ncei.noaa.gov/cdo-web/>
21. Kent E, Okafor C, Caldwell M, Walker T, Whitlock B, Lear A. Control of *Salmonella* Dublin in a bovine dairy herd. J Vet Intern Med. 2021;35: 2075-2080. doi: 10.1111/jvim.16191
22. Dohoo I, Martin W, Stryhn H. Veterinary Epidemiologic Research. 2nd edition ed. Charlottetown, CA: VER Inc.; 2009. 865 p.
23. Steinbach G, Dinjus U, Gottschaldt J, Kreutzer B, Staak C. Course of infection and humoral immune reaction in calves infected orally with different *Salmonella* serovars. Zentralbl Veterinarmed B. 1993;40: 515-521. doi: 10.1111/j.1439-0450.1993.tb00171.x
24. Nielsen LR, Toft N, Ersbøll AK. Evaluation of an indirect serum ELISA and a bacteriological faecal culture test for diagnosis of *Salmonella* serotype Dublin in cattle using latent class models. J Appl Microbiol. 2004b; 96:311-319. doi: 10.1046/j.1365-2672.2004.02151.x
